# Genetic Underpinnings of Risky Behaviour Relate to Altered Neuroanatomy

**DOI:** 10.1101/862417

**Authors:** Gökhan Aydogan, Remi Daviet, Richard Karlsson Linnér, Todd A. Hare, Joseph W. Kable, Henry R. Kranzler, Reagan R. Wetherill, Christian C. Ruff, Philipp D. Koellinger, Gideon Nave

## Abstract

Previous research points to the heritability of risk-taking behaviour. However, evidence on how genetic dispositions are translated into risky behaviour is scarce. Here, we report a genetically-informed neuroimaging study of real-world risky behaviour across the domains of drinking, smoking, driving, and sexual behaviour, in a European sample from the UK Biobank (*N*= 12,675). We find negative associations between risky behaviour and grey matter volume (GMV) in distinct brain regions, including amygdala, ventral striatum, hypothalamus, and dorsolateral prefrontal cortex (dlPFC). These effects replicate in an independent sample recruited from the same population (*N*=13,004). Polygenic risk scores for risky behaviour, derived from a genome-wide association study in an independent sample (*N*=297,025), are inversely associated with GMV in dlPFC, putamen, and hypothalamus. This relation mediates ~2.2% of the association between genes and behaviour. Our results highlight distinct heritable neuroanatomical features as manifestations of the genetic propensity for risk taking.

**One Sentence Summary:** Risky behaviour and its genetic associations are linked to less grey matter volume in distinct brain regions.

## Main

Taking risks—an essential element of many human experiences and achievements—requires balancing uncertain positive and negative outcomes. For instance, exploration, innovation and entrepreneurship can yield great benefits, but are also prone to failure^1^. Conversely, excessive risk-taking in markets can have enormous societal costs, such as speculative price bubbles^2^. Similarly, common behaviours such as *smoking, drinking, sexual promiscuity*, or *speeding* are considered rewarding by many but might expose individuals and those around them to deleterious health, social, and financial consequences. In 2010, the combined economic burden in the United States of these risky behaviours was estimated to be about $593.3 billion^3–6^. Although previous findings point to the partial heritability of risk tolerance and risky behaviours^7^ and neuroanatomical measures exhibit high heritability^8,9^, little is known about the brain features involved in translating genetic dispositions into risky behavioural phenotypes^8^.

Recent research using structural brain-imaging data from small, nonrepresentative samples (comprising up to a few hundred participants) has identified several neuroanatomical associations with risk tolerance^10–12^. However, this literature is limited by low statistical power^13,14^, and the generalizability of their findings to other populations is questionable. Small sample sizes have also limited the ability to control systematically for many factors that could confound observed relations between brain features and risky behaviour, such as height^15^ and genetic population structure^16,17^. Moreover, despite evidence that the effects of genetic factors are likely mediated by their influence on the brain and its development^7,18^, neuroscientific and genetic approaches to understanding the biology of risky behaviour have largely proceeded in isolation–perhaps due to the lack of large study samples that include both genetic and brain imaging measures.

Here, we utilize data obtained in a prospective epidemiological study of ~500,000 individuals aged 40 to 69 years [the UK Biobank (UKB)^19,20^] to carry out a pre-registered investigation (https://osf.io/qkp4g/, see Supplementary Methods for deviations from the analysis plan) of the relationship between individual differences in brain anatomy and the propensity to engage in risky behaviour across four domains (*N*= 12,675). We replicate our findings in an independent sample recruited from the same population (*N*=13,004). Further, we isolate specific differences in brain anatomy that are linked to the genetic disposition for risky behaviour— quantified via polygenic risk scores (PRS) derived from a genome-wide association study (GWAS) in an independent sample (*N* = 297,025)—and investigate how these neuroanatomical endophenotypes mediate the influence of genetics on the behavioural phenotype.

## Results

### Grey Matter Volume Associations with Risky Behaviour

Akin to a previous investigation^7^, we construct a measure of risky behaviour by extracting the first principal component from four self-reported measures of drinking, smoking, speeding on motorways, and sexual promiscuity (*N* = 315,855; see Figure 1A, Supplementary Methods 1.1, Supplementary Figures 1-2 and Supplementary Tables 1-2 for descriptive statistics). This measure of risky behaviour is genetically correlated with many other traits, including cannabis use (*r*_g_ = 0.72, *SE* = 0.02), general risk tolerance (*r*_g_ = 0.56, *SE* = 0.02), self-employment (*r_g_* = 0.52; *SE* = 0.30), suicide attempt (*r*_g_ = 0.47, *SE* = 0.07), antisocial behaviour (*r*_g_ = 0.45, *SE* = 0.14), extraversion (*r*_g_ = 0.34, *SE* = 0.04), and age at first sexual experience (*r*_g_ = −0.54, *SE* = 0.02) (Supplementary Methods 1.5 and Supplementary Table 3). Thus, our measure is partly rooted in genetic differences between people and relates to a broad range of relevant events and behaviours.

**Figure 1.**
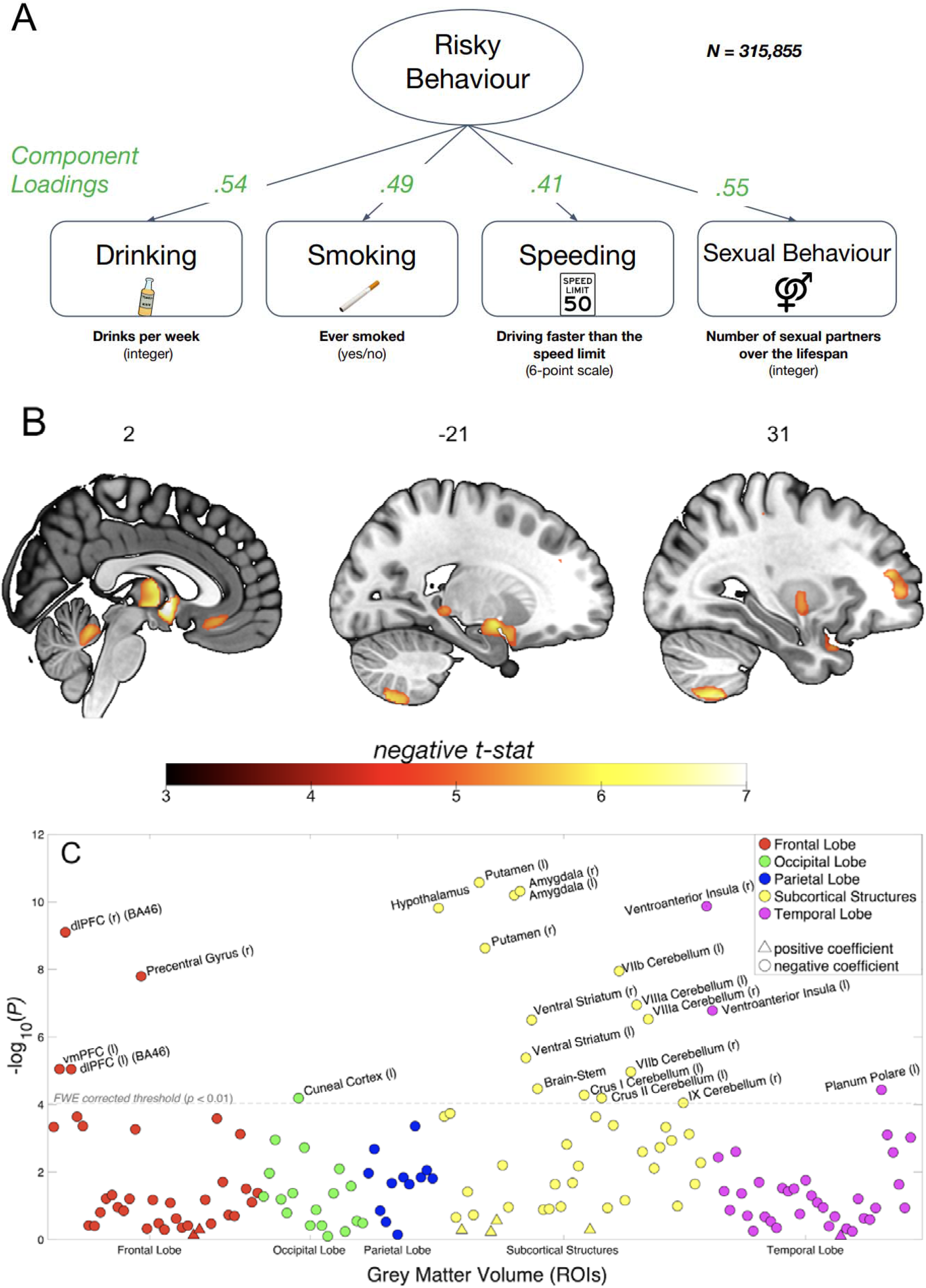
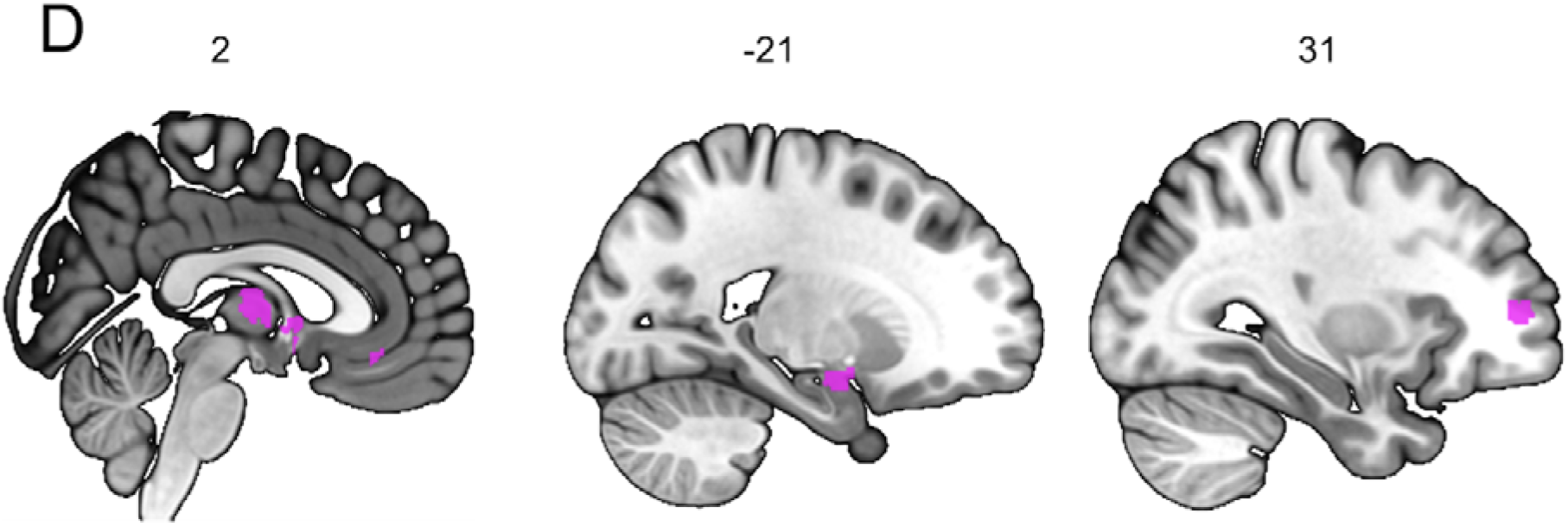
Association between Risky Behaviour and Imaging-Derived Phenotypes (IDPs) of Grey Matter Volume (GMV). **(A)** Loadings for the first principal component are extracted from four self-reported measures of risky behaviour in the *drinking, smoking, driving* and *sexual* domains (*N* = 315,855) (see Supplementary Figures 1 and 2 for descriptive statistics). We use this first principal component as a measure of risky behaviour. **(B)** Voxel-level GMV negatively associated with risky behaviour (*N* = 12,675). We observe associations in subcortical areas, including *thalamus, posterior hippocampus, amygdala, putamen, ventral striatum* and *cerebellum.* Associations with cortical areas include *posterior middle temporal gyrus, precentral gyrus, dlPFC, anterior insula* and *vmPFC.* **(C)** Associations between risky behaviour and GMV in 148 regions of interest (ROIs)’ (*N* = 12,675). The grey dotted line shows the FWE corrected threshold of *P* = 0.01 (see Methods for details). **(D)** The conjunction of GMV differences associated with risky behaviour reported here and a meta-analysis of 101 fMRI studies based on the key word “risky” reveals overlapping voxels in the thalamus, amygdala, vmPFC and dlPFC (see Methods for details).

Our main analysis includes a sample of 12,675 European-ancestry participants from the UKB. We first regress our measure of risky behaviour on total (whole-brain) grey matter volume (GMV) while controlling for age, birth year, gender, handedness, height, total intracranial volume and the first 40 genetic principal components, which account for genetic population structure (see Supplementary Methods 1.2). To exclude confounding effects of excessive alcohol consumption^21^, we excluded from the analysis all current or former heavy drinkers (see Methods)^22,23^. We find an inverse association between total GMV and risky behaviour (standardized *β* = -.122; 95% confidence interval (CI) [-.156, -.087]; *P* < 4.86 × 10^-12^, two-sided).

To identify specific brain regions related to risky behaviour, we perform a wholebrain voxel-based morphometry (VBM)^24^ analysis that regresses our measure of risky behaviour separately on GMV in each voxel across the brain, adjusting for the same control variables. We identify localized inverse associations between risky behaviour and GMV in distinct regions, only some of which were expected based on previous small-scale studies (see Figure 1B, Supplementary Figure 3 and Supplementary Table 4). In subcortical areas, we identify associations bilaterally in expected areas such as the amygdala and ventral striatum (VS), as well as in less expected areas such as the posterior hippocampus, putamen, thalamus, hypothalamus, and cerebellum. We also identify bilateral associations between risky behaviour and GMV in cortical regions that include the ventral medial prefrontal cortex (vmPFC), dorsolateral prefrontal cortex (dlPFC), ventro-anterior insula (aINS) and the precentral gyrus. In all of these regions, GMV is negatively associated with the propensity to engage in risky behaviours. We find no positive associations between GMV and risky behaviour anywhere in the brain.

To quantify effect sizes of the associations between risky behaviour and GMV in anatomically-defined brain structures and to investigate the convergence of our findings across MRI processing pipelines^25^, we conduct a follow-up analysis at the region of interest (ROI) level. This analysis primarily relies on the imaging-derived phenotypes (IDPs) provided by the UKB brain imaging processing pipeline^8,26,27^, which used parcellations from the Harvard-Oxford cortical and subcortical atlases and the Diedrichsen cerebellar atlas. We derived additional IDPs using unbiased masks based on the results of the voxel-level analysis (see Supplementary Methods 1.3.3). This analysis identifies negative associations between risky behaviour and GMV in 23 anatomical structures, with standardized *β*s between −0.079 and −0.036 (Figure 1C and Supplementary Figure 4), the largest of which is in the right ventro-aINS; (*β* = −0.079; 95% CI [-.103, -.055]; *P_uncorr_* = 1.34 × 10^-10^, two-sided).

We carry out several additional analyses to assess the differential contributions of various factors to the associations we observe. First, we re-estimate the ROI-level regressions with additional controls for various socioeconomic and cognitive outcomes that may be linked to both brain anatomy and risky behaviour, either as antecedents or downstream consequences. These controls include participants’ years of education and fluid intelligence (13-item measure)^17^, a zip-code level measure of the Townsend social deprivation index^28^, household income and size, and birth location (binned into 100 geographical clusters, see Supplementary Methods 1.2 and Supplementary Figure 1B). The direction and magnitude of all ROI effects are comparable to the main analysis (range of standardized *β*s between −0.079 and −0.035), with the largest effect again located in the right ventro-aINS (*β* = −0.079; 95% CI [-.104, -.055]; *P_uncorr_* = 3.3 × 10^-10^, two-sided). Furthermore, 19 of the 23 ROIs (all except the left Cuneal Cortex, left Crus I of the Cerebellum, left Planum Polare and the Brain Stem) are statistically significant after correction for multiple comparisons in this analysis (see Supplementary Figure 5 and Supplementary Table 5).

Second, we re-estimate our ROI-level regressions with additional controls for current levels of drinking (binned into 10 deciles) and smoking (binned into 3 categories). Although introducing these controls into the model regresses out variance of interest from the main outcome measure, which likely decreases the size of the effects, this analysis allows us to test whether any of the identified associations can reliably be attributed to risky behaviour that is not limited to the substance-use domain. In this analysis, we find that all of the effects originally identified remain negative in sign, yet are smaller (range of standardized *β*s between −0.041 and −0.011; see Supplementary Table 7). Nonetheless, the effects in 9 subcortical ROIs (including the Amygdala, Putamen, VS and Cerebellum) remain statistically significant after correction for multiple comparisons (see Supplementary Figure 6), with the strongest association identified in the left Amygdala (*β* = −0.041; 95% CI [-0.059, −0.023]; *P_uncorr_* = 1.1 × 10^-5^, two-sided). Thus, these subcortical IDPs are reliably associated with risky behaviour in non-substance use domains.

### Replication in an Independent Sample

Several months after the completion of our original analyses (in February 2020; https://biobank.ndph.ox.ac.uk/showcase/exinfo.cgi?src=timelines), the UKB released brain images of 20,316 additional participants— providing us with an opportunity to replicate our findings in an independent dataset that contains the same variables, and participants recruited in the same way from the same population^29^. After applying the same exclusion criteria as in our original analysis, our replication sample consists of 13,004 participants, roughly the same size as our original sample (see Methods for details). We repeat both the voxel-level and ROI-level analyses in this dataset. In the voxel-level analysis, we apply a significance threshold that corresponds to a familywise-error (FWE) rate of 5% in all voxels that showed significance in the original analysis (*p_uncorr_* = 2.956 x 10^-04^, with *t_uncorr_* = 3.62, two-sided). We find that 92.6% of the original voxels (located in 20 of the 21 clusters originally identified, with the exception of a cluster in cerebellar lobules I-IV) successfully replicate (see Supplementary Figure 7). Furthermore, the un-thresholded *t*-map^25^ of the original dataset strongly correlates with the un-thresholded *t*-map of the replication dataset (*r* = .767; 95% CI [.766, .768]; *P* < 10^-10^, two-sided). Likewise, our ROI-level analysis successfully replicates 21 of the original 23 ROI-level findings (*p_uncorr_* = 3.35 x 10^-03^, with *t_uncorr_* = 2.93, two-sided; see Supplementary Table 8). The two ROIs that do not replicate are the Cuneal Cortex (left) and the cerebellar lobule II (left).

### Overlap between Grey Matter Volume Differences and Functional MRI (fMRI) Meta-Analysis

To investigate whether the neuroanatomical associations of real-world risky behaviour correspond spatially with the localized activation patterns commonly identified in fMRI studies of risky decision-making, we conduct a conjunction analysis to compare our VBM results with data obtained from a publicly available meta-analysis of fMRI studies of risky behaviour^30^ (*N* = 4,717 participants, *K* = 101 individual studies, see Supplementary Table 9). The analysis reveals several brain regions whose anatomical associations with risky behaviour converge with functional engagement, including the thalamus, amygdala, vmPFC, and dlPFC (Figure 1D).

### Association of Polygenic Risk Scores for Risky Behaviour with Grey Matter Volume

Finally, we explore whether participants’ genetic disposition for risky behaviour, proxied via their polygenic risk scores (PRS), are associated with the neuroanatomical correlates of the trait, and test whether these associated neuroanatomical correlates mediate the relationship between genetic predisposition and behaviour. To this end, we first conducted a GWAS in an independent sample of UKB participants of European ancestry (*N*=297,025), exclusive of 18,796 genotyped individuals with usable MRI images (main sample) and their relatives. From the GWAS, we constructed a PRS that aggregated the effects of 1,176,729 single nucleotide polymorphisms (SNPs) on risky behaviour for all of the participants with MRI data in our independent target sample (see Supplementary Methods 1.4). The PRS predicts ~3% of the variance in risky behaviour in our target sample. Although the PRS are not associated with whole-brain GMV (standardized *β* = -.004; 95% CI [-.050 .020]; *P* > 0.41, two-sided), they are inversely associated with GMV in distinct regions, specifically the right dlPFC, right putamen and hypothalamus (Figure 2A.; regressions include all standard control variables, including total intracranial volume). Thus, GMV in these specific brain areas is negatively associated with the genetic disposition for risky behaviour.

**Figure 2.**
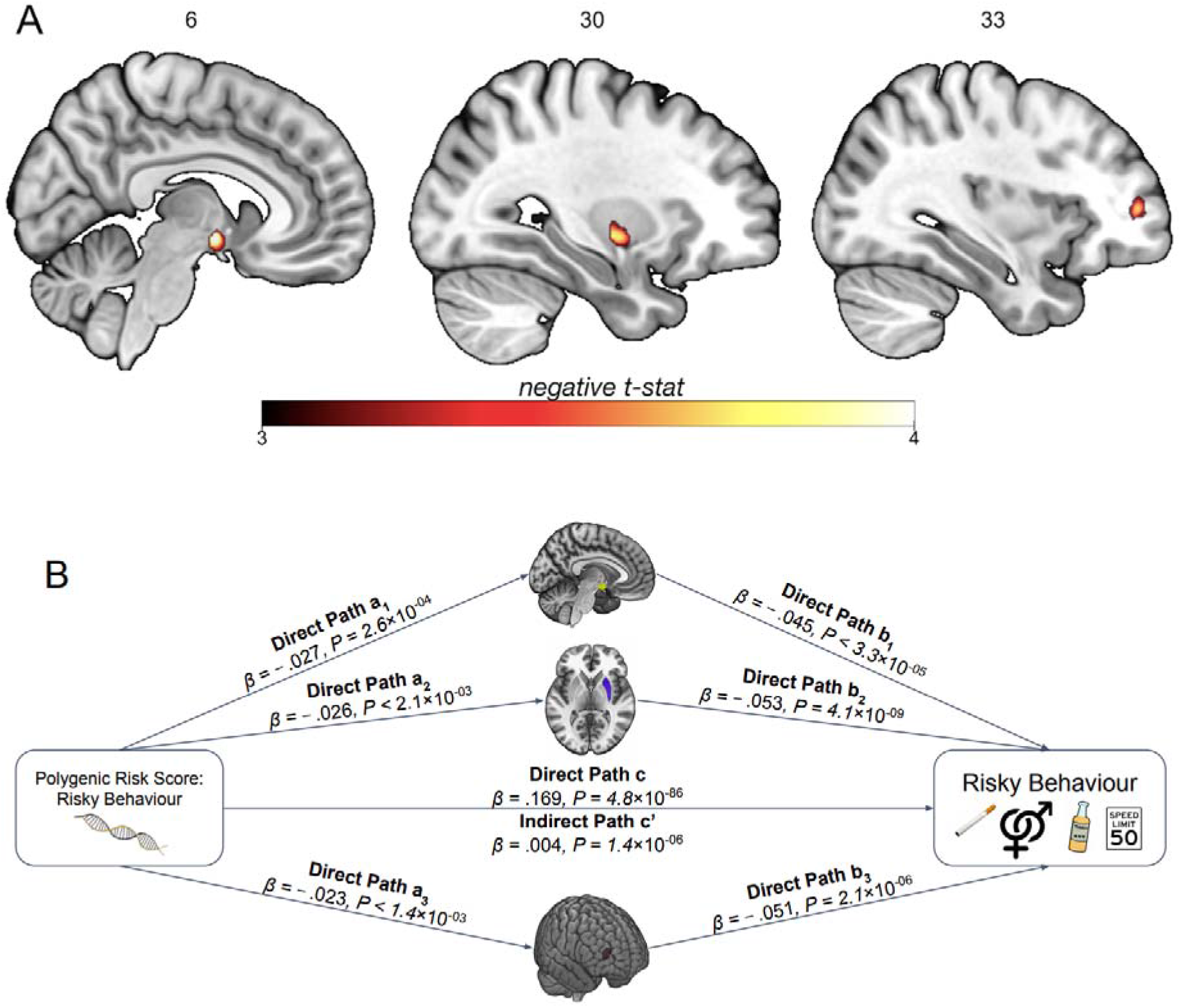
Association of Polygenic Risk Scores (PRS) for Risky Behaviour and Grey Matter Volume (GMV). **(A)** We constructed a PRS of risky behaviour from a GWAS in an independent sample (*N*=297,025) and investigated its associations with GMV in brain voxels that we identified as linked to risky behaviour. The PRS negatively correlates with GMV in the right dlPFC, putamen and hypothalamus. **(B)** GMV differences in hypothalamus (path 1, designated as a_1_ and b_1_), right putamen (path 2, designated as a_2_ and b_2_) and right dlPFC (path 3, designated as a_3_ and b_3_) mediate ~2.2% of the association between the PRS and risky behaviour. Arrows depict the direction of the structural equation modelling and do not imply causality (*N* = 12,675).

Based on these results, we use the previously extracted GMV of these three ROIs to examine whether it mediates the observed gene-behaviour associations. A structural equation model including all standard controls reveals that ~2.2% of the association between the PRS and risky behaviour is mediated through individual differences in GMV in the three regions (indirect path c’; standardized *β* = 0.004, 95% CI [.002, .005], *P* = 1.4 × 10^-06^, two-sided) (Figure 2B).

## Discussion

We investigate in a genetically-informed neuroimaging study (a) the association between GMV and real-world risky behaviour in a large population sample of European ancestry (main sample of 12,675 individuals and replication sample of 13,004 individuals), and (b) how the genetic disposition for risky behaviour is linked to GMV differences in a network of distinct brain areas. Several of the areas whose GMVs are linked to risky behaviour in this study have also often been functionally engaged during risky decision-making in small-scale fMRI studies that used stylized tasks. For instance, such correlations have been observed in the aINS, thalamus, dlPFC, vmPFC and VS^31,32^. These findings have led to proposals that upward and downward risks are encoded by distinct circuits, with upward risk mainly represented by areas encoding rewards (VS and vmPFC) and downward risk encoded by areas related to avoidance and negative arousal (aINS). Here, we substantiate previous functional studies with large-scale evidence that the structural properties of the same areas relate to risky behaviour in an ecologically valid setting^33^ when long-term health consequences are at stake.

Our results extend previous findings by showing that the neural foundation of risky behaviour is complex. Our analyses identify additional negative associations between risky behaviour and GMV in several areas, including the cerebellum, posterior hippocampus, hypothalamus, and putamen. While it is not yet clear how structural differences are manifested in properties of brain function^34^, our results suggest that risk taking draws on manifold neural processes, not just the representation and integration of upward and downward risks in the brain. Specifically, considering previous metaanalyses, the areas identified in this study are involved broadly in memory (posterior hippocampus), emotion processing (amygdala, ventro-aINS)^35^, neuroendocrine processes (hypothalamus)^36,37^, reward processing (vmPFC, VS and putamen)^38^ and executive functions (dlPFC)^39^. Thus, it appears that risky behaviour taps into multiple elements of human cognition, ranging from inhibitory control^40^ to emotion regulation^41^ and the integration of outcomes and risks^42^. This mirrors previous findings showing that risky behaviour is also a genetically complex trait^7^.

Additionally, our results underscore the long-suspected role of the hypothalamic–pituitary–adrenal (HPA) axis in regulating risk-related behaviours, in line with hormonal studies that link risky behaviour and sensation seeking to stress responsivity^36,37,43–45^. Furthermore, our finding that risky behaviour is linked to the structure of several cerebellar areas confirms the under-appreciated importance of the cerebellum for human cognition and decision-making, and highlights the need for further research on the specific behavioural contributions of this brain area^46^.

Of note, it remains an open question the extent to which the differences in GMV that we identify can be ascribed to specific heritable micro-anatomical traits, such as neuronal size, dendritic or axonal arborisation, or relative count of different cell types^47^. Moreover, it is not yet clear how these individual differences, in turn, influence behaviour. Nonetheless, our results provide evidence that neuro-anatomical structure constitutes the micro-foundation for neuro-computational mechanisms underlying individual differences in risky behaviours^34^.

Several of the observed neuroanatomical correlates of risky behaviour are also associated with the genetic disposition for the phenotype. Specifically, we find that GMV in the hypothalamus, the putamen and the dlPFC share variance with both risky behaviour and its polygenic risk score. This finding extends previous studies showing correlational^36,42,43^ and causal evidence^37,48^ of the involvement of these areas in risky behaviour by indicating that a genetic component partly underlies the associations. Although our analyses cannot identify the direction of the causal relationships (see Supplementary Discussion for an additional discussion of limitations), they show that risky behaviour and its genetic associations share variance with distinct GMV features and provide an overarching framework for how the genetic dispositions for risky behaviour may be expressed in the corresponding behavioural phenotype.

Our results are also in line with the bioinformatic annotation of the largest GWAS on risk tolerance to date, conducted in over 1 million individuals^7^, which implicated specific areas in the prefrontal cortex (BA9, BA24), striatum, cerebellum and the amygdala. However, bioinformatics tools used for GWAS annotation cannot be considered conclusive, as they rely on gene expression patterns in relatively small samples of (post-mortem) human brains or non-human samples^49^. Moreover, they cannot speak to whether changes in a particular tissue or cell type have negative or positive effects on the phenotype or how strong the effects are. Here, we show an alternative approach to annotating GWAS findings using a different type of data (large-scale population samples that include *in-vivo* brain scans and genetic data), relying on different assumptions than those used by bioinformatics tools. Our results add new insight by showing that lower GMV in specific brain areas is related to more risky behaviour and by implicating brain regions (i.e., putamen, hypothalamus, and dlPFC) in addition to those previously annotated. The effect sizes we observe here (standardized *β* < 0.08) are an order of magnitude larger than those found in GWAS on risky behaviour, but they nonetheless require very large samples for identification.

Finally, while many features of the brain are heritable, the environment indisputably plays an important role in brain development. We therefore see our results not as independent from, or of greater importance than, the effects of environmental and developmental factors. Rather, our study constitutes one step towards understanding how the complex development of human risky behaviour may be constrained by genetic factors.

## Methods

### Sample Characteristics and Selection Criteria

#### Main sample

We use publicly available data from the UKB, which recruited 502,617 people aged 40 to 69 years from the general population across the United Kingdom^19,20^. All UKB participants provided written informed consent and the study was granted ethical approval by the North West Multi-Centre Ethics committee. Our initial sample consists of 18,796 individuals with brain scans and genotype data, all of the imaged UKB participants as of October 2018. We excluded participants with putative sex chromosome aneuploidy (*N* = 6) or a mismatch between genetic and reported sex (*N* = 10), participants of non-European ancestry (*N* = 893), and participants who did not pass the UKB quality-control thresholds (*N* = 14), described in Bycroft et al.^27^. To minimize the potential influence of neurotoxic effects due to excessive alcohol intake^21,22^, we also excluded current heavy drinkers (531 females consuming more than 18 drinks per week and 793 males consuming more than 24 drinks per week)^22,23^. To exclude potential former drinkers, we also removed 426 participants who indicated that they did not drink alcohol.

All structural T1 MRI images used in the study underwent automated quality control by the UKB brain imaging processing pipeline^26^. We performed two additional quality checks using the Computational Anatomy Toolbox (CAT; www.neuro.uni-jena.de/cat/) for SPM 12 (www.fil.ion.ucl.ac.uk/spm/software/spm12/). First, we relied on the CAT12 automated Image and Pre-processing quality assessment, which included quality parameters for resolution, noise, and bias of images, 57 of which were automatically excluded and not pre-processed due to low image quality. Second, after pre-processing, we utilized an automated quality check of sample homogeneity to identify outliers that exhibited substantially different GMV patterns than the rest of the sample (see the CAT12 manual for details; http://www.neuro.uni-jena.de/cat12/CAT12-Manual.pdf, p. 17ff). In total, we excluded 690 individuals with scans of high image inhomogeneity (two standard deviations below the mean). Finally, we excluded 2,701 participants with incomplete behavioural data of interest or control variables. Our final dataset consists of *N* = 12,675 individuals.

#### Replication sample

The initial replication sample consisted of 20,316 individuals with usable brain scans and genotype data, all of the imaged UKB participants as of February 2020 who were not included in our main sample. Following the original analysis, we excluded participants with putative sex chromosome aneuploidy (*N* = 7), a mismatch between genetic and reported sex (*N* = 11), non-European ancestry (*N* = 1,143), heavy drinking or abstinence from alcohol (*N* = 1,376), and those that did not pass the UKB quality-control thresholds (*N* = 31)^27^. All T1 MRI images used in the study underwent the same quality control procedure as in our original sample, resulting in the removal of additional 391 individuals. Finally, we excluded participants with incomplete data (*N* = 4,353). Our final replication sample consists of *N* = 13,004. The empirical distributions of the main variables used in our replication analysis are depicted in Supplementary Figure 1B.

### Measures

#### Risky Behaviour

We closely follow Karlsson Linnér et al.^7^ to derive a measure of risky behaviour across domains based on participants’ self-reports of: (1) Number of alcoholic drinks per week, (2) Ever having smoked, (3) Number of sexual partners, and (4) Frequency of driving faster than the motorway speed limit (see Supplementary Methods 1.1). We perform principal component analysis (PCA) on N = 315,855 UKB participants and extract the first principal component (PC) of the four measures as the main outcome of interest (referred to as “risky behaviour”). The first PC explains ~37% of the variance in the four measures, and it is the only PC that loads positively on all of them. Summary statistics and factor loadings of the PCA are available in Tables S1 and S2. The code for generating the variable of ‘risky behaviour’ is accessible at https://osf.io/qkp4g/.

#### Control variables

The full list of control variables and the methods used to generate them are available in Supplementary Methods 1.2.

#### T1 MRI Image Processing

We use T1-weighted structural brain MRI images in NIFTI format provided by the UKB. Images were acquired using 3-T Siemens Skyra scanners, with a 32-channel head coil (Siemens, Erlangen, Germany), with the following scanning parameters: repetition time = 2000 ms; echo time = 2.1 ms; flip angle = 8°; matrix size = 256 × 256 mm; voxel size =1 × 1 × 1 mm; number of slices = 208. A detailed description of the methods used to pre-process the images and derive voxel-level and ROI-level IDPs is available in Supplementary Methods 1.3.1-1.3.3.

#### Polygenic Risk Score (PRS) for Risky Behaviour

To construct a PRS, we first re-estimated the GWAS of our main measure described in ref^7^ after excluding 18,796 genotyped individuals with usable T1 MRI images and their relatives up to the third degree (final GWAS sample: *N* = 297,025 individuals of European ancestry). The GWAS was performed using linear mixed models (LMM), implemented via BOLT-LMM version 2.3.2^50^. Next, we performed quality control (QC) of the GWAS results using a standardized QC protocol, described in detail in ref^7^. This protocol removes rare and low-quality single-nucleotide polymorphisms (SNPs) based on minor allele frequency (MAF) < 0.001, imputation quality (INFO) < 0.7, and SNPs that could not be aligned with the Haplotype Reference Consortium (HRC) reference panel, among other filters. After QC, a total of 11,514,220 SNPs remained in the GWAS summary statistics. Thereafter, we calculated a PRS for each participant by weighting his or her genotype across SNPs by the corresponding regression coefficients estimated in the GWAS (see Supplementary Methods 1.4 for further information).

### Analysis

#### Voxel-based Morphometry (VBM)

We identify associations between risky behaviour and localized GMV across the brain using whole-brain voxel-based morphometry (VBM), a method that normalizes the anatomical brain images of all participants in one stereotactic space^24^. We regress risky behaviour separately on each voxel of the smoothed GMV images (see Supplementary Methods 1.3.1) and the control variables. We correct for multiple comparisons by adjusting the FWE rate to *α* = 0.01 using permutation tests (*P_uncorr_* = 1.248 x 10^-06^, with |*t_uncorr_*| = 4.85, two-sided). See Supplementary Figure 3 and Supplementary Table 4 for the summary statistics of each cluster and the coordinates of the peak voxel within that cluster.

#### Region of Interest (ROI)-level Analysis

We compute the associations between risky behaviour and 139 IDPs of GMV extracted by the UKB brain imaging processing pipeline^26^ using parcellations from the Harvard-Oxford cortical and subcortical atlases and Diedrichsen cerebellar atlas, in addition to 9 IDPs derived using unbiased masks based on the results of our voxel-level analysis (see Supplementary Methods 1.3.2 and 1.3.3). We regress risky behaviour separately on each IDP and the control variables and correct for multiple comparisons by adjusting the FWE rate of *α* = 0.01 using permutation tests (*p_uncorr_* = 9.37 x 10^-05^, with |*t_uncorr_*| = 3.91, two-sided).

#### Replication of the Voxel-level and ROI-level Analyses

We repeat the VBM analyses in the replication sample for all voxels that showed significance in the original sample, correcting for multiple comparisons by adjusting the FWE rate to α = 0.05 (two-tailed) using permutation tests (*p_uncorr_*= 2.956 x 10^-04^, with *t_uncorr_*= 3.62). We also repeat the ROI-level analysis in all ROIs that showed significance in the original sample, and correct for multiple comparisons by adjusting the FWE rate to α = 0.05 (two-tailed) using permutation tests (*p_uncorr_* = 3.35 x 10^-03^, with *t_uncorr_* = 2.93). We define replication success as observing a statistically significant effect at the 5% level (corrected for multiple hypothesis testing) in the same direction as the original finding^51^.

#### Comparison of VBM Results with a Meta-Analysis of Functional MRI (fMRI) Studies

We compare our VBM results with a publicly available meta-analysis of fMRI studies provided by Neurosynth^30^, an online platform for large-scale, automated synthesis of fMRI data (https://neurosynth.org/). The meta-analysis was based on the keyword ‘risky’. It consisted of *K* = 101 individual studies with a total of *N* = 4,717 participants (see Supplementary Table 9 for details), and was conducted using a uniformity test (assuming that random activations are evenly distributed across all voxels). The meta-analytic statistical image was corrected for multiple comparisons by applying a false discovery rate (FDR) of .01 (implemented by Neurosynth). The summary of studies included in the meta-analysis is available on Supplementary Table 9. The 3D activation map that resulted from the meta analysis is available on https://neurosynth.org/analyses/terms/risky/. To compare our VBM results (i.e. brain structure) with the meta-analysis of data on brain function, we perform a whole-brain voxel-level conjunction analysis of the two (see Supplementary Figure 8) that exhibits the spatial overlap of all voxels that are significant in both analyses (Figure 1D). Thus, both the structure and function of the brain regions identified by this conjunction are significantly associated with risky behaviour.

#### Voxel-based Morphometry (VBM) Analysis of Risky Behaviour PRS

We repeat the VBM analysis in all voxels identified to be associated with risky behaviour, using the PRS as the dependent variable. This approach allows the identification of brain regions that were likely to mediate the effect of genetic disposition on risky behaviour. The PRS was constructed using GWAS results from an independent sample, to ensure that the effect size estimates in this analysis are not inflated due to overfitting. We account for multiple comparisons using a permutation test with an FWE rate of *P_FWE_* < 0.05. This part of the analysis was not pre-registered.

#### Mediation Analysis

We conduct mediation analyses (implemented in STATA 14) to test whether GMV differences in the brain regions whose GMVs are associated with the PRS of risky behaviour (right dlPFC, right putamen and hypothalamus) mediate the association between the PRS and risky behaviour. We first extract GMV from these three ROIs using the same unbiased masks as in the ROI-level analysis (see Supplementary Methods 1.3.3). Then, we estimate a structural equation model (SEM) to quantify the effect of the PRS on risky behaviour mediated via GMV differences in these ROIs. All SEM equations include the aforementioned standard control variables (listed in Supplementary Methods 1.2). We carry out an additional robustness check by estimating an SEM that assumes one single path (i.e., the sum of all ROIs), which yields the same pattern of results (see Supplementary Figure 9).

#### Family-Wise-Error Correction using Permutation Tests

To account for multiple hypothesis testing, we determine the appropriate FWE corrected *p*-value threshold with a permutation test procedure in each of our analyses^52^. To this end, we generated 1,000 datasets with randomly permuted phenotypes (i.e., breaking the link between the outcome and explanatory variables), estimated regression models for all IDPs per analysis, and recorded the lowest *p*-value of each run to generate an empirical distribution of the test statistic under the null hypothesis. To obtain the FWE rate of any given alpha, we use the *n*^th^ = ⍰ x 1000 lowest *p*-value from the 1,000 permutation runs as the FWE corrected *p*-value threshold.

#### Pre-registration of Analysis Plan and Unplanned Deviations

We pre-registered our analysis plan on Open Science Framework (OSF, https://osf.io/qkp4g/, registered Dec. 2018). Our pre-registered plan specified the construction of the dependent variable, the control variables, the inclusion criteria and quality controls, the VBM analyses and the main ROI-level analyses. We summarize all deviations from the analysis plan in Supplementary Methods 2.

## Data Availability

Data and materials are available via UK Biobank at http://www.ukbiobank.ac.uk/.

## Code Availability

The analysis code used in this study is publicly available on https://osf.io/qkp4g/.

## Acknowledgments

This research was carried out under the auspices of the Brain Imaging and Genetics in Behavioural Research (https://big-bear-research.org/) consortium. The authors thank Nadja C. Furtner for helpful comments and Dylan Manfredi for research assistance. The research was conducted using UK Biobank resources under application 40830. The study was supported by funding from an NSF Early Career Development Program grant (1942917) and The Wharton School Dean’s Research fund to G.N., an ERC Consolidator Grant to P.D.K. (647648 EdGe), and a Swiss National Science Foundation grant to C.C.R (100019L_173248). R.R.W. was financially supported by NIAAA K23 grant (K23 AA023894) and H.R.K. was supported by NIDA grant P30 DA046345. G.N. thanks Carlos and Rosa de la Cruz for ongoing support. The work was carried out on the Dutch national e-infrastructure with the support of the SURF Cooperative. Data can be accessed via the UKBiobank, and data analysis scripts are available on OSF (https://osf.io/qkp4g/).

## Author contributions

G.A., R.D., J.W.K., P.D.K. and G.N. designed the research plan. G.N., P.D.K. and C.C.R. oversaw the study. G.A., R.D. and R.K.L analysed the data with critical input from P.D.K., G.N. and C.C.R.. G.A. G.N. and P.D.K. wrote the paper and Supplementary Materials. All authors contributed to and critically reviewed the manuscript.

## Supplementary Methods

### 1. Measures

#### 1.1. Main risky behaviour measure

We closely follow the methods of ref 1 to derive a measure of risky behaviour based on participants’ self-reports across the drinking, smoking, driving, and sexual domains.

Specifically, we use the following UK Biobank variables:

- Number of alcoholic drinks per week (Data-Fields: 1558, 1568, 1578, 1588, 1598, 1608, 5364, 4407, 4418, 4429, 4440, 4451, 4462)
- Ever smoking (Data-Field 20116, 1249, 1239)
- Frequency of driving faster than the motorway speed limit (Data-Field 1100)
- Lifetime number of sexual partners (Data-Field 2149)^1^

The full description of each Data-Field can be found in the online data showcase of the UKB (http://biobank.ctsu.ox.ac.uk/crystal/search.cgi). The annotated STATA code used to derive all behavioural phenotypes and control variables can be found in our pre-registered analysis plan (https://osf.io/qkp4g/).

All variables above were measured on at least one of 3 occasions: (1) the initial assessment visit, (2) the first repeat assessment visit, and (3) the imaging visit. Data from (2) and (3) are only available for a subset of the original sample. In cases where participants provided answers across more than one visit, we compute the average of their reports.

To obtain a measure that captures the common variance in risky behaviour shared across domains, we perform principal component analysis (PCA) on *N* = 315,855 UKB participants and extract the first principal component (PC) as our main outcome of interest for this study (referred to as “risky behaviour”). Compared to experimental procedures that elicit risk tolerance, selfreported measures exhibit higher external validity and test-retest reliability^3–5^. Furthermore, by extracting the first principal component of the four risky behaviours, we reduce measurement noise due to the aggregation of signals across various measures, while capturing behavioural tendencies across domains that are independent of idiosyncratic differences in the four specific behaviours. The PCA summary statistics are available in Supplementary Table 1, and the component loadings are available in Supplementary Table 2. The first PC explained about 37% of the variance in the different phenotypes of risky behaviours in the sample, and it was the only PC that positively loaded on all of four phenotypes.

While the GWAS by Linnér et al. (2019)^1^, was primarily based on a meta-analysis of two very crude, noisy, single-item measures of risk taking that were available in the two largest samples (UKB and 23andMe, which had slightly different questions on risk taking), this choice (in the GWAS) was made to maximize the sample size for genetic discovery, following the logic outlined in ref 6, i.e., that in genetic discovery studies, sample size typically trumps phenotypic accuracy in terms of statistical power. (The supplementary material of ref 6 includes a mathematical derivation that illustrates this). In the current work, we also wanted to maximize statistical power, albeit the situation here is different, as the sample size was exogenously determined by the UKB. Thus, the only means to increase statistical power was via increasing the quality of the phenotypic measurement. We decided to focus on the first PC of the four risky behaviours introduced in Linnér et al. (2019) for the following reasons: (1) It is available for a large part of the scanned subsample. (2) Linnér et al. (2019) showed that this first PC has a higher SNP-based heritability than any of the general risk-taking measures or individual phenotypes, which is partly because the first PC is less affected by random measurement error than any input variable considered separately. (3) The high heritability of the first PC suggests that similar genetic factors influence risk taking across various domains, making this a promising trait to study in connection with other biomarkers such as brain anatomy. (4) This variable has been studied in the literature, limiting our degrees of freedom for the current study. (5) GWAS results for this variable were readily available.^6^

#### 1.2. Control Variables

All of our analyses systematically control for several genetic, socio-demographic and anthropometric factors that could potentially confound the observed associations [e.g. sex^7^, height^8^ and genetic population structure^9^]. Specifically, we use the following control variables, as provided by the UKB:

- Age at the time of brain scan (Data-Field 21003)
- Birth year (Data-Field 33)
- Sex (self reported and genetically identified, Data-Fields 31 & 22001, dummy coded)
- Height (Data-Field 50)
- Handedness (Data-Field 1707, categorical variable: Right-handed, Left-handed, ambidextrous, N/A)
- Sex x birth year interactions (binned into fields containing at least 20 participants each)
- The first 40 PCs of the genetic data (Data-Field 22009)
- Total intracranial volume (TIV), derived using the CAT12 toolbox from T1 images.

We carry out an additional analysis that further controls for the following socio-economic and cognitive outcomes (provided by the UKB):

- Educational attainment (Data-Field 6138)
- A 13-item measure of fluid IQ (Data-Fields 20016 and 20191)
- Zip-code level measure of the Townsend social deprivation index (Data-Field 189)
- Household income (Data-Field 738)
- Number of household members (Data-Field 709)
- Place of birth, binned in 100 clusters based on North and East birth location coordinates (Data-Fields 129 and 130). Clusters were calculated using the *k*-means algorithm, which minimizes within-cluster variances (squared Euclidean distances) of *k* = 100 clusters with 10,000 iterations after random seeding.

The empirical distributions of the main variables used in our main analysis and the correlations between them are depicted in Supplementary Figures 1A-C and Supplementary Figure 2.

#### 1.3. Imaging-derived Phenotypes (IDPs)

##### 1.3.1 T1 MRI Image Processing

Our voxel-level analysis uses T1-weighted structural brain MRI images in NIFTI format provided by the UKB (data field 20252). The images were acquired using 3-T Siemens Skyra scanners, with a 32-channel head coil (Siemens, Erlangen, Germany), with the following scanning parameters: repetition time = 2000 ms; echo time = 2.1 ms; flip angle = 8°; matrix size = 256 × 256 mm; voxel size = 1 × 1 × 1 mm; number of slices = 208.

We preprocessed the data using the Computational Anatomy Toolbox (CAT; www.neuro.uni-jena.de/cat/) for SPM (www.fil.ion.ucl.ac.uk/spm/software/spm12/), a fully automated toolbox for deriving neuroanatomical measurements at voxel and ROI levels. Image pre-processing used the default setting of CAT12 (accessible online at http://www.neuro.uni-jena.de/cat12/CAT12-Manual.pdf). Images were corrected for bias-field inhomogeneities, segmented into gray matter, white matter, and cerebrospinal fluid (CSF), spatially normalized to the MNI space using linear and non-linear transformations, and were modulated to preserve the total amount of signal in the original image during spatial normalization (the specific SPM-processing parameters can be found in the pre-registered document on OSF https://osf.io/qkp4g/). We applied spatial smoothing with 8-mm Full-Width-at-Half-Maximum (FWHM) Gaussian kernel for the segmented, modulated images for grey matter volume (GMV). Finally, to ensure that only voxels that likely contain grey matter enter the analyses, we constructed a brain mask based on the average of all GMV images. Specifically, following standard VBM procedures (see SPM/CAT12 http://www.neuro.uni-jena.de/cat12/CAT12-Manual.pdf) we thresholded the average of all brain images at 250 GMV intensity units. The resulting image was binarized and applied as a pre-mask to all individual images before running analyses. Additionally, on an individual level, we excluded all voxels that exhibited a lower grey matter volume than .1 from the analyses (see standard parameters of SPM/CAT12 http://www.neuro.uni-jena.de/cat12/CAT12-Manual.pdf).

##### 1.3.2 Region of interest (ROI)-level IDPs Processed by the UKB

We use all of the GMV IDPs that were processed and provided by the UKB [for details see ref 10]. These IDPs include GMV of 139 ROIs derived using parcellations from the Harvard-Oxford cortical and subcortical atlases, and Diedrichsen cerebellar atlas.

##### 1.3.3 Additional ROI-level IDPs

Based on our voxel-level results (see 2.1), we extracted 5 additional ROI-level IDPs that quantified GMV in anatomical substructures that were not derived by the UKB. These ROIs were extracted bilaterally from unbiased masks and included the dorsolateral prefrontal cortex (dlPFC; BA 46), hypothalamus, posterior hippocampus, ventro-anterior insula and ventromedial prefrontal cortex (vmPFC). For the dlPFC, ventro-anterior insula and vmPFC masks, we used recent functional parcellations based on resting state data. The dlPFC mask was derived using the Sallet Dorsal Frontal resting state connectivity-based parcellation (cluster 7/BA46)^11^. Functionally, this area exhibits coupling with the frontal-parietal network (incl. anterior cingulate cortex, parietal cortex and inferior parietal lobe), as well as with the vmPFC. Anatomically, its boundaries show resemblance to BA 46 — an area functionally related to executive function that shows distinct cytoarchitectonic properties.

We extracted GMV from the vmPFC using a parcellation of the medial wall of the prefrontal cortex, based on resting state functional coupling^12^. Specifically, we extracted GMV from 14m — an area linked to cost-benefit integration in value-based decision-making^13–16^, which maintains strong positive coupling with hypothalamus, ventral striatum, and amygdala^17^. The hypothalamus mask was derived from a high-resolution atlas of human subcortical brain nuclei^18^. The posterior hippocampus mask was derived according to recent recommendations for long-axis segmentation of the hippocampus in human neuroimaging^19^. We labeled hippocampal voxels posterior to the coronal plane at *y* = −21 in MNI space (which corresponds to the uncal apex of the parahippocampal gyrus), as posterior hippocampus. To ensure spatial precision across participants, we used a minimum 80% likelihood of each voxel being in the anatomical structure for all of the aforementioned masks. The ventro-anterior insula mask was derived following a recent parcellation of the insula based on a resting state functional connectivity analysis by ref 20, which reported that this brain region showed functional coactivation with limbic areas including amygdala, ventral tegmental area (VTA), superior temporal sulcus, and posterolateral orbitofrontal cortex. The raw mask was thresholded at *z* = 10.

##### 1.4 Polygenic Risk Score (PRS) for Risky behaviour

We use the genetic data provided by the UKB to construct a polygenic risk score (PRS) for risky behaviour. As a first step, we rerun the genome-wide association study (GWAS) of risky behaviour (the same measure used in the current study) as reported in ref 1 after excluding the 18,796 genotyped individuals with usable T1 NIFTI structural brain images (UKB field 20252) and all of their relatives up to the third degree (defined using the KING coefficient^21^ based on a pairwise coefficient >0.0442). The final GWAS sample includes 297,025 individuals of European ancestry. We use BOLT-LMM version 2.3.2^22^ to perform GWAS with linear mixed models (LMM), which outperforms linear regression in terms of statistical power and controlling for relatedness^23^.

Next, we perform quality control (QC) of the GWAS results using a standardized QC protocol, described in detail in ref 1. This protocol removes rare and low-quality single-nucleotide polymorphisms (SNPs) based on minor allele frequency (MAF) < 0.001, imputation quality (INFO) < 0.7, and SNPs that could not be aligned with the Haplotype Reference Consortium (HRC) reference panel. After QC, a total of 11,514,220 SNPs remains in the GWAS summary statistics.

Thereafter, we calculate for each participant *i* a PRS, *S_i_* by weighting his or her genotype across SNPs (*j*), *g_ij_*, by the corresponding regression coefficients, *β_j_* estimated in the GWAS described above. Thus, the PRS is a linear combination of genetic effects, calculated as:

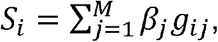

where the set of SNPs, *M*, is restricted to the consensus genotype set of 1.4 million SNPs established by the International HapMap 3 Consortium^24^, which has been successfully employed for polygenic prediction in many previous studies. Furthermore, the PRS is constructed only with autosomal, bi-allelic SNPs with *MAF* > 0.01 and *INFO* > 0.9 in the UKB. The resultant PRS is based on a total of *M*=1,176,729 SNPs. The PRS is then standardized to mean zero and unit variance in the prediction sample.

#### 1.5 Genetic Correlations of Risky behaviour

We rely on the results of the risky behaviour GWAS to estimate genetic correlations between this phenotype and 85 other traits, using bivariate LD Score regression^25^. The estimates are reported in Supplementary Table 3. For this purpose, we query the “GWAS ATLAS”^26^ to identify publicly archived GWAS results that we consider relevant. We supplement the publicly available GWAS with a soon-to-be published GWAS on diet composition^27^. Notably, the collected traits span across many different outcomes, including the anthropometric, behavioural, cognitive, psychiatric, medical, and socioeconomic domains.

We find moderate to strong genetic correlations between our main measure and a range of phenotypes that are considered risky behaviours, including ever consuming cannabis (*r_g_* = 0.72; *SE* = 0.03), self-employment (*r_g_* = 0.52; *SE* = 0.30), and age at first sexual experience (*r_g_* = −0.54; *SE* = 0.02). Our measure of risky behaviour is also genetically correlated with a range of mental disorders including bipolar disorder (*r_g_* = 0.23; *SE* = 0.03), major depressive disorder (*r_g_* = 0.22; *SE* = 0.03), and schizophrenia (*r_g_* = 0.17; *SE* = 0.02). Finally, risky behaviour is genetically correlated in the expected direction with the personality traits of conscientiousness (*r_g_* = −0.25; *SE* = 0.10) and extraversion (*r_g_* = 0.34; *SE* = 0.05).

### 2. Pre-registration of Analysis Plan and Unplanned Deviations

We pre-registered our analysis plan on Open Science Framework (OSF, https://osf.io/qkp4g/). Our pre-registered plan specifies the construction of the dependent variable, the control variables, the inclusion criteria and quality controls, the VBM analyses and the main ROI-level analyses.

We deviated from the pre-registered plan in several cases, which are outlined in the following. These deviations occurred when the computational burden of following the preregistered plan was unexpectedly high, and when alternative measures that we were not aware of at the time of the pre-registration were made available by the UKB. Specifically, we decided not to use alternative segmentations of the cortex (e.g. Hammer’s atlas) as robustness checks for our ROI-level analysis because of the significant computational burden in deriving those measures. Instead, based on the voxel-level analysis, we derived additional ROIs only when they were not derived in sufficient granularity in the IDPs provided by the UKB (see 1.3.3).

Similarly, we did not derive cortical thickness (CT) measures because of the high computational burden using FreeSurfer, which is the gold standard in cortical thickness estimation. While other means to derive CT would have been available (e.g. CAT toolbox), they would provide relatively lower quality data and would not allow analyses of subcortical areas. Additionally, the UKB was expected to release CT measures derived from FreeSurfer before this work was finalized (see the UKB Data Showcase website for public announcements). The lack of CT measures has also led us to decide to postpone the conduct of an additional preregistered multivariate analysis.

Finally, our pre-registered plan states that we would run additional robustness checks to control for potential neurotoxic effects of excessive alcohol intake. Upon examining the data for a different project that is focused on the effects of alcohol intake on the brain, we observed effects that were mainly driven by individuals who were heavy drinkers. We therefore decided to deviate from our original plan and exclude all participants who qualified as current or former regular heavy drinkers. However, we also provide additional analyses that include weekly alcohol intake and smoking habits as a covariate (see Supplementary Figure 6). Finally, the preregistered analysis of white-matter volume is not reported here, because we decided to focus our manuscript on GMV differences.

## Supplementary Discussion

Our study highlights the importance of using large samples to study associations of neuroanatomy with complex behavioural traits. The largest effect we identify for the relationship between any cluster of voxels and risky behaviour is Δ*R*^2^ = 0.6% (see Supplementary Table 4). It would require more than 1,750 participants to have 90% statistical power at a liberal *p*-value threshold of 0.05 (uncorrected) to identify effects of this magnitude. This is a lower bound for the required sample size for such studies that does not reflect the upward bias in our effect size estimate due to the statistical “winner’s curse”, and the need to correct for multiple testing. Previous large-scale VBM studies (*N* > 1000) with other behavioural phenotypes^28^ found effect sizes of similar magnitude and suggest that large samples are a prerequisite to detect such an association reliably. Of note, the largest previous study of risk tolerance employed a sample of 108 participants^29^ and would have only 12% power to detect Δ*R*^2^ = 0.6% at *α* = 0.05 (uncorrected).

A possible limitation of our study is that, the specific features of the component phenotypes (e.g., smoking) rather than their first PC (risky behaviour) could have driven the associations we report (quantified via standardized regression coefficients). To further investigate this possibility, we repeat our ROI-based analysis with the individual phenotypic measures (instead of their first PC) as outcome variables (see Supplementary Table 6). We find that 22 out of 23 ROIs are significantly associated with more than one phenotype (the exception is IX Cerebellum (r), which is significantly associated only with the number of sexual partners, yet the standardized coefficient denoting its relationship with the first PC is greater in magnitude than the coefficient denoting its relationship with the number of sexual partners). Furthermore, the standardized coefficients quantifying the relationships between the ROIs and the individual phenotypes are either smaller than or at the same order of magnitude as the coefficients quantifying their relationship with the first PC.

While our study is larger and more representative than any previous investigation of the topic, and although we control for various potential confounds and replicate our findings in an independent sample, it was conducted in a population of UK individuals of European descent that were over 40 years old at the time of measurement, which limits the generalizability of our results to other populations. Moreover, our results do not exclude the possibility of bias due to other unobserved variables that our analyses do not account for. With the rise of large publicly available data sets [e.g. ref 30], we hope that future studies will be able to test the generalizability of our findings to populations of different ethnicities and age groups (e.g., adolescents).

Finally, while our analyses identify distinct brain areas that mediate gene-phenotype associations for risky behaviour (i.e., putamen, hypothalamus and dlPFC), they do not provide evidence for their causal relationship. For instance, it is possible that a person’s genetic disposition would lead them to select into environments that influence both risky behaviour and features of brain anatomy.

## Supplementary Figures

**Supplementary Figure 1A.**
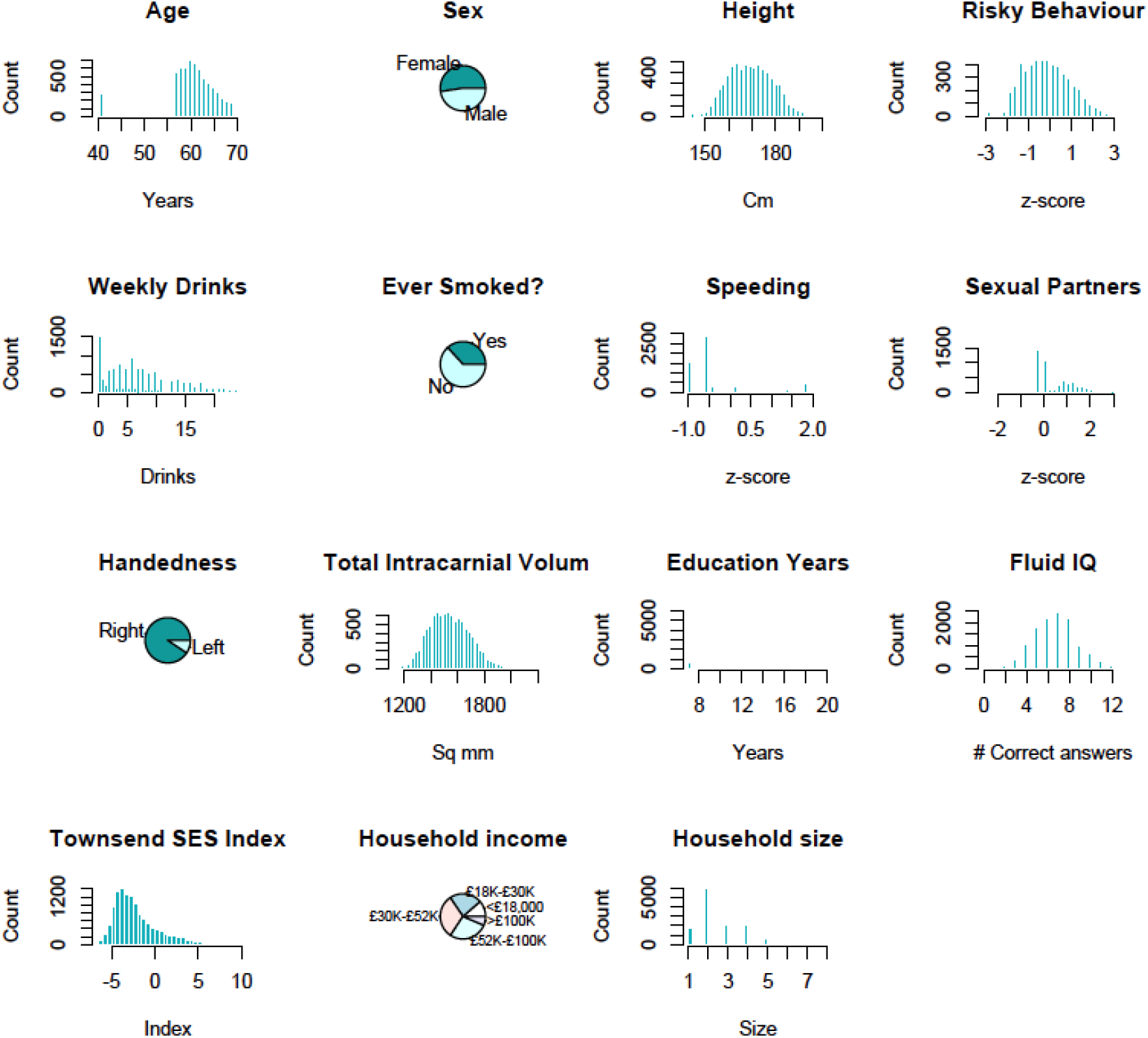
Empirical distributions of variables in the main study sample (*N*=12,675).

**Supplementary Figure 1B.**
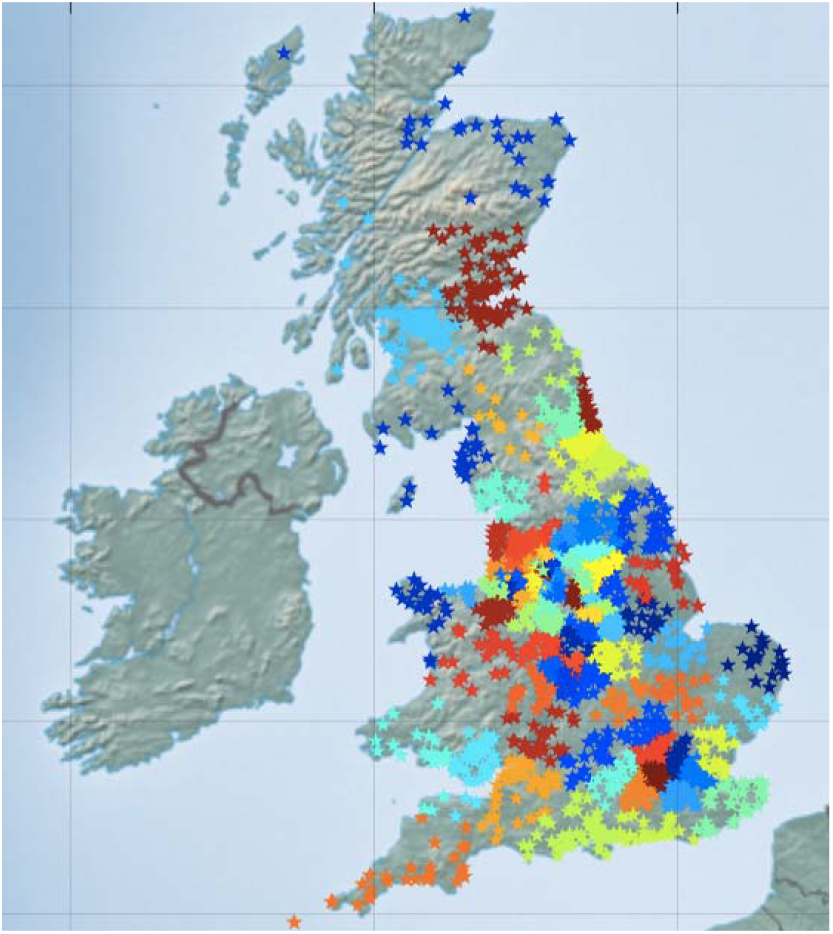
Geographical birth location clusters of the study’s participants (main sample). Each star represents the birthplace of a participant (non-jittered). Colors denote 100 geographical clusters, calculated using a *k*-means clustering algorithm with *k* = 100 and 10,000 iterations after random seeding (*N*=12,675).

**Supplementary Figure 1C.**
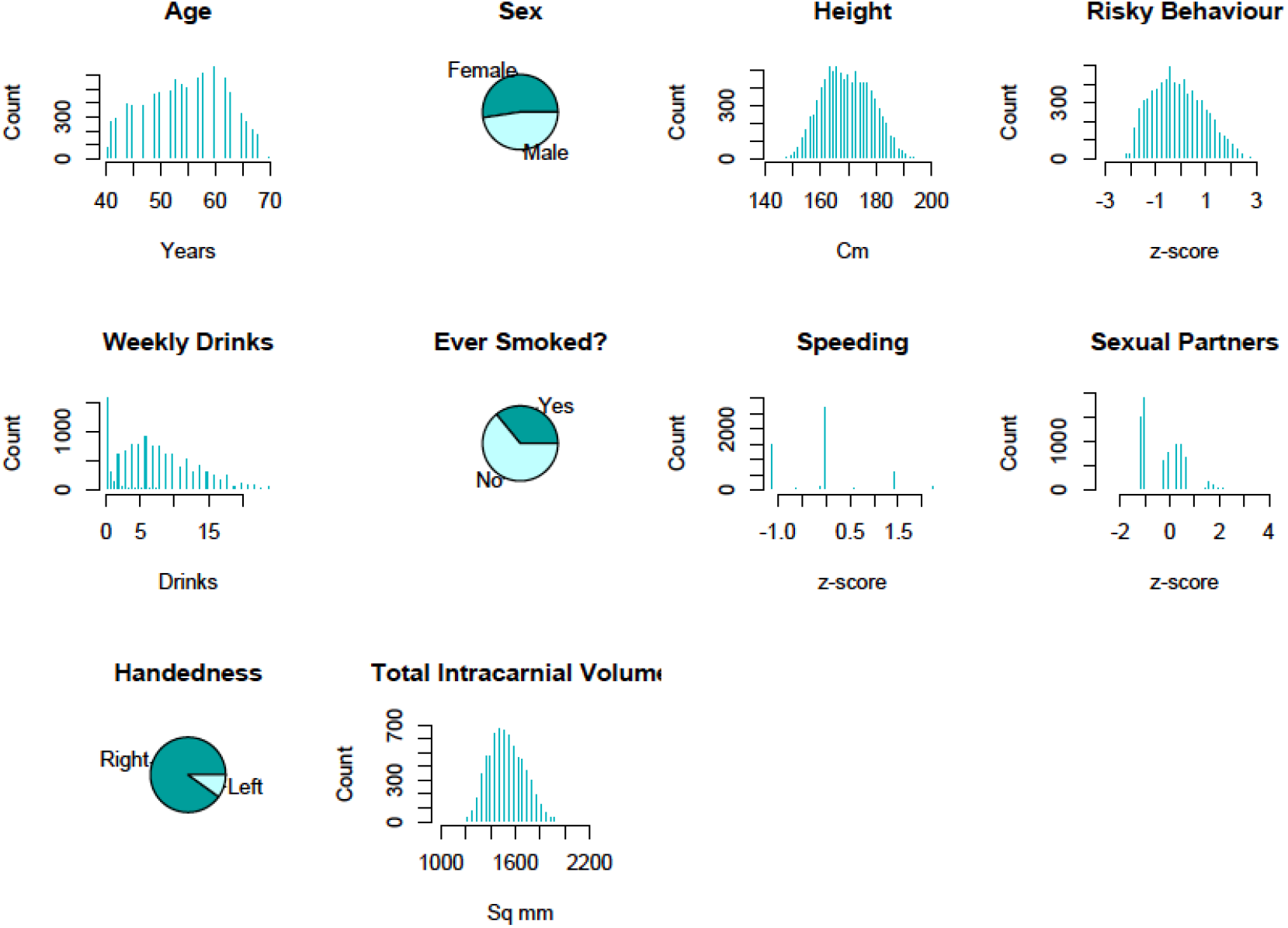
Empirical distributions of variables in the replication sample (*N*=12675).

**Supplementary Figure 2.**
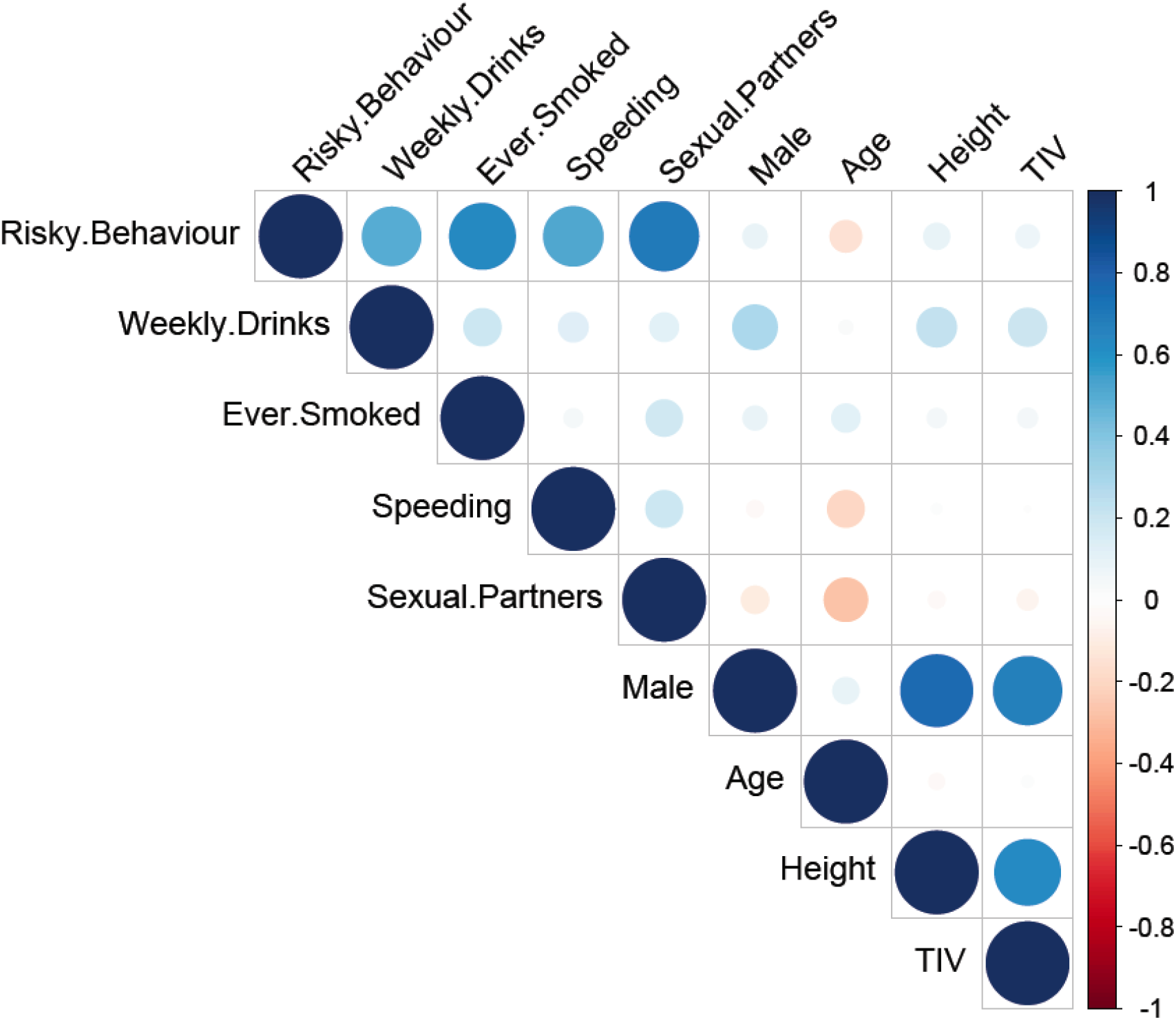
Bivariate correlations between variables used in the main study sample(*N*=12,675).

**Supplementary Figure 3.**
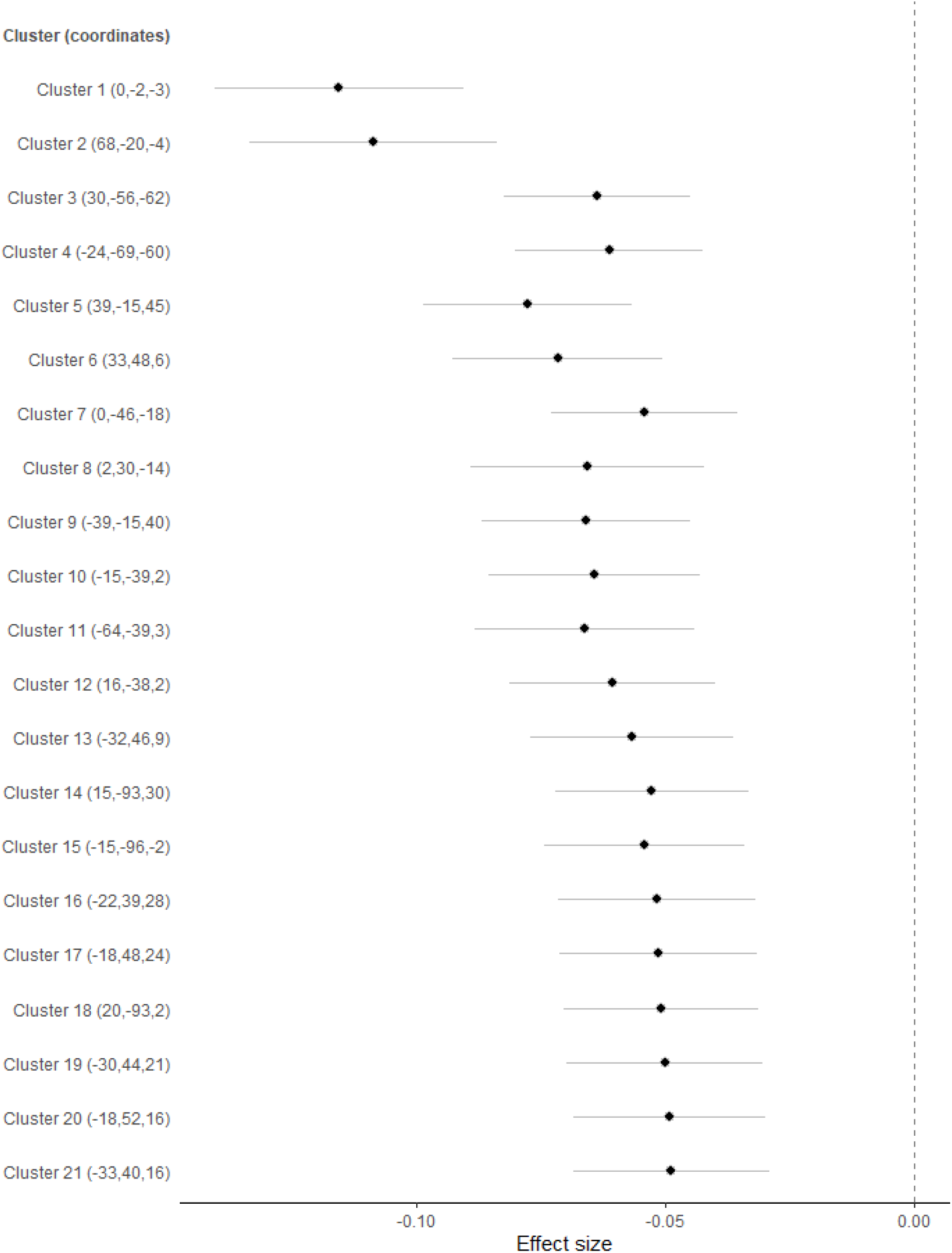
Effect sizes (standardized betas) of associations between risky behaviour and grey matter volume (GMV) in voxel clusters showing significant associations at *P* < .01% (FWE-corrected) (*N*=12,675). Coordinates of peak association for each cluster are reported in parentheses (in mm). Standard errors denote uncorrected 95% confidence intervals. See Supplementary Table 4 for more details.

**Supplementary Figure 4.**
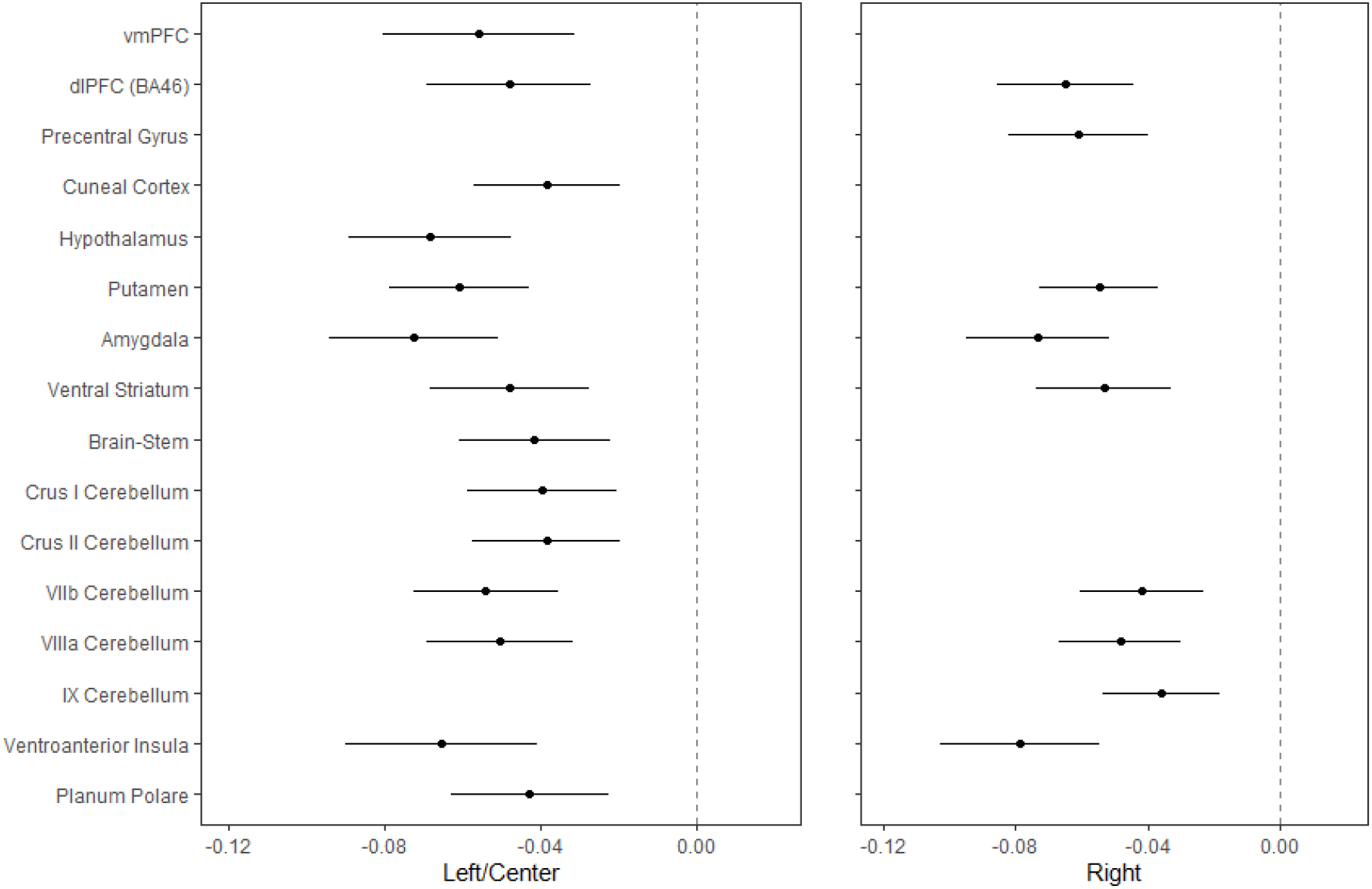
Effect sizes (standardized betas) of association between risky behaviour and IDPs of grey matter volume (GMV) showing significant associations at *P*<0.01 level (FWE-corrected) (*N*=12,675). Standard errors denote uncorrected 95% confidence intervals.

**Supplementary Figure 5.**
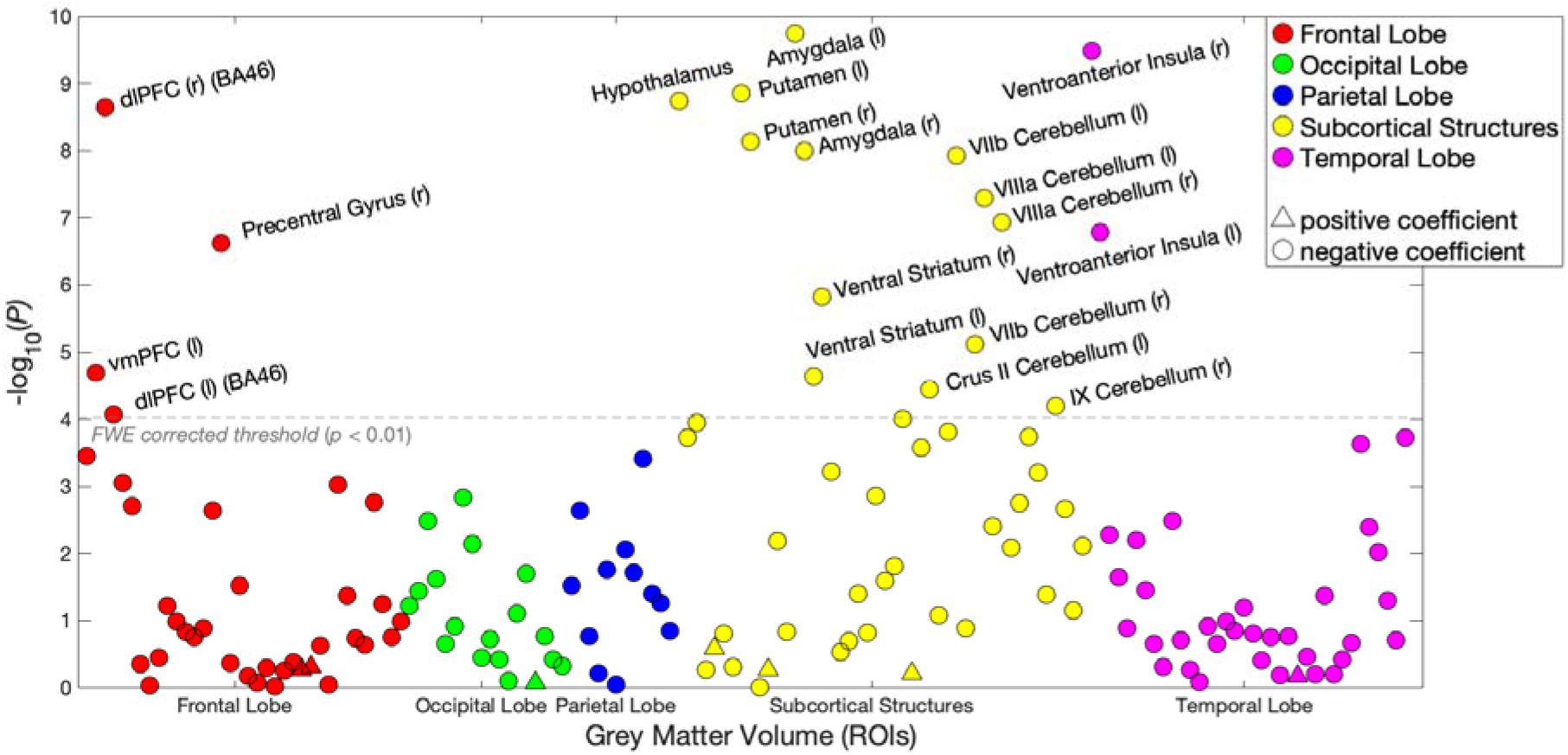
Associations (p-values) between risky behaviour and 148 ROI-level imaging-derived phenotypes (IDPs) of grey matter volume (GMV), controlling for cognitive and socioeconomic outcomes (*N* = 11,864). Control variables include *education years, fluid IQ, zip-code level social deprivation, household income, number of household members, birth location*, and all standard controls.

**Supplementary Figure 6.**
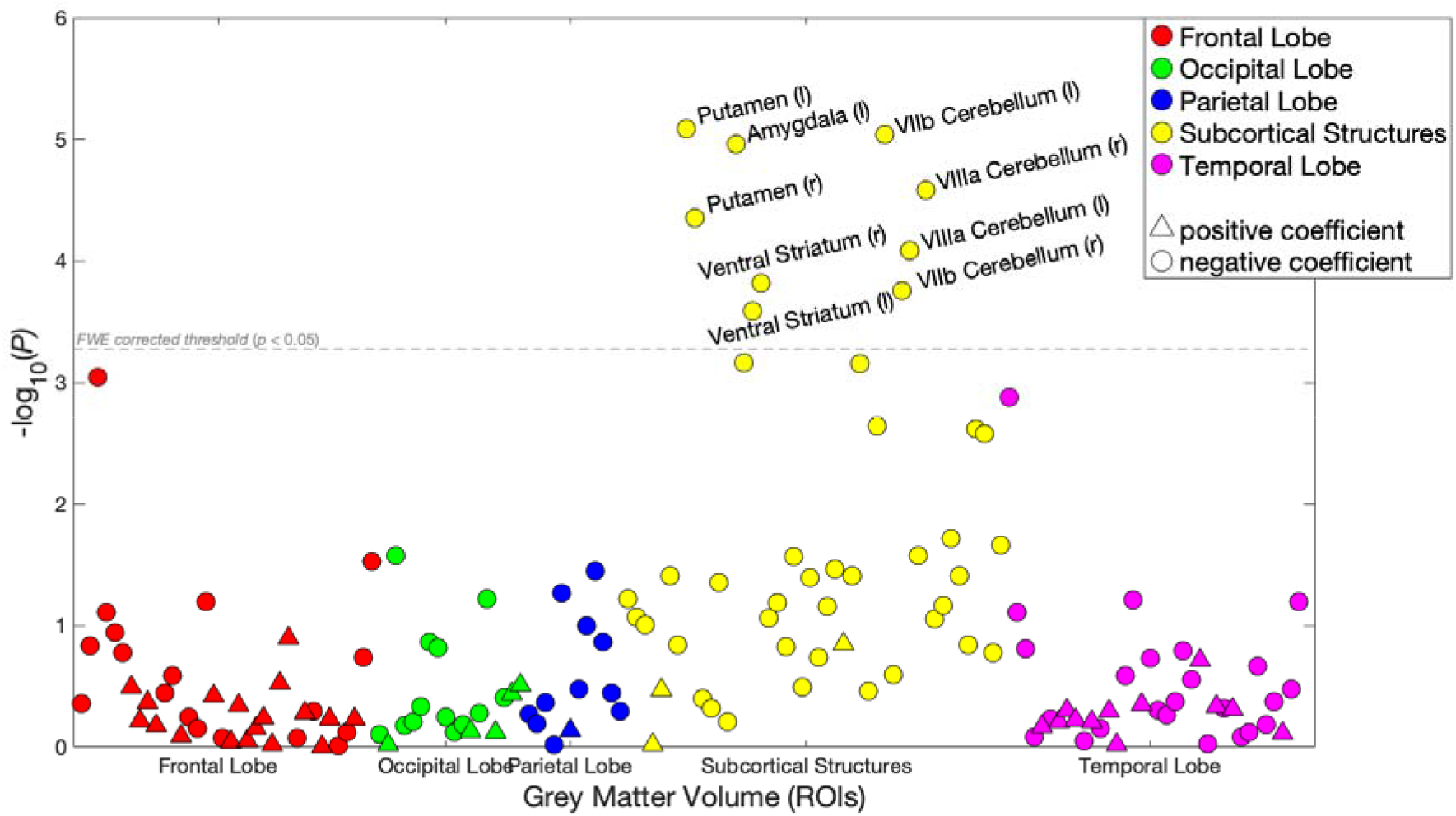
Associations (*p*-values) between risky behaviour and 148 ROI-level imaging-derived phenotypes (IDPs) of grey matter volume (GMV), controlling for current drinking levels (binned in deciles) and smoking levels (binned in 3 categories) in addition to all standard controls (N = 12,675).

**Supplementary Figure 7.**
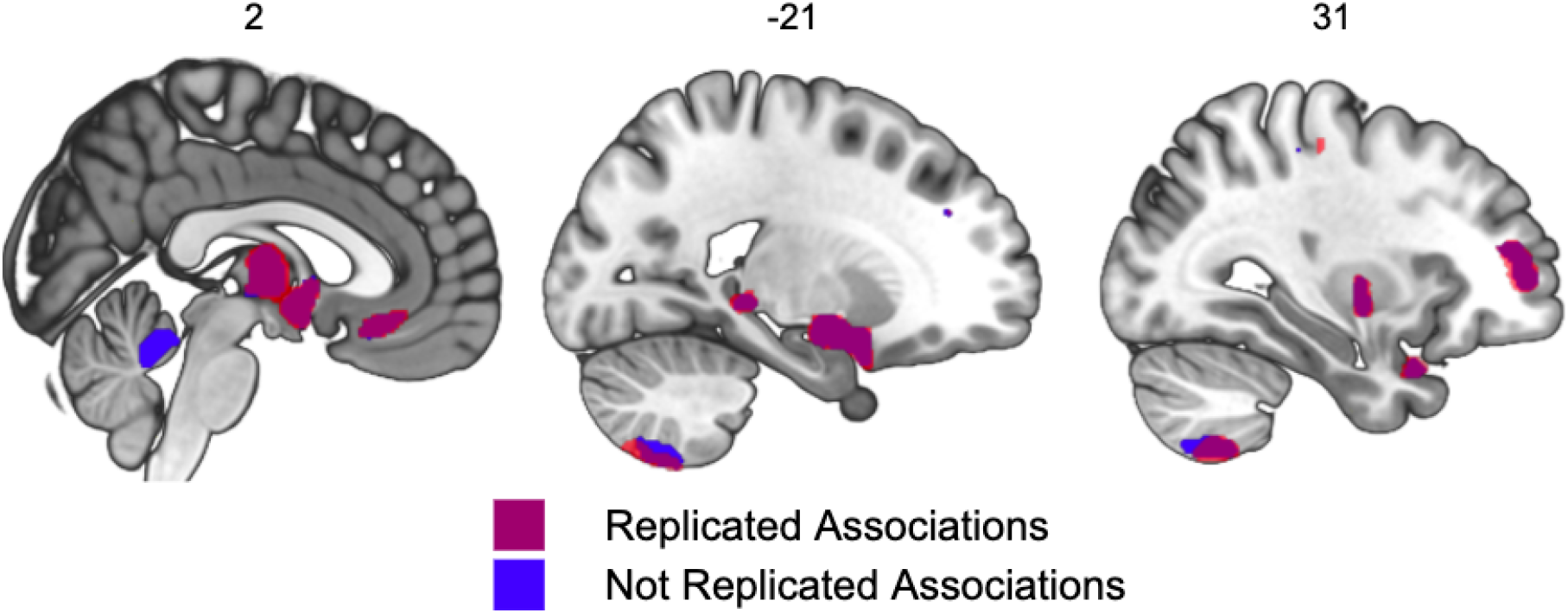
Voxel-level GMV associated with risky behaviour in the replication sample (*N* = 13,004). 92.6% of the voxels identified in our original analysis (located in 20 out of 21 original clusters identified, marked in purple) successfully replicate (corrected for multiple testing at the 5% level using a permutation test). Non replicated voxels are located in the cerebellar lobules I-IV (marked in blue).

**Supplementary Figure 8.**
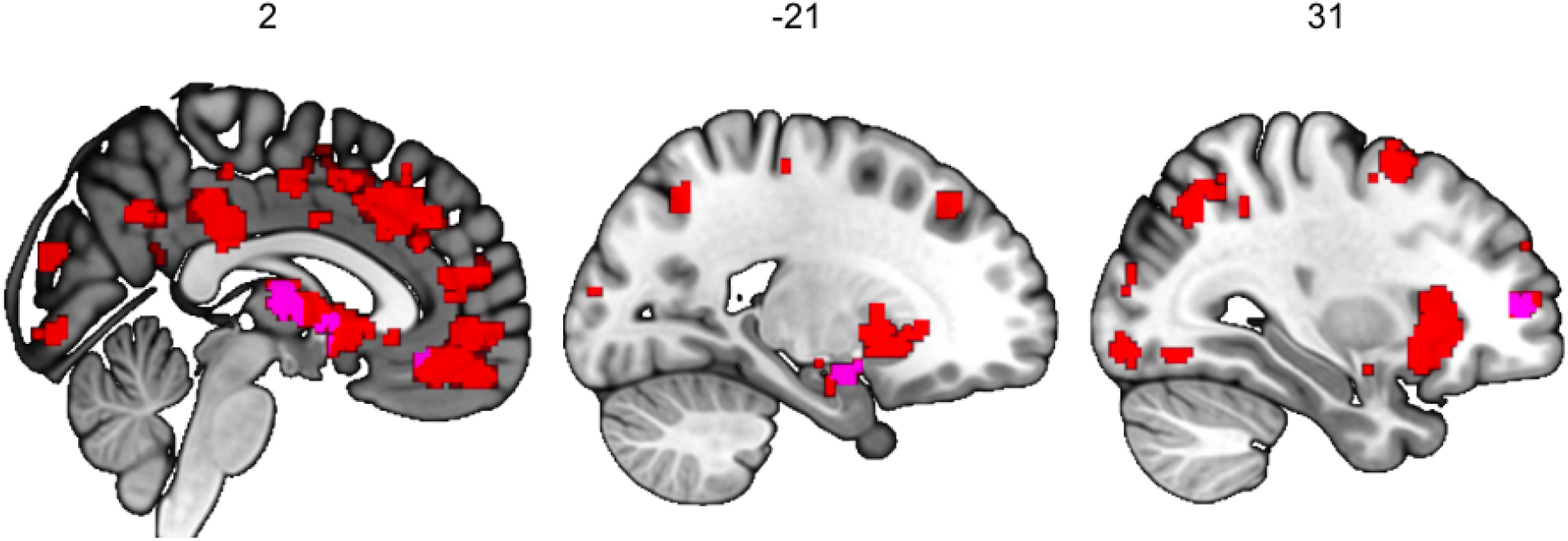
Meta analysis of functional MRI studies of risky behaviours, provided by Neurosynth (*N* = 4,717 participants and *K* = 101 studies. Conjunction with areas showing negative GMV association with risky behaviour (including thalamus, vmPFC, amygdala and dlPFC) is marked in magenta (see Supplementary Table 9). Additional meta-analytic functional activation areas are marked in red.

**Supplementary Figure 9.**
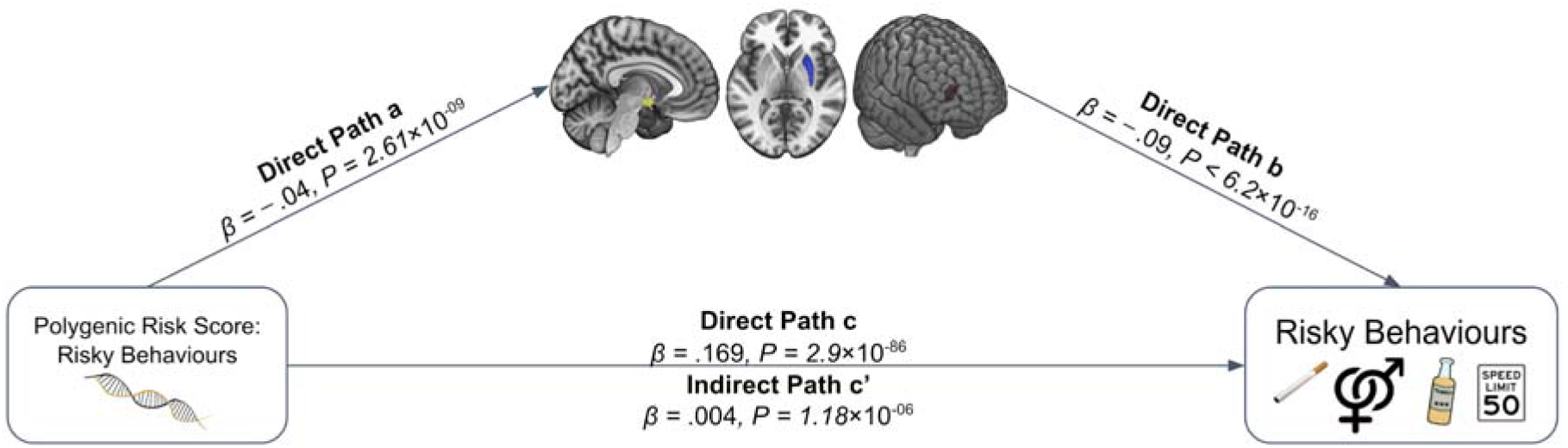
Meditation analysis of the association between PRS and risky behaviour with GMV in dlPFC, putamen and hypothalamus. The sum of all GMV differences in right dlPFC, putamen and hypothalamus (based on the activation masks from Fig 2A) mediated ~2.07% of the association between the PRS and risky behaviour. Arrows depict the direction of the structural equation modelling and do not imply causality (*N*=12,675).

## Supplementary Tables

**Supplementary Table 1.** Eigenvalues of the four Principal Components of Risky behaviours.

| Component | Eigenvalue | Cumulative Variance Explained |
| --- | --- | --- |
| Component 1 | 1.474 | 0.3685 |
| Component 2 | 0.975 | 0.6121 |
| Component 3 | 0.819 | 0.817 |
| Component 4 | 0.732 | 1 |

**Supplementary Table 2.** Eigenvectors of the four Principal Components of Risky behaviours.

| Variable | Comp1 | Comp2 | Comp3 | Comp4 |
| --- | --- | --- | --- | --- |
| Speeding | 0.4096 | 0.7643 | 0.1402 | 0.4779 |
| Sexual behaviour | 0.5543 | 0.1448 | -0.6327 | -0.5211 |
| Drinking | 0.5351 | -0.1876 | 0.7339 | -0.374 |
| Smoking | 0.4885 | -0.5998 | -0.2036 | 0.6002 |

**Supplementary Table 3.**
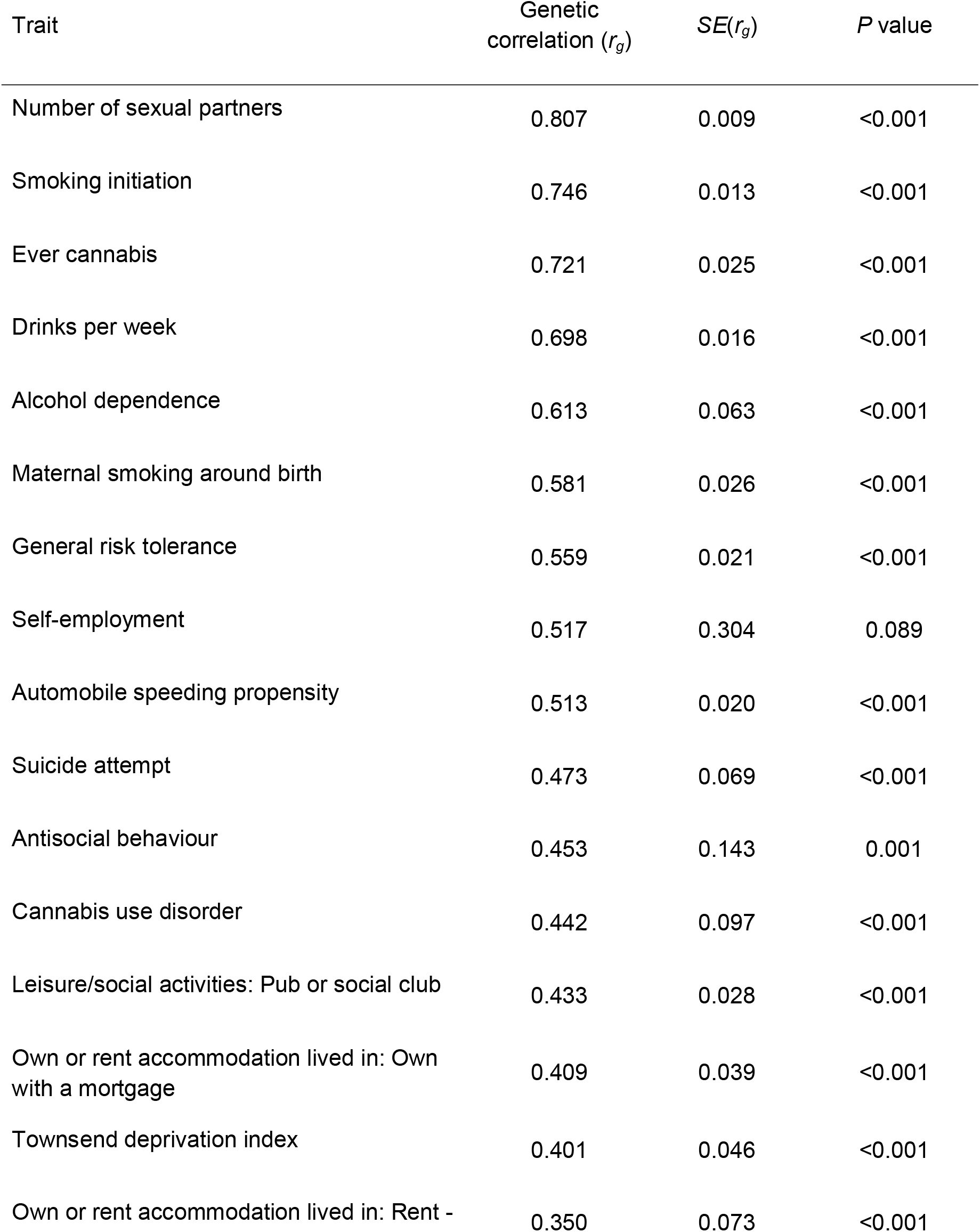
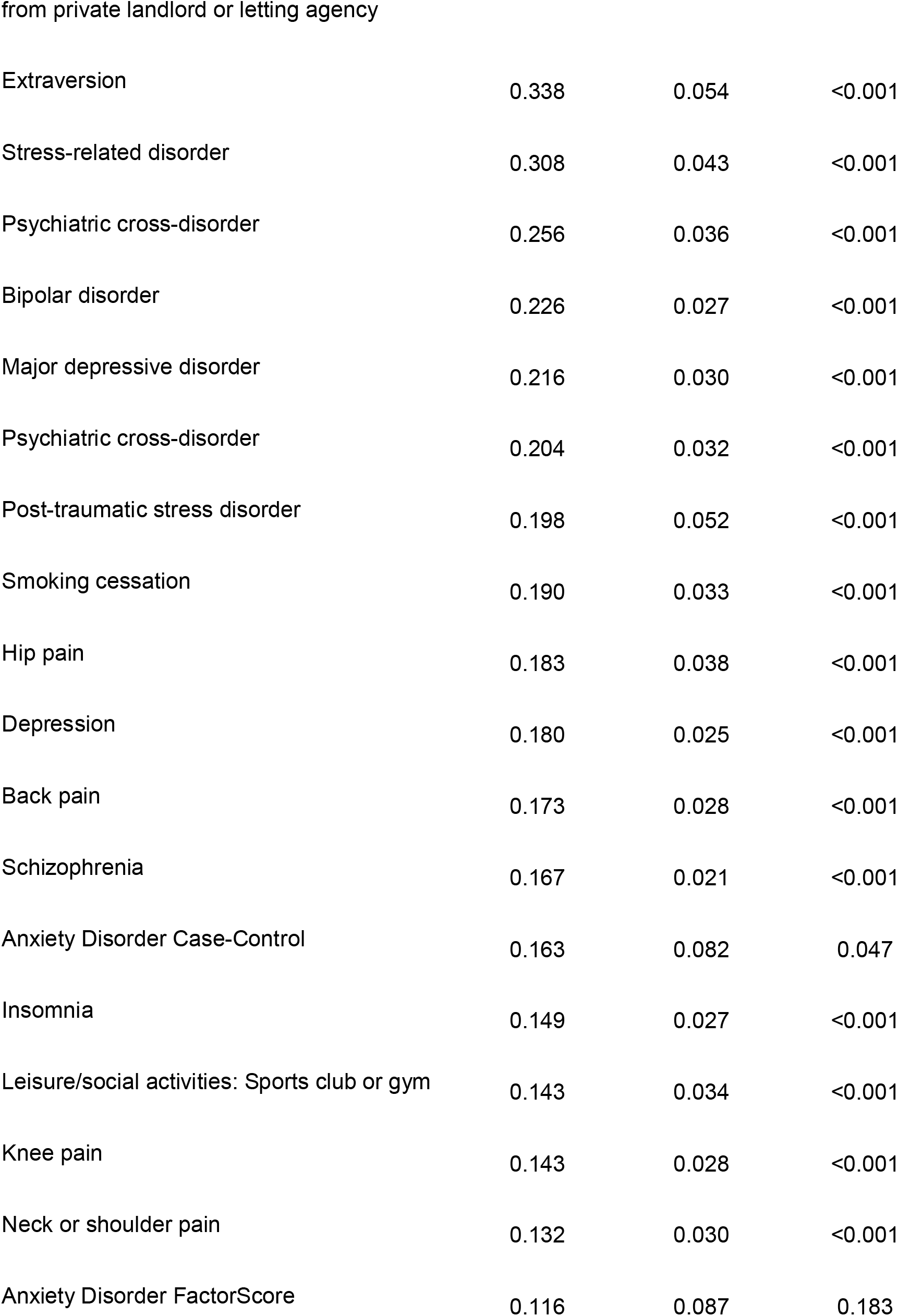
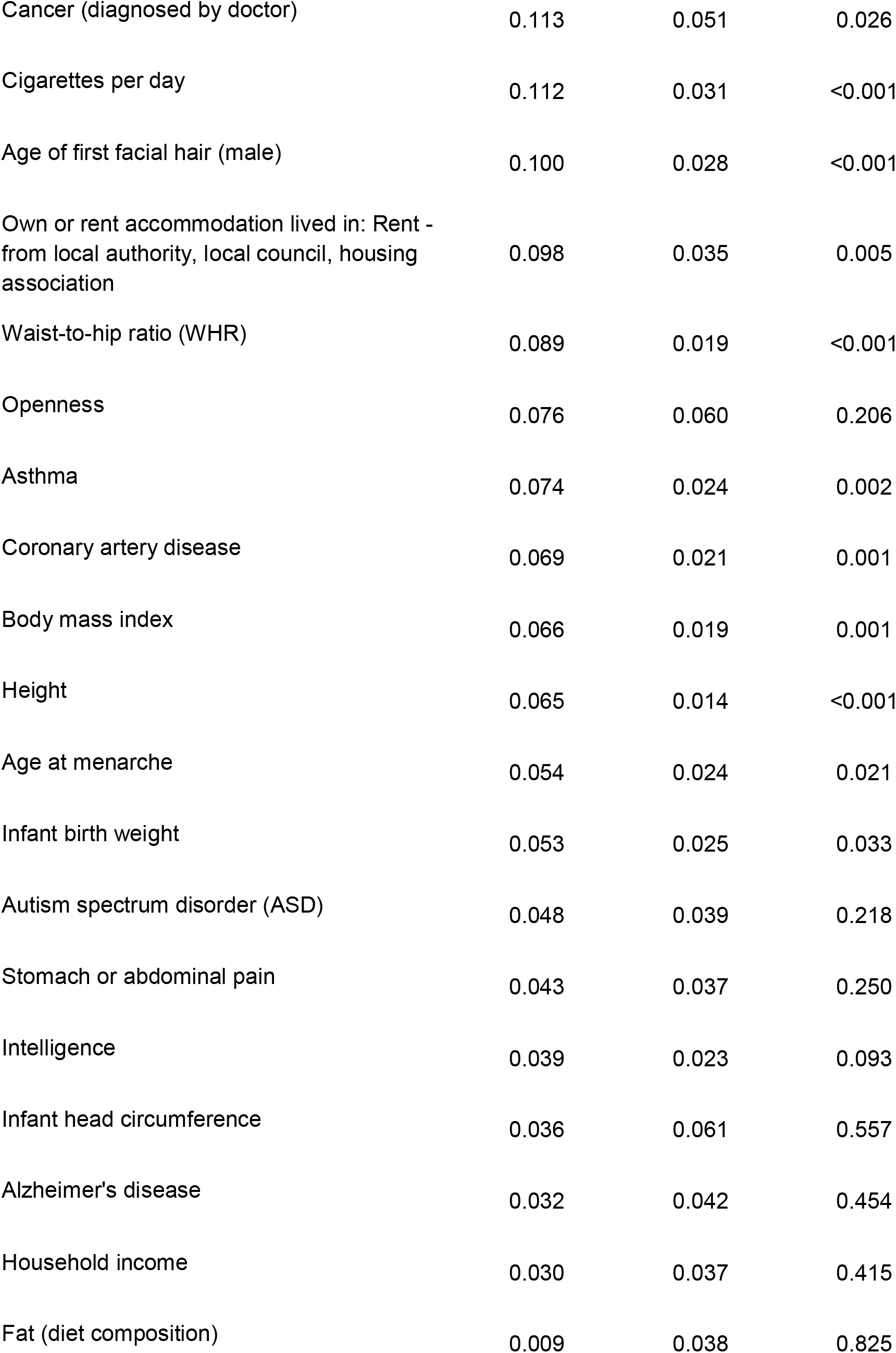
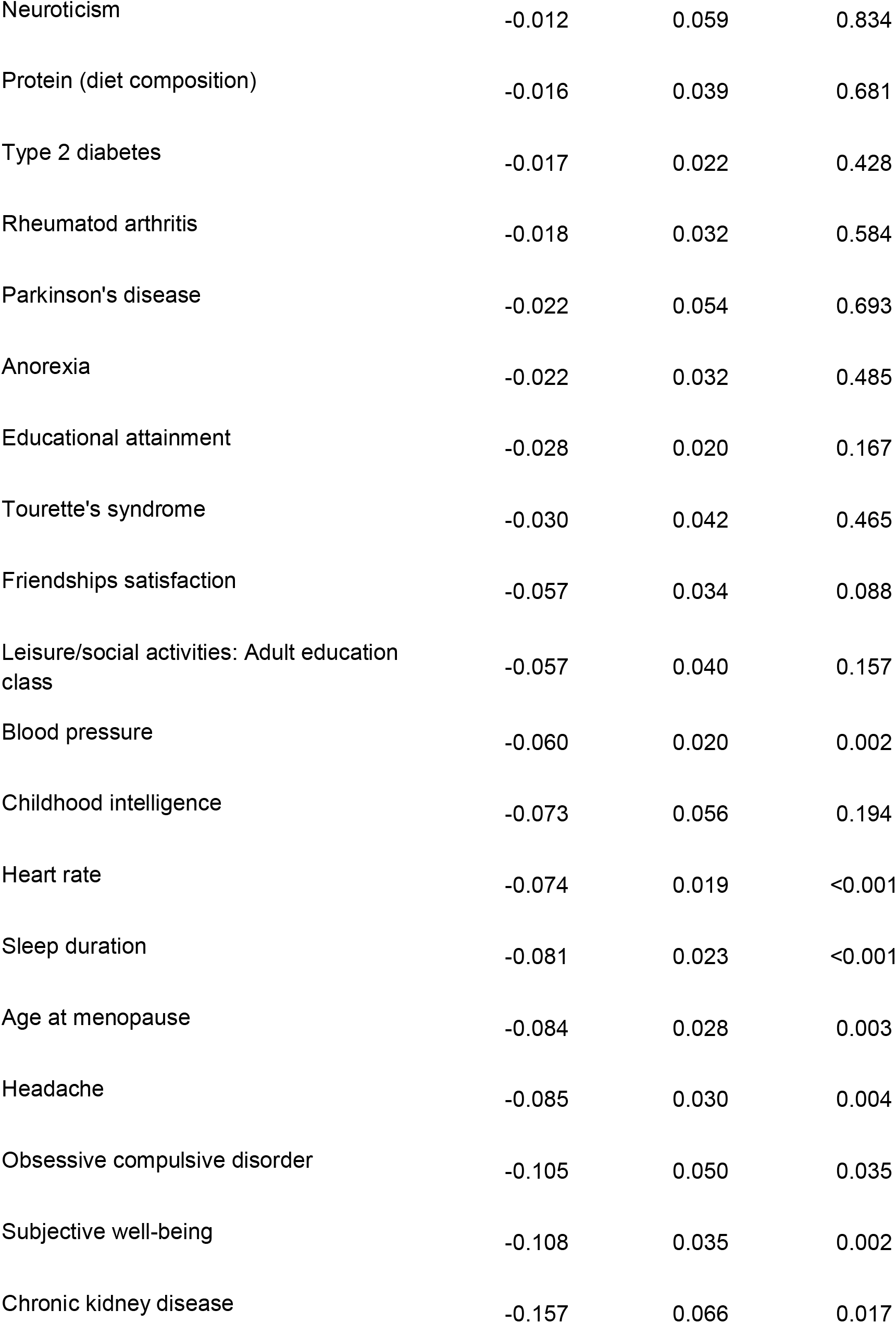
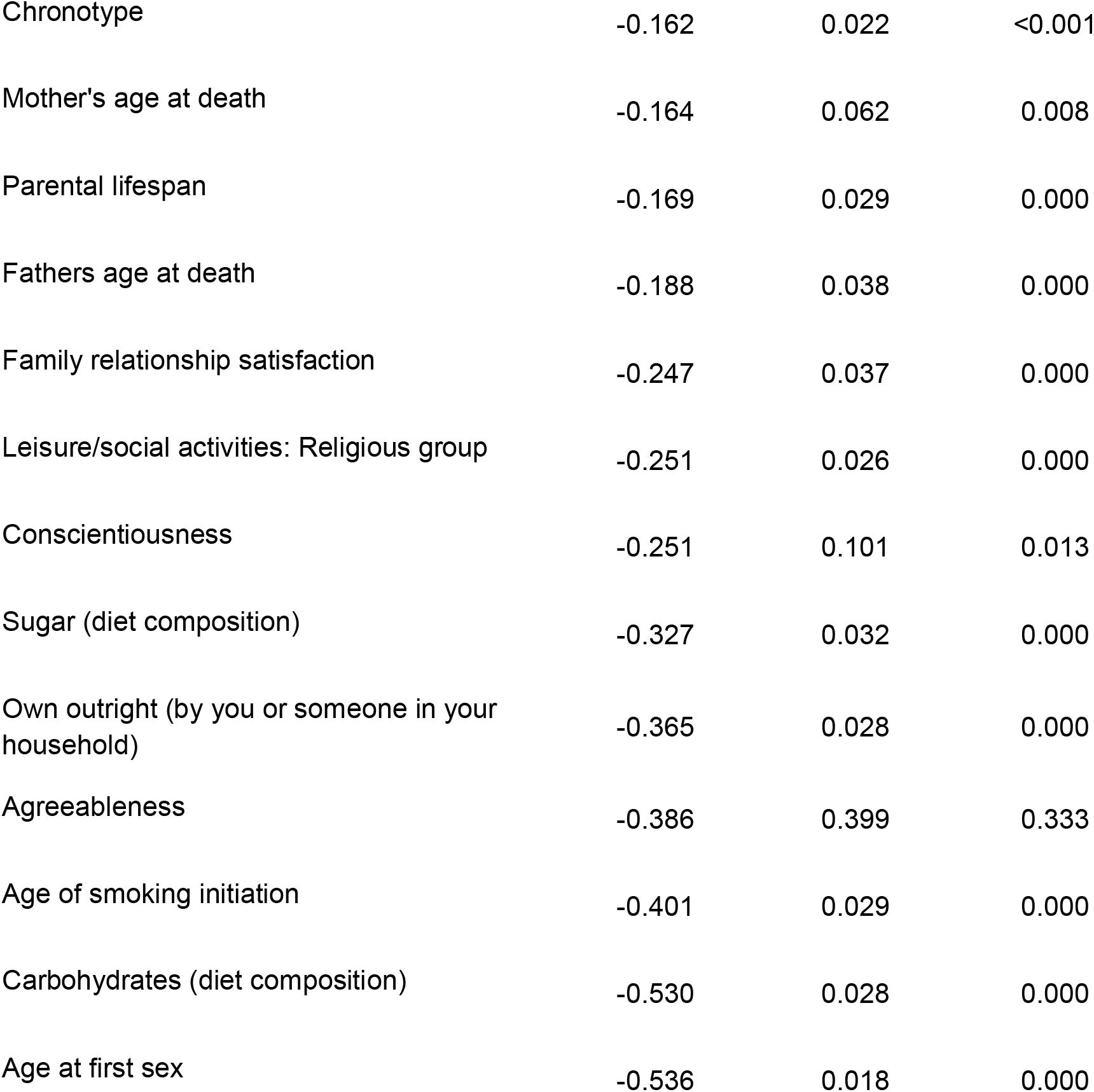
Genetic correlations (*r_g_*) between risky behaviour (GWAS *N* = 297,025) and 85 traits, estimated using bivariate LD Score regression^25^. All *P* values are based on two-sided statistical tests.

**Supplementary Table 4.** Association between risky behaviour and grey matter volumes (GMV) in clusters of voxels (*N*=12,675). Depicted are the summarized regression statistics per cluster. Δ*R*^2^ indicates the marginal increase in variance explained compared to a model that excludes GMV from the respective cluster. The corresponding coordinates of the peak activation in each cluster can be found in Supplementary Figure 3.

| Cluster | $\beta$ | SE | $P_{uncorr}$ | T | $R^2$ | $\Delta R^2$ |
| --- | --- | --- | --- | --- | --- | --- |
| 1 | -0.1154 | 0.012697 | $1.15 \times 10^{-19}$ | -9.0888 | 0.048208 | 0.006308 |
| 2 | -0.10848 | 0.012657 | $1.15 \times 10^{-17}$ | -8.5707 | 0.047519 | 0.005619 |
| 3 | -0.063596 | 0.00956 | $3.01 \times 10^{-11}$ | -6.6522 | 0.045312 | 0.003412 |
| 4 | -0.061123 | 0.009608 | $2.06 \times 10^{-10}$ | -6.3618 | 0.045026 | 0.003126 |
| 5 | -0.077548 | 0.010659 | $3.66 \times 10^{-13}$ | -7.2753 | 0.045969 | 0.004069 |
| 6 | -0.071566 | 0.010686 | $2.21 \times 10^{-11}$ | -6.6974 | 0.045358 | 0.003458 |
| 7 | -0.054051 | 0.009533 | $1.46 \times 10^{-08}$ | -5.6697 | 0.044394 | 0.002494 |
| 8 | -0.065617 | 0.011953 | $4.11 \times 10^{-08}$ | -5.4895 | 0.044242 | 0.002342 |
| 9 | -0.065851 | 0.010695 | $7.64 \times 10^{-10}$ | -6.1571 | 0.044832 | 0.002932 |
| 10 | -0.064221 | 0.010776 | $2.6 \times 10^{-09}$ | -5.9596 | 0.04465 | 0.00275 |
| 11 | -0.066171 | 0.011275 | $4.50 \times 10^{-09}$ | -5.8686 | 0.044569 | 0.002669 |
| 12 | -0.060586 | 0.010544 | $9.33 \times 10^{-09}$ | -5.7463 | 0.044461 | 0.002561 |
| 13 | -0.056703 | 0.01038 | $4.77 \times 10^{-08}$ | -5.463 | 0.04422 | 0.00232 |
| 14 | -0.052643 | 0.009898 | $1.06 \times 10^{-07}$ | -5.3187 | 0.044102 | 0.002202 |
| 15 | -0.054197 | 0.010214 | $1.14 \times 10^{-07}$ | -5.3061 | 0.044092 | 0.002192 |
| 16 | -0.051553 | 0.010108 | $3.44 \times 10^{-07}$ | -5.1003 | 0.043929 | 0.002029 |
| 17 | -0.05134 | 0.010063 | $3.42 \times 10^{-07}$ | -5.1017 | 0.04393 | 0.00203 |
| **18** | -0.050687 | 0.009961 | $3.66 \times 10^{-07}$ | -5.0884 | 0.04392 | 0.00202 |
| **19** | -0.050047 | 0.010073 | $6.84 \times 10^{-07}$ | -4.9685 | 0.043828 | 0.001928 |
| **20** | -0.049129 | 0.009852 | $6.23 \times 10^{-07}$ | -4.9866 | 0.043842 | 0.001942 |
| **21** | -0.048736 | 0.010007 | $1.19 \times 10^{-06}$ | -4.8702 | 0.043755 | 0.001855 |

**Supplementary Table 5.** Effect sizes (standardized betas) and 95% confidence interval (uncorrected) of associations between risky behaviour and grey matter volumes (GMV) in 23 ROIs, with and without controlling for cognitive and socioeconomic outcomes (*N* = 11,864). Additional controls include education years, fluid IQ, zip-code level social deprivation, household income, number of household members, birth location. Both models include all standard controls. The sample size of the analysis with additional controls is reduced due to missing data for some variables. *FWE-rate of 5%; **FWE-rate of 1%.

| **ROI** | **Risky behaviour**<br>**(with additional controls)** | **Risky behaviour**<br>**(with standard controls)** |
| --- | --- | --- |
| **vmPFC (l)** | -0.055** [-0.081, -0.03] | -0.056** [-0.08,-0.031] |
| **dIPFC (r) (BA46)** | -0.065** [-0.087, -0.044] | -0.065** [-0.086,-0.044] |
| **dIPFC (l) (BA46)** | -0.044** [-0.066, -0.022] | -0.048** [-0.069,-0.027] |
| **Precentral Gyrus (r)** | -0.057** [-0.079, -0.036] | -0.061** [-0.082,-0.04] |
| **Cuneal Cortex (l)** | -0.031 [-0.051, -0.012] | -0.038** [-0.057,-0.02] |
| **Hypothalamus** | -0.066** [-0.087, -0.044] | -0.068** [-0.089,-0.047] |
| **Putamen (l)** | -0.057** [-0.076, -0.039] | -0.061** [-0.079,-0.043] |
| **Putamen (r)** | -0.055** [-0.073, -0.036] | -0.055** [-0.073,-0.037] |
| **Amygdala (l)** | -0.073** [-0.095, -0.051] | -0.072** [-0.094,-0.051] |
| **Amygdala (r)** | -0.066** [-0.089, -0.043] | -0.073** [-0.095,-0.051] |
| **Ventral Striatum (l)** | -0.045** [-0.066, -0.024] | -0.048** [-0.068,-0.027] |
| **Ventral Striatum (r)** | -0.052** [-0.072, -0.031] | -0.053** [-0.074,-0.033] |
| **Brain-Stem** | -0.035 [-0.055, -0.015] | -0.041** [-0.061,-0.022] |
| **Crus I Cerebellum (l)** | -0.039* [-0.059, -0.019] | -0.04**<br>[-0.059,-0.02] |
| **Crus II Cerebellum (l)** | -0.041** [-0.06, -0.021] | -0.038** [-0.057,-0.02] |
| **VIIb Cerebellum (l)** | -0.055** [-0.074, -0.036] | -0.054** [-0.072,-0.035] |
| **VIIb Cerebellum (r)** | -0.044** [-0.063, -0.025] | -0.042** [-0.06,-0.023] |
| **VIIIa Cerebellum (l)** | -0.053** [-0.072, -0.034] | -0.05** [-0.069,-0.032] |
| **VIIIa Cerebellum (r)** | -0.052** [-0.071, -0.032] | -0.048** [-0.067,-0.03] |
| **IX Cerebellum (r)** | -0.038** [-0.056, -0.019] | -0.036** [-0.054,-0.018] |
| **Ventroanterior Insula (r)** | -0.079** [-0.104, -0.055] | -0.079** [-0.103,-0.055] |
| **Ventroanterior Insula (l)** | -0.068** [-0.093, -0.042] | -0.065** [-0.09,-0.041] |
| **Planum Polare (l)** | -0.039* [-0.06, -0.018] | -0.043** [-0.063,-0.022] |

**Supplementary Table 6.**
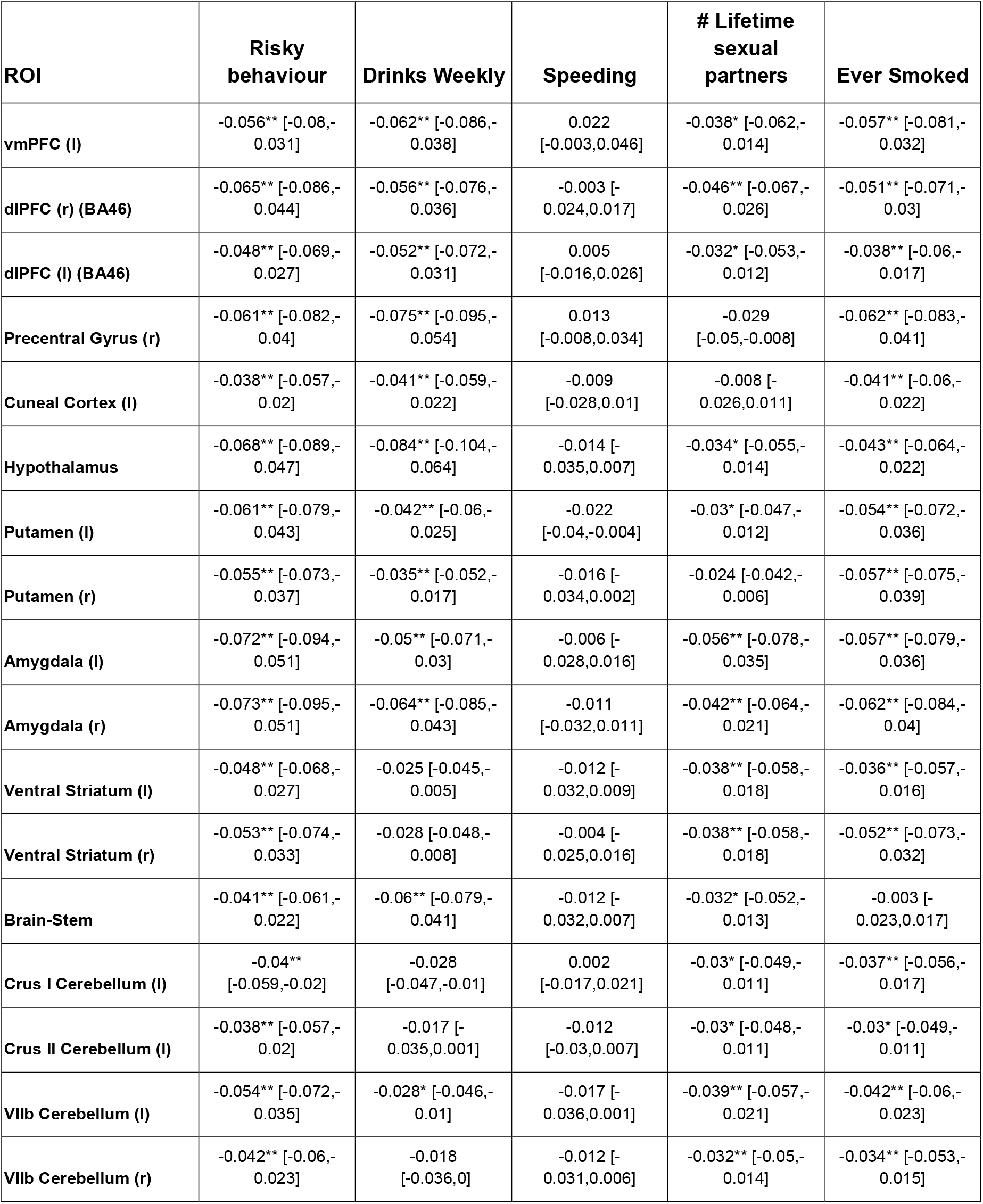
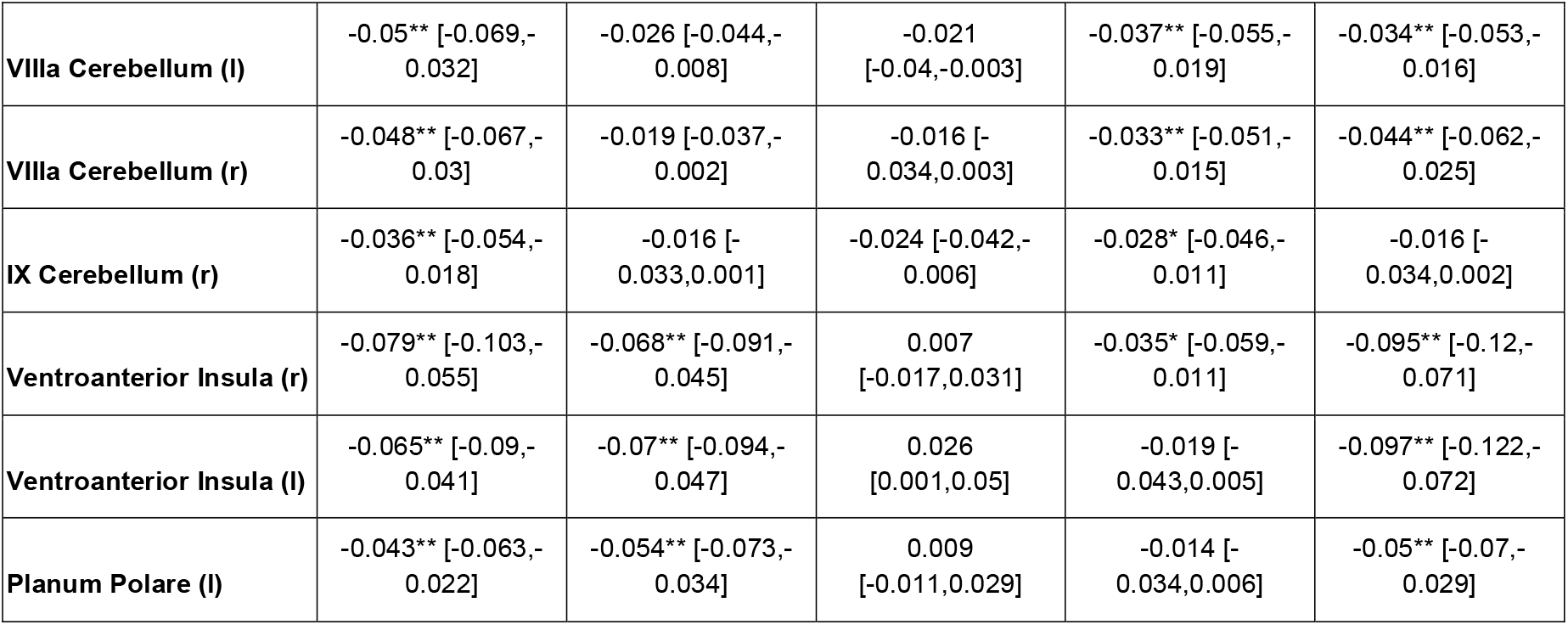
Effect sizes (standardized betas) and the corresponding 95% confidence interval (uncorrected) of the associations between grey matter volumes in 23 ROIs and individual phenotypes of risky behaviour (*N*=12,675). Models include all of the standard control variables. *FWE-rate of 5 %; **FWE-rate of 1%.

**Supplementary Table 7.** Effect sizes (standardized betas) and the corresponding 95% confidence intervals (uncorrected) of the associations between risky behaviour and grey matter volume in 23 ROIs, with and without controlling for current drinking level (binned in deciles) and current smoking level (binned in 3 categories). Both models include all of our standard controls (*N* = 12,675). *FWE-rate of 5 %; **FWE-rate of 1%.

| ROI | Risky behaviour<br>(with drinking/smoking<br>controls) | Risky behaviour<br>(with standard controls) |
| --- | --- | --- |
| vmPFC (l) | -0.015 [-0.036, 0.005] | -0.056** [-0.08,-0.031] |
| dIPFC (r) (BA46) | -0.029 [-0.047, -0.012] | -0.065** [-0.086,-0.044] |
| dIPFC (l) (BA46) | -0.016 [-0.033, 0.002] | -0.048** [-0.069,-0.027] |
| Precentral Gyrus (r) | -0.017 [-0.034, 0.001] | -0.061** [-0.082,-0.04] |
| Cuneal Cortex (l) | -0.012 [-0.028, 0.004] | -0.038** [-0.057,-0.02] |
| Hypothalamus | -0.017 [-0.034, 0.001] | -0.068** [-0.089,-0.047] |
| Putamen (l) | -0.034** [-0.049, -0.019] | -0.061** [-0.079,-0.043] |
| Putamen (r) | -0.031** [-0.046, -0.016] | -0.055** [-0.073,-0.037] |
| Amygdala (l) | -0.041** [-0.059, -0.023] | -0.072** [-0.094,-0.051] |
| Amygdala (r) | -0.032 [-0.05, -0.013] | -0.073** [-0.095,-0.051] |
| Ventral Striatum (l) | -0.032* [-0.049, -0.015] | -0.048** [-0.068,-0.027] |
| Ventral Striatum (r) | -0.033* [-0.05, -0.016] | -0.053** [-0.074,-0.033] |
| Brain-Stem | -0.014 [-0.031, 0.002] | -0.041** [-0.061,-0.022] |
| Crus I Cerebellum (l) | -0.017 [-0.033, -0.001] | -0.04**<br>[-0.059,-0.02] |
| Crus II Cerebellum (l) | -0.027 [-0.043, -0.011] | -0.038** [-0.057,-0.02] |
| VIIb Cerebellum (l) | -0.035** [-0.05, -0.02] | -0.054** [-0.072,-0.035] |
| VIIb Cerebellum (r) | -0.03* [-0.045, -0.014] | -0.042** [-0.06,-0.023] |
| VIIIa Cerebellum (l) | -0.031** [-0.047, -0.016] | -0.05** [-0.069,-0.032] |
| VIIIa Cerebellum (r) | -0.033** [-0.048, -0.018] | -0.048** [-0.067,-0.03] |
| **IX Cerebellum (r)** | -0.023 [-0.038, -0.008] | -0.036** [-0.054,-0.018] |
| **Ventroanterior Insula (r)** | -0.033 [-0.053, -0.013] | -0.079** [-0.103,-0.055] |
| **Ventroanterior Insula (l)** | -0.018 [-0.039, 0.002] | -0.065** [-0.09,-0.041] |
| **Planum Polare (l)** | -0.011 [-0.028, 0.006] | -0.043** [-0.063,-0.022] |

**Supplementary Table 8.**
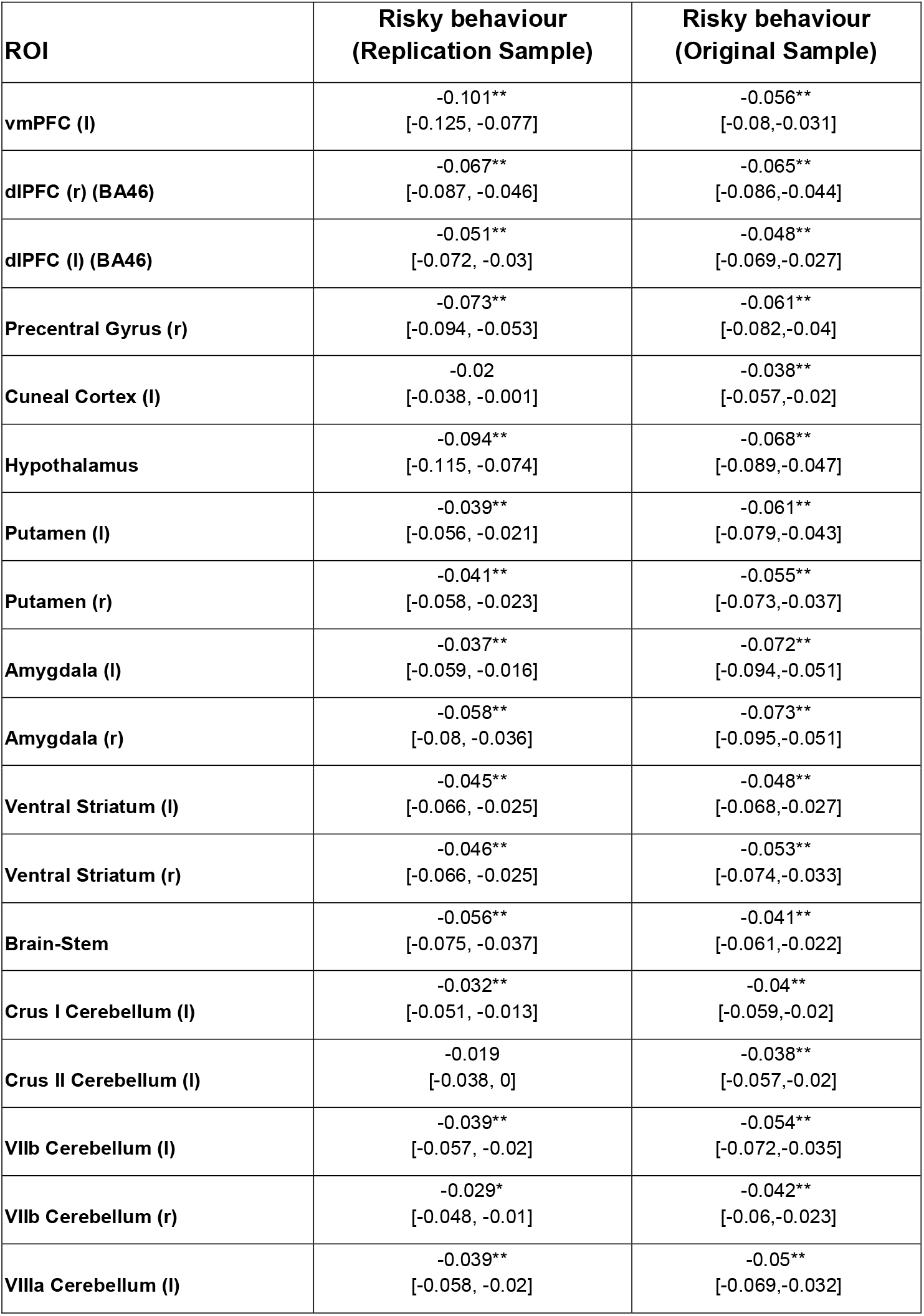
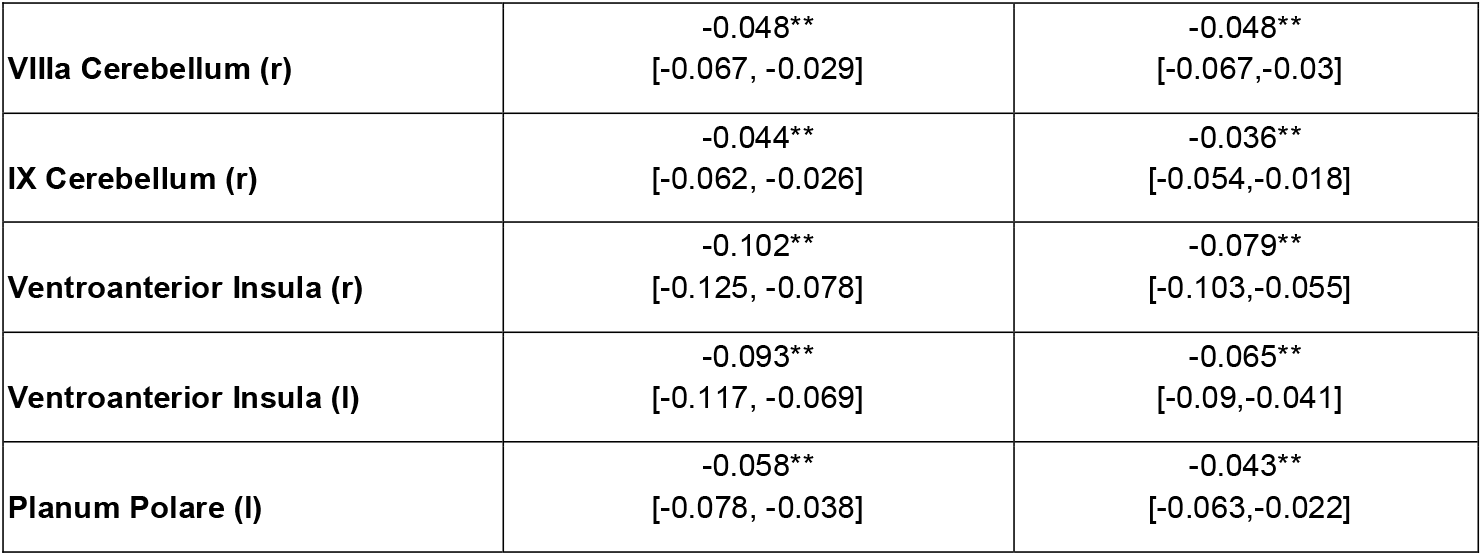
Effect sizes (standardized betas) and the corresponding 95% confidence intervals (uncorrected) of the associations between risky behaviour and ROI-level imaging-derived phenotypes (IDPs) of grey matter volume (GMV) in the replication sample (*N* = 13,004) and original sample (*N*=12,675). All beta coefficients are consistently negative across samples and 21 of 23 ROIs identified in the original analysis replicate (corrected for multiple testing using a permutation test, see methods). *FWE-rate of 5%; **FWE-rate of 1%.

**Supplementary Table 9.**
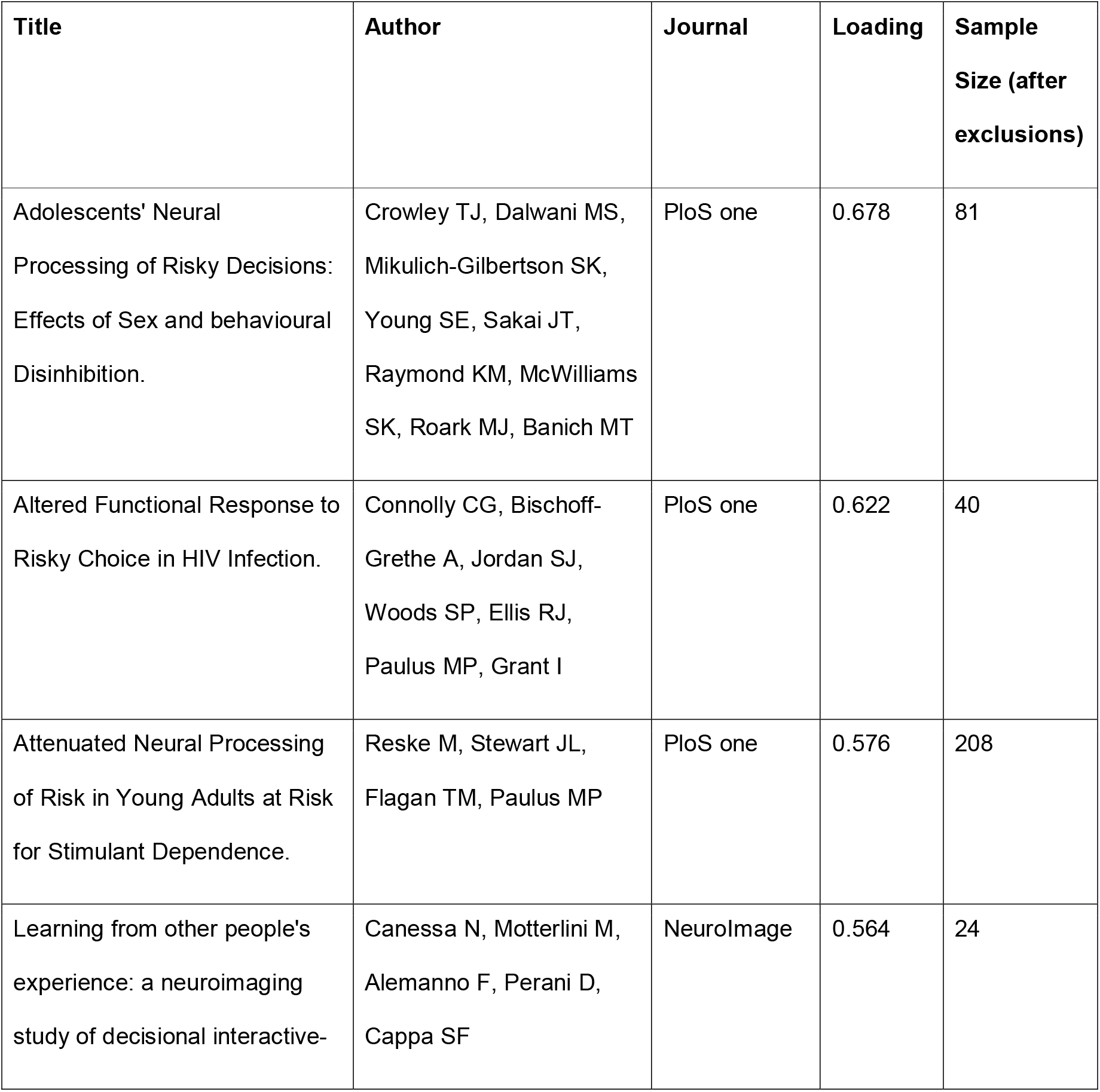
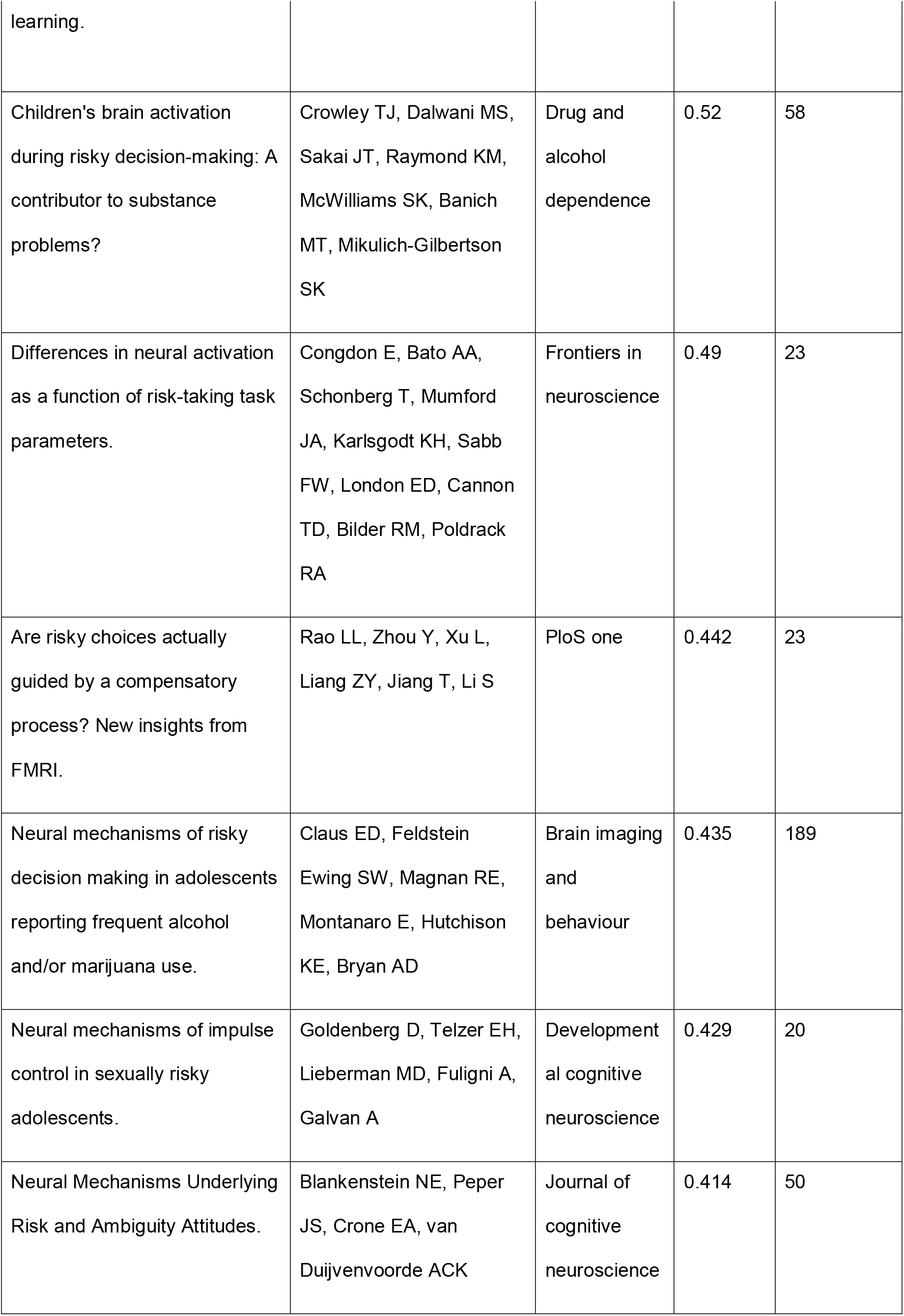
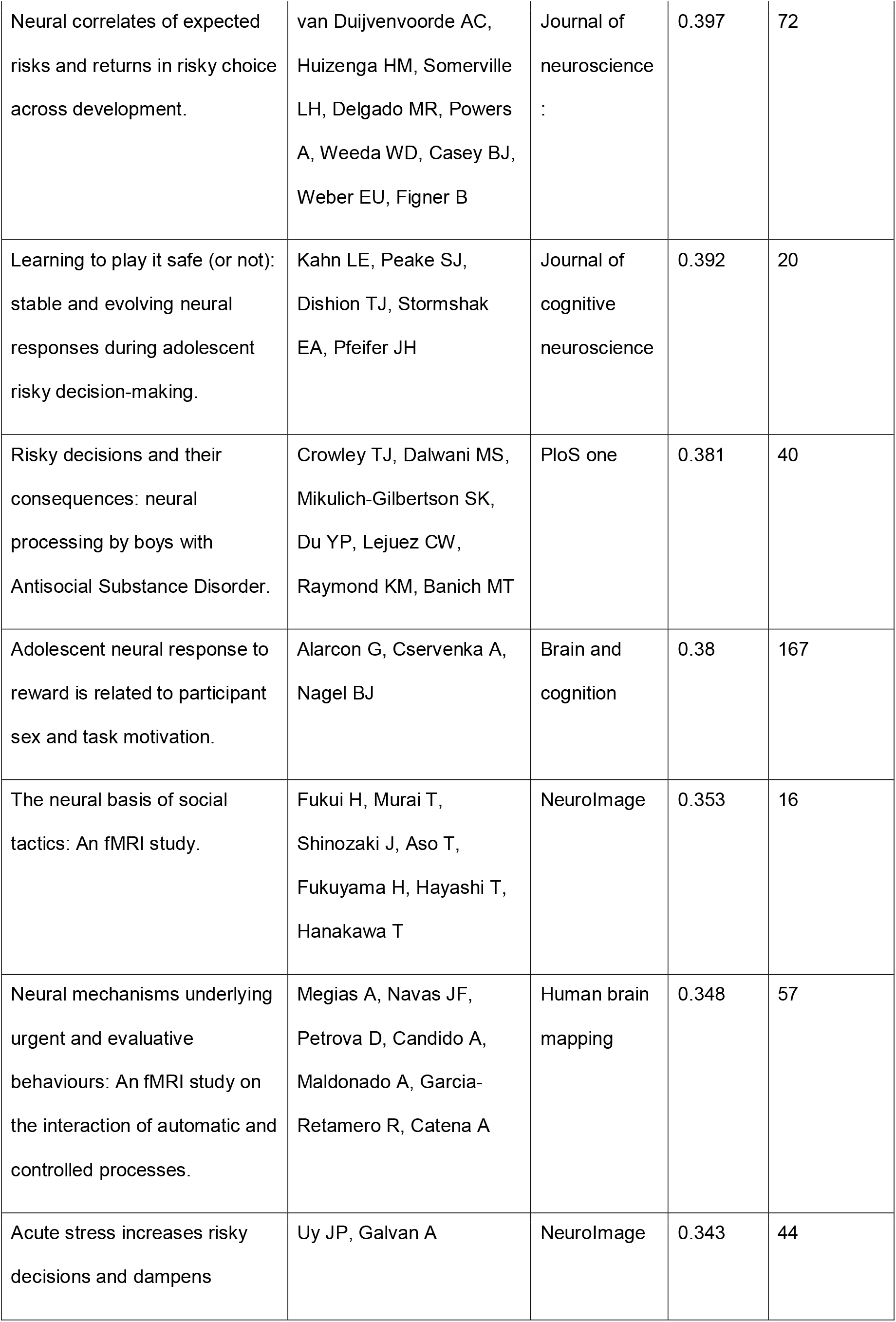
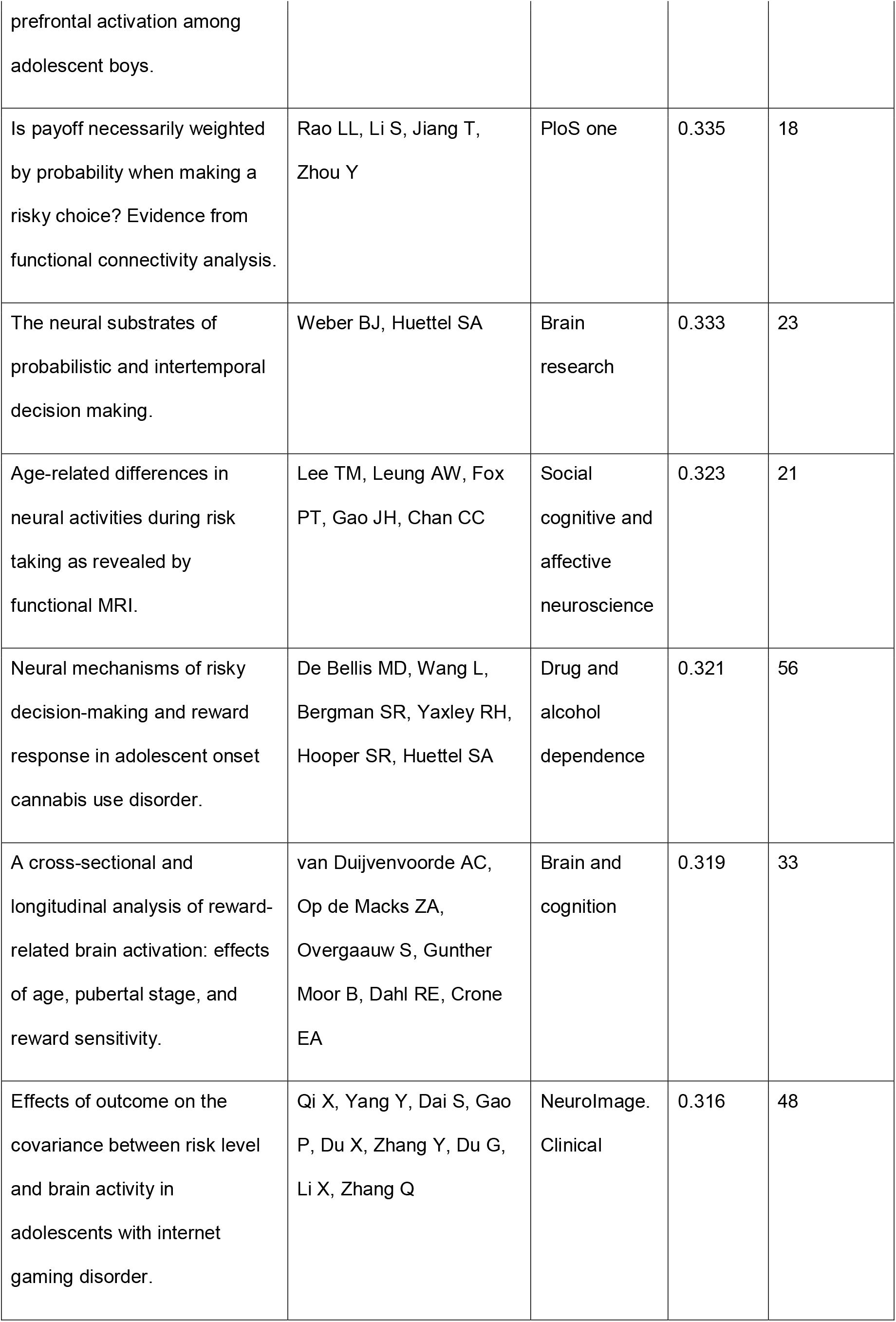
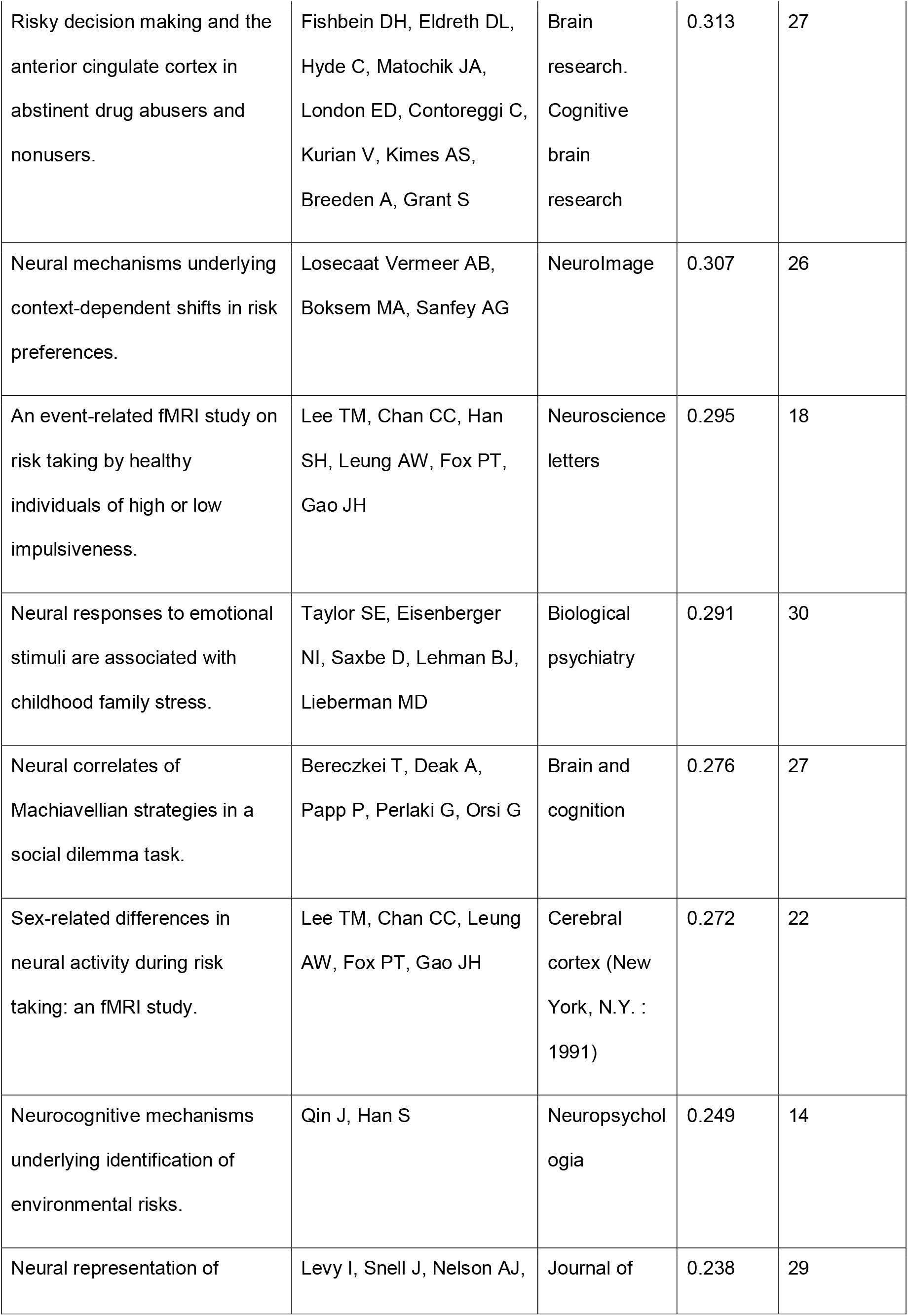
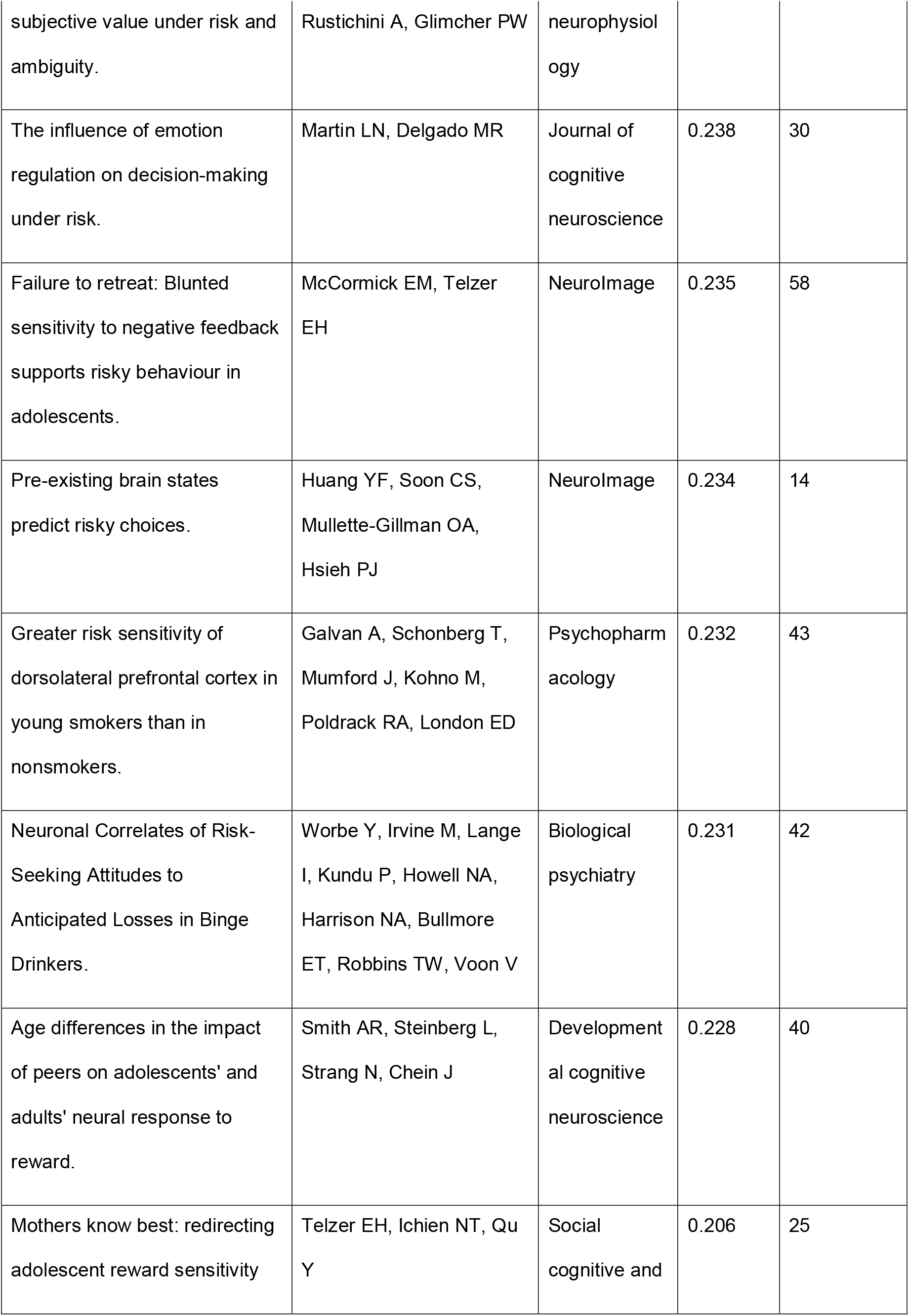
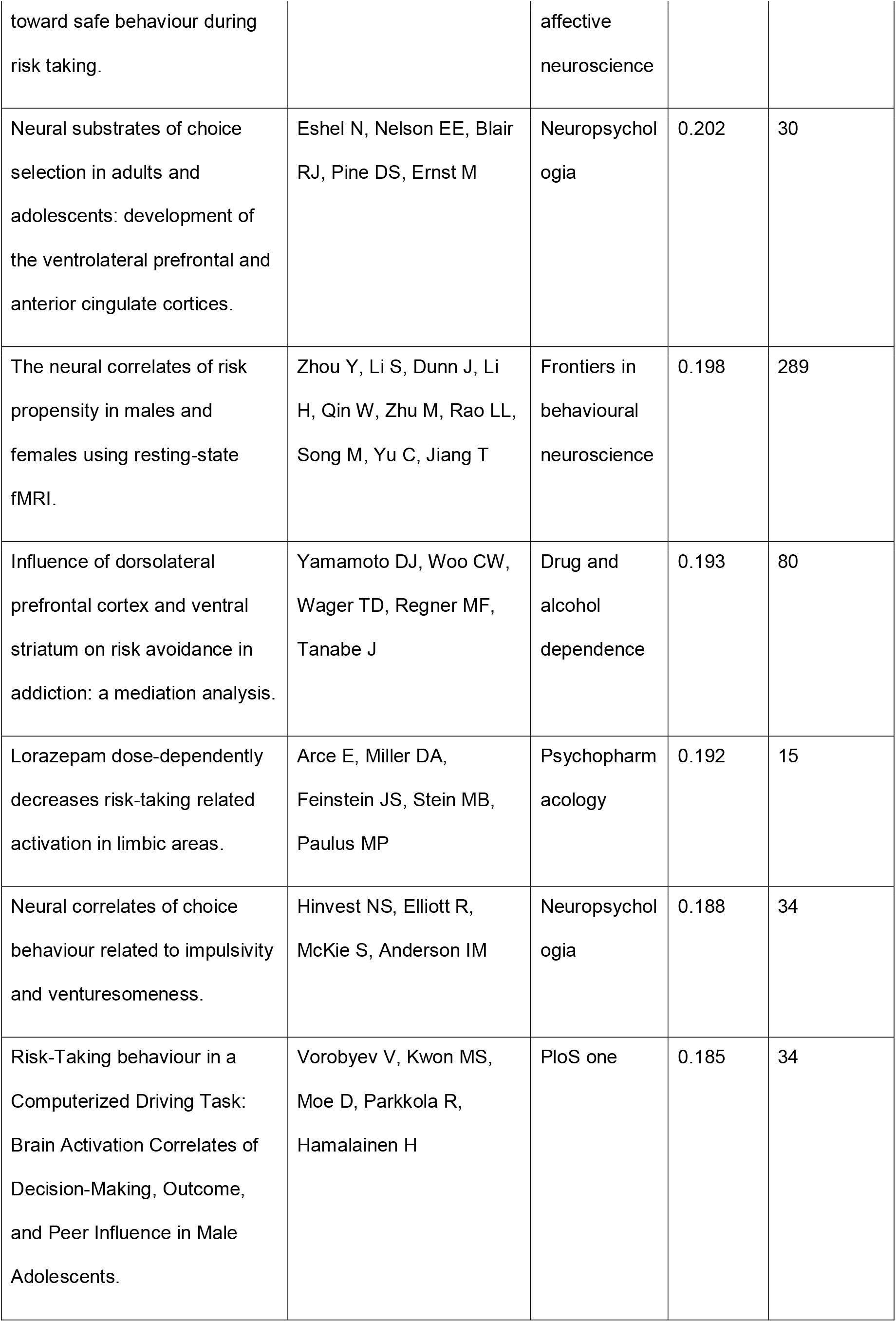
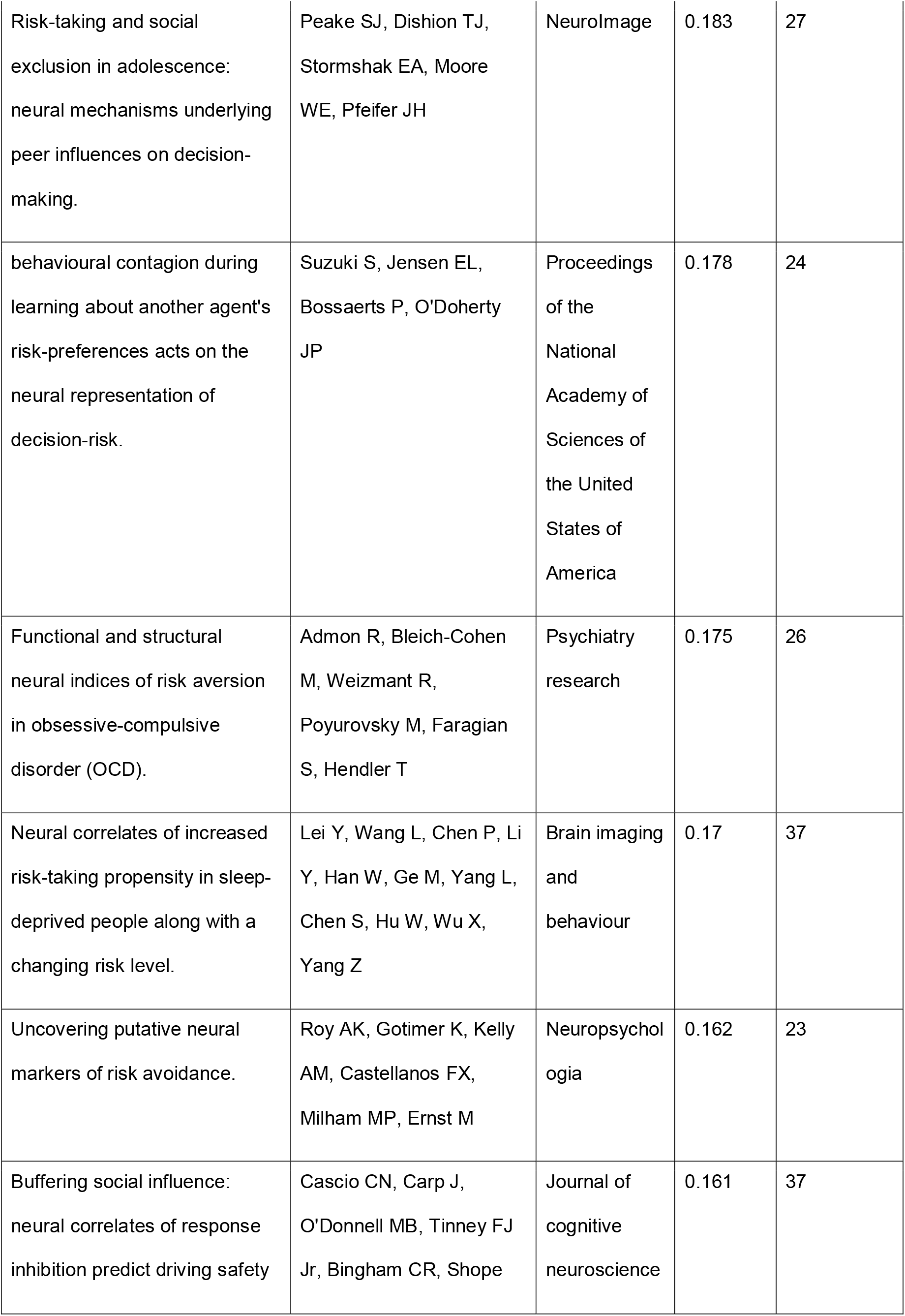
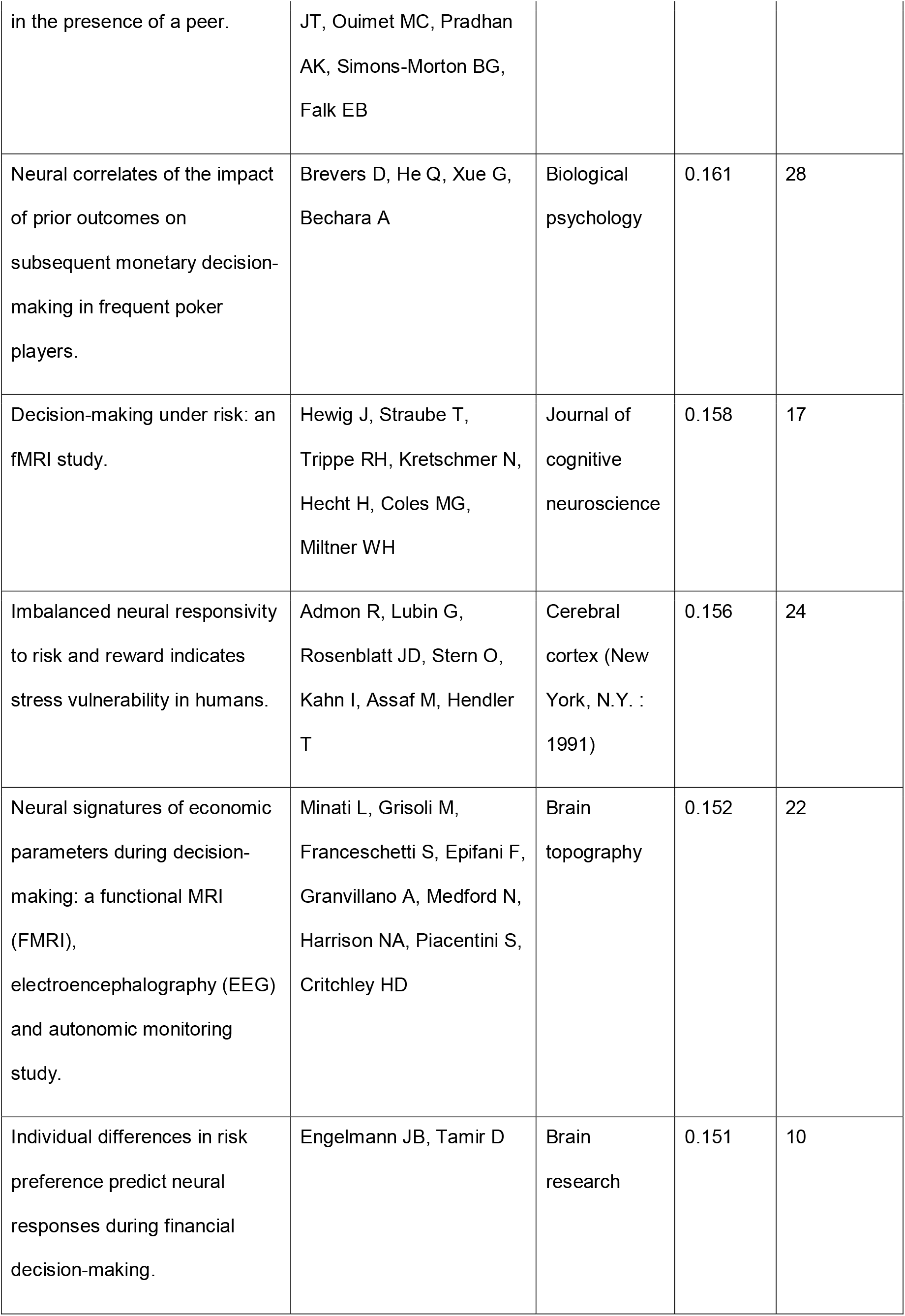
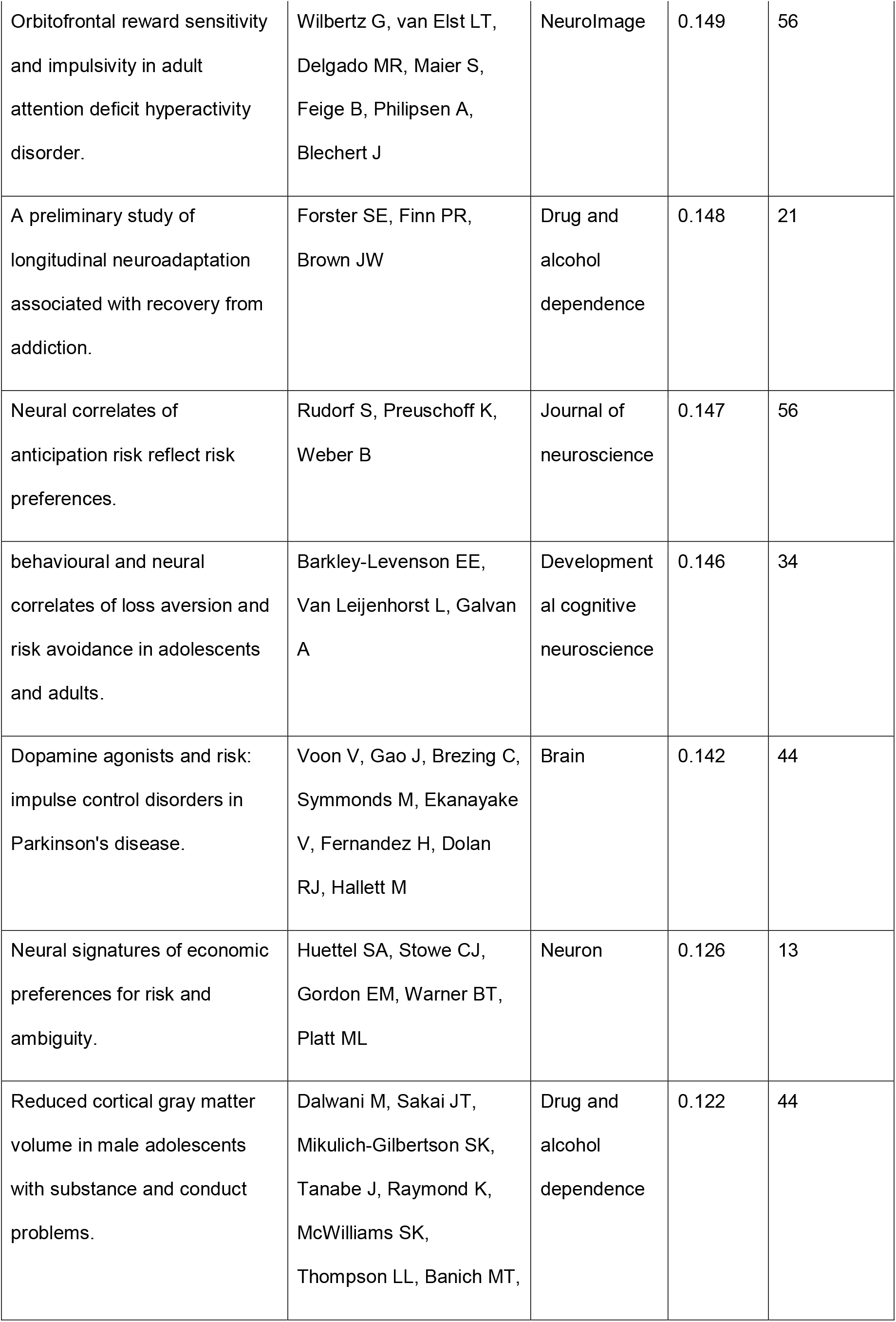
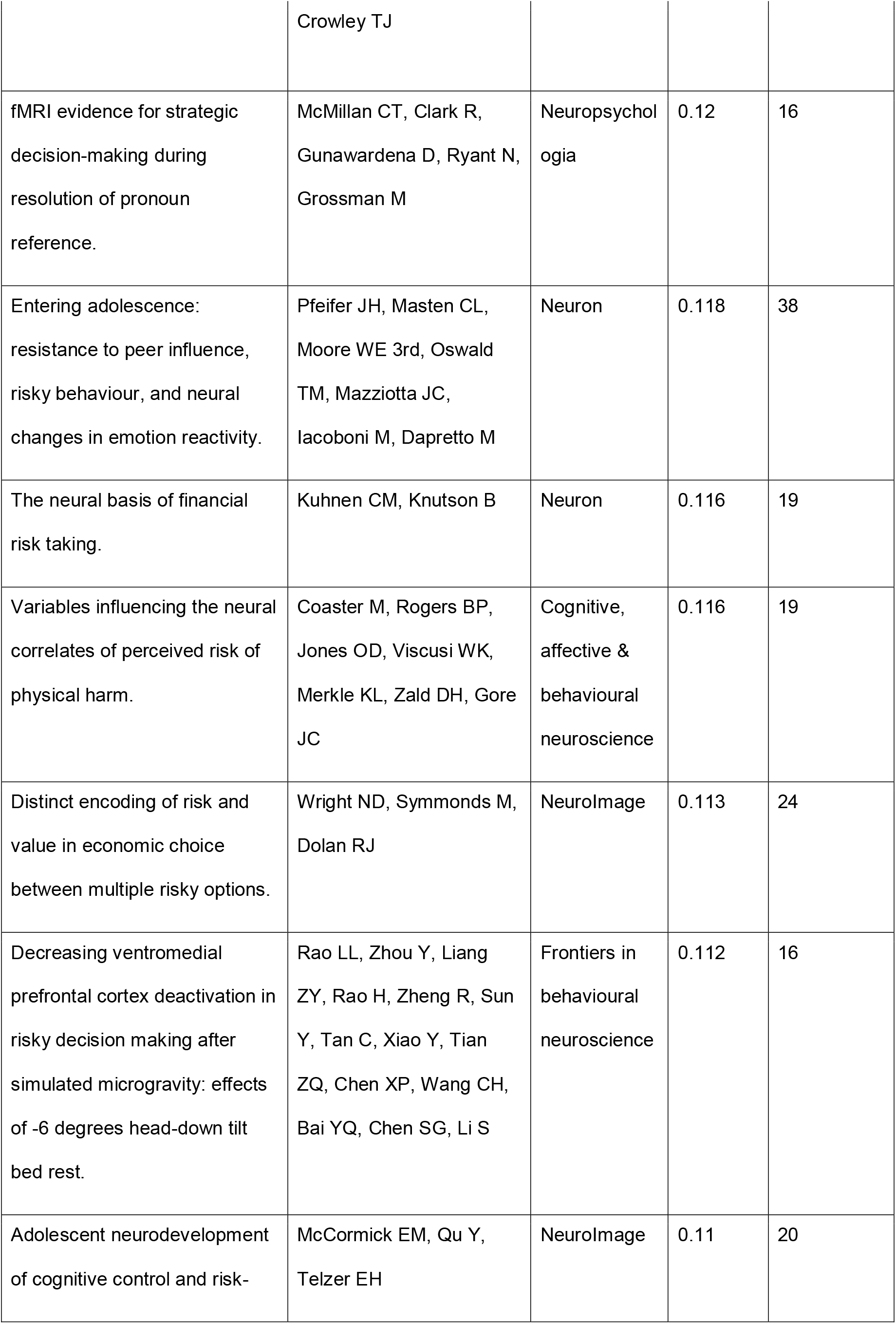
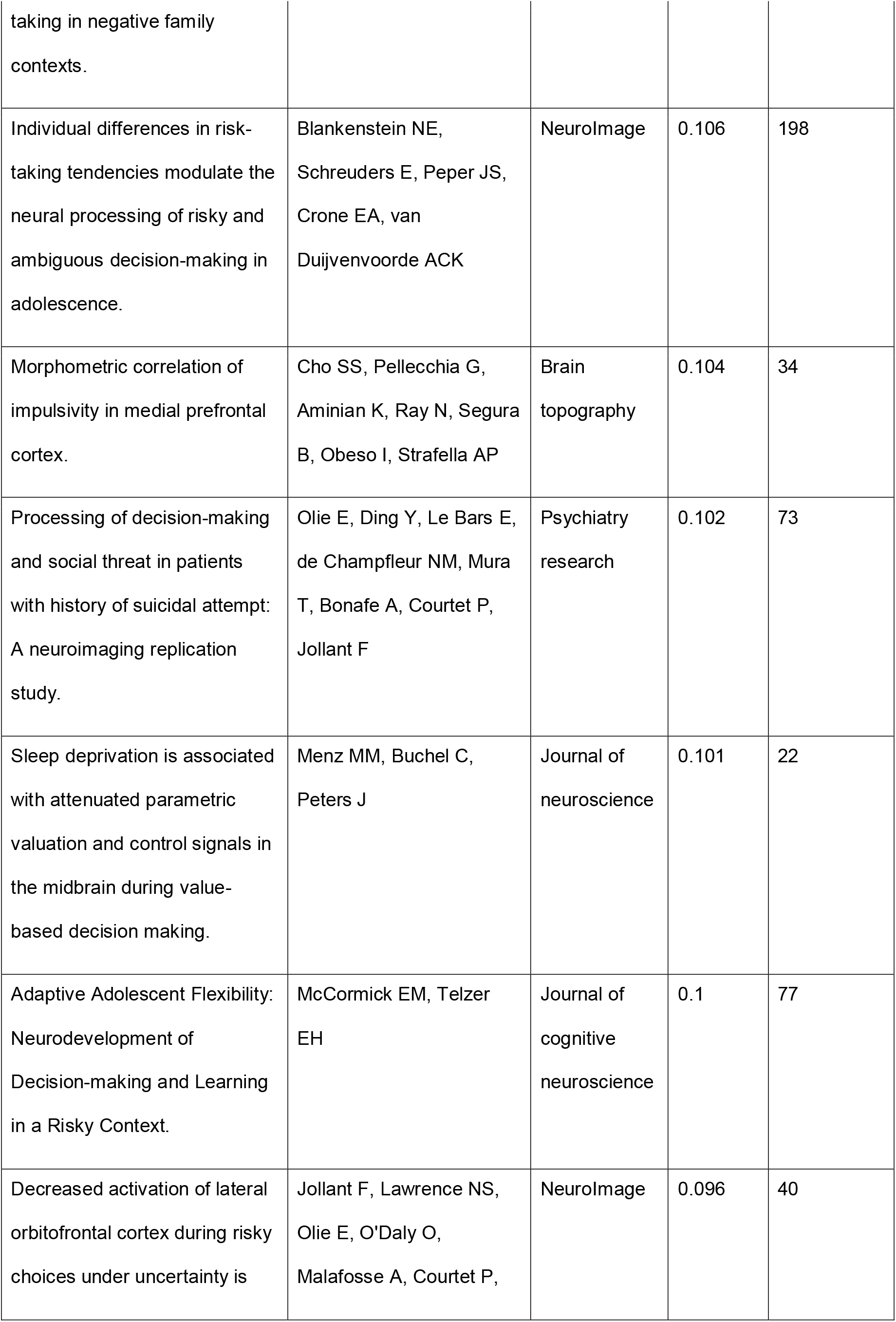
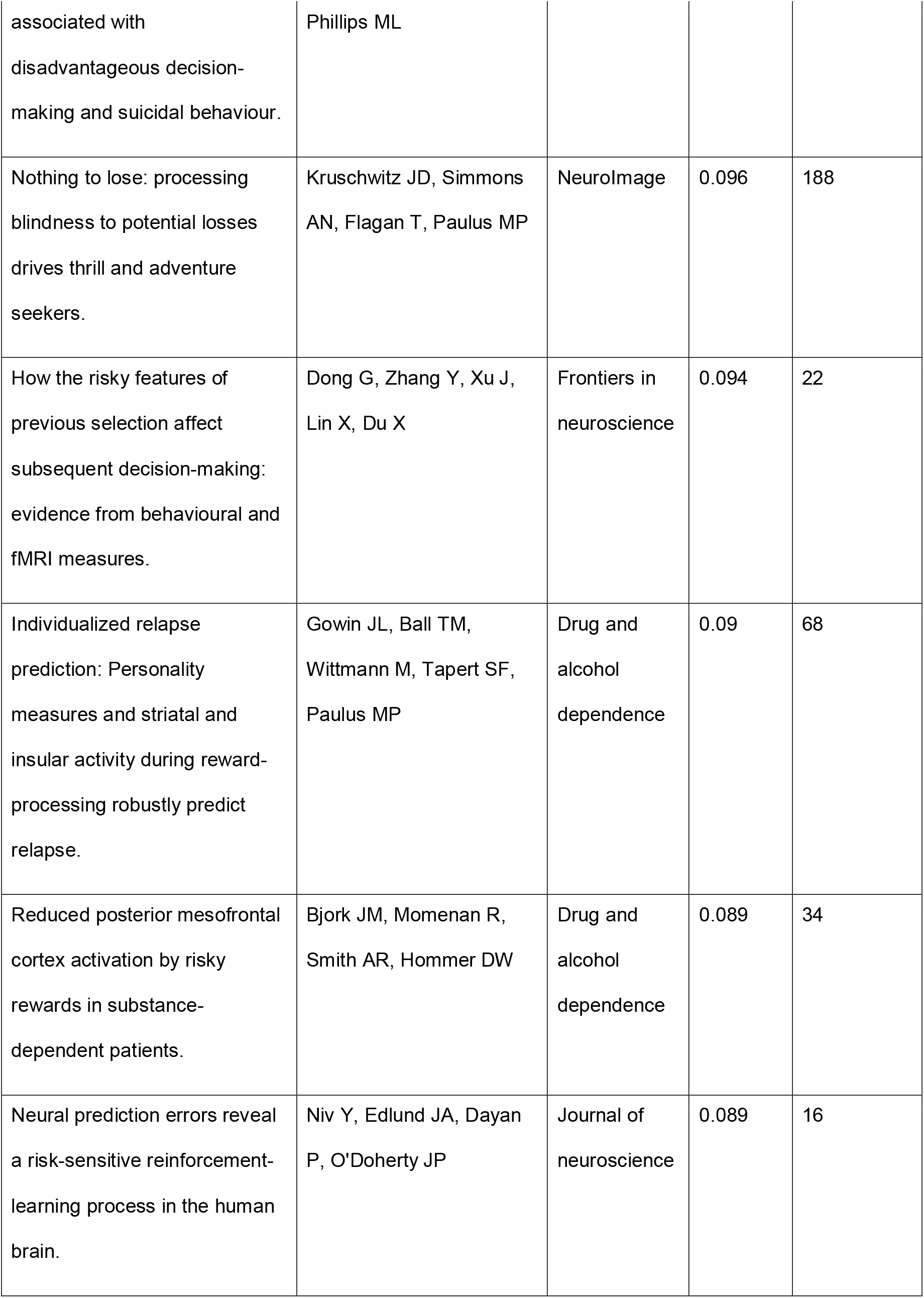
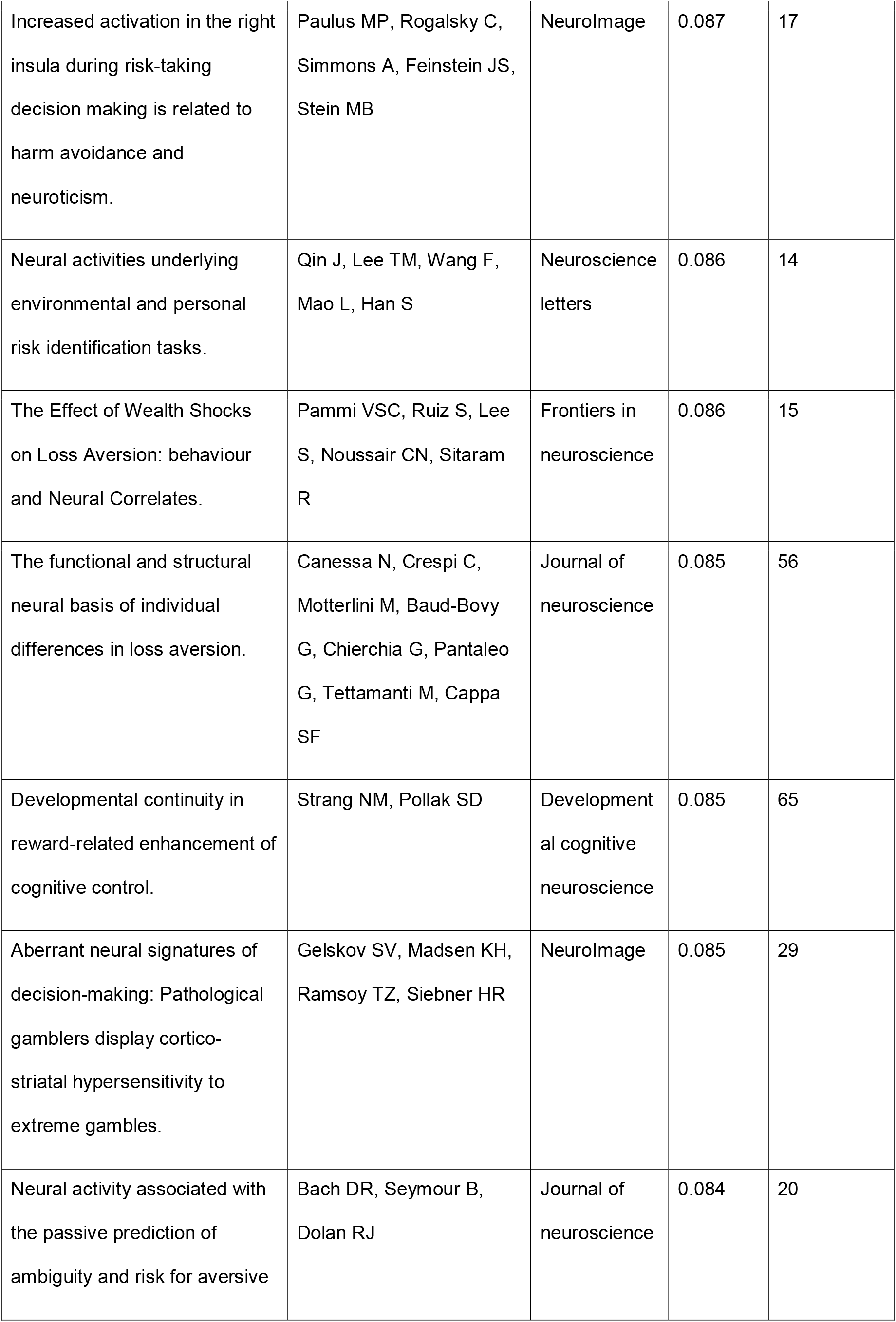
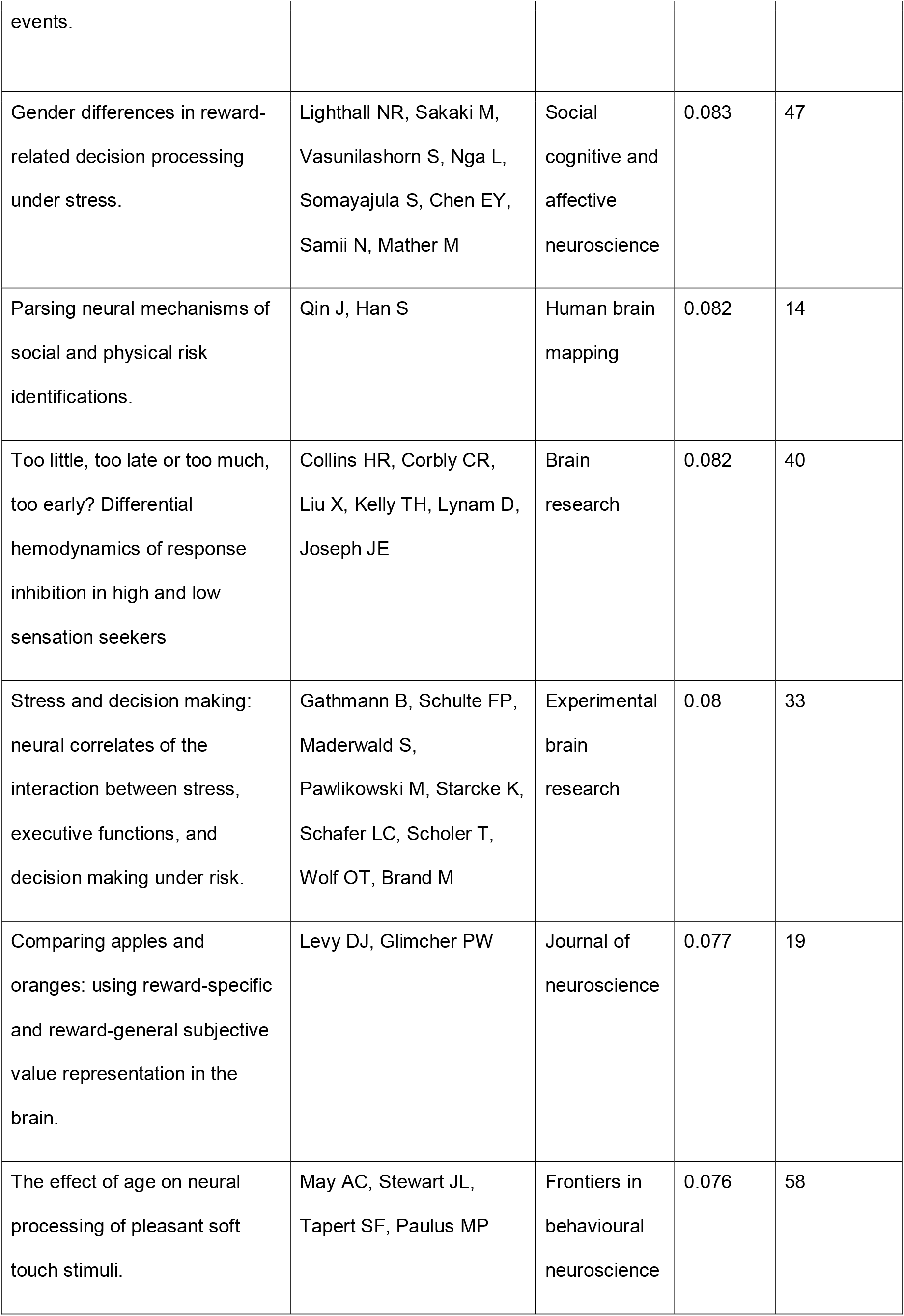
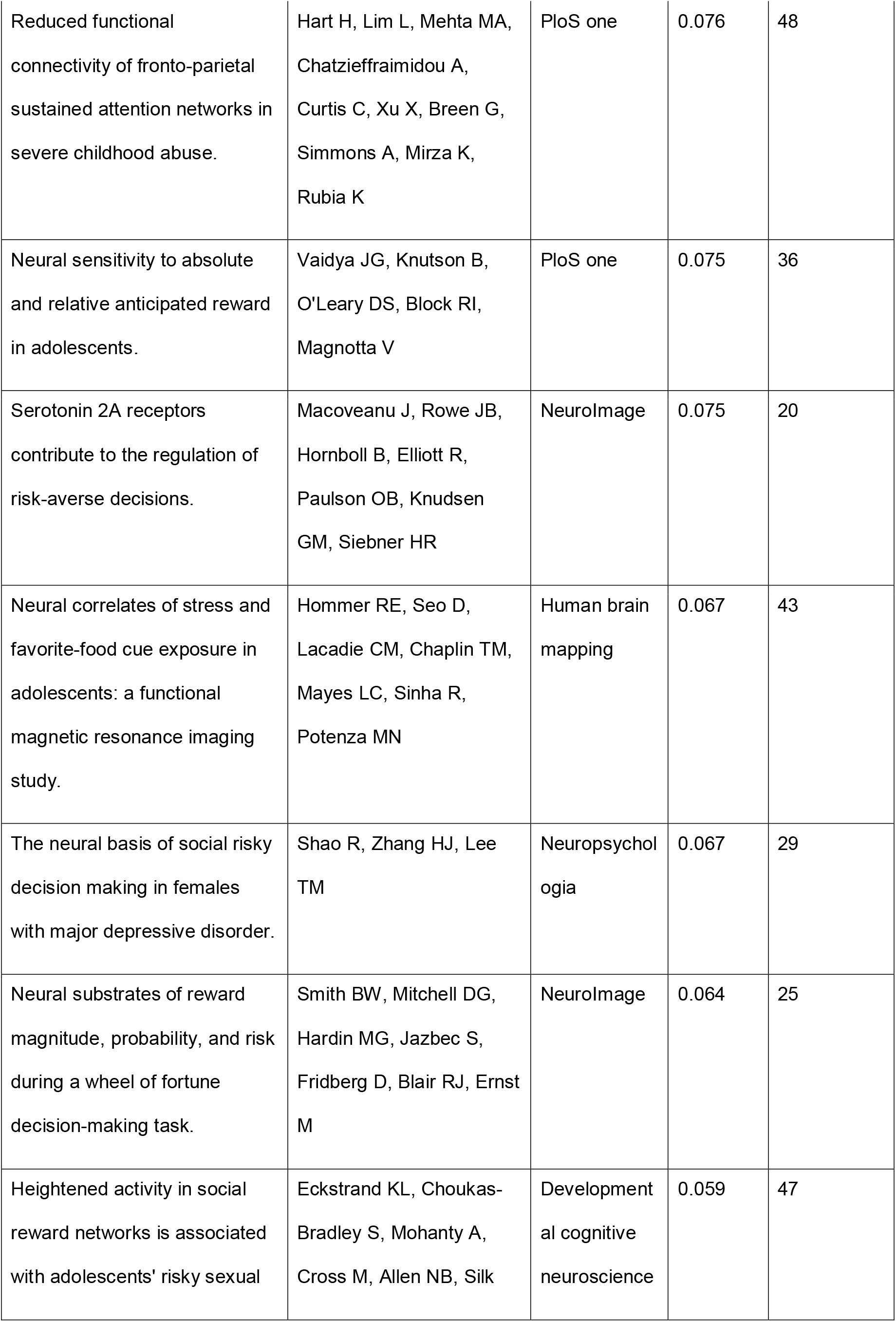
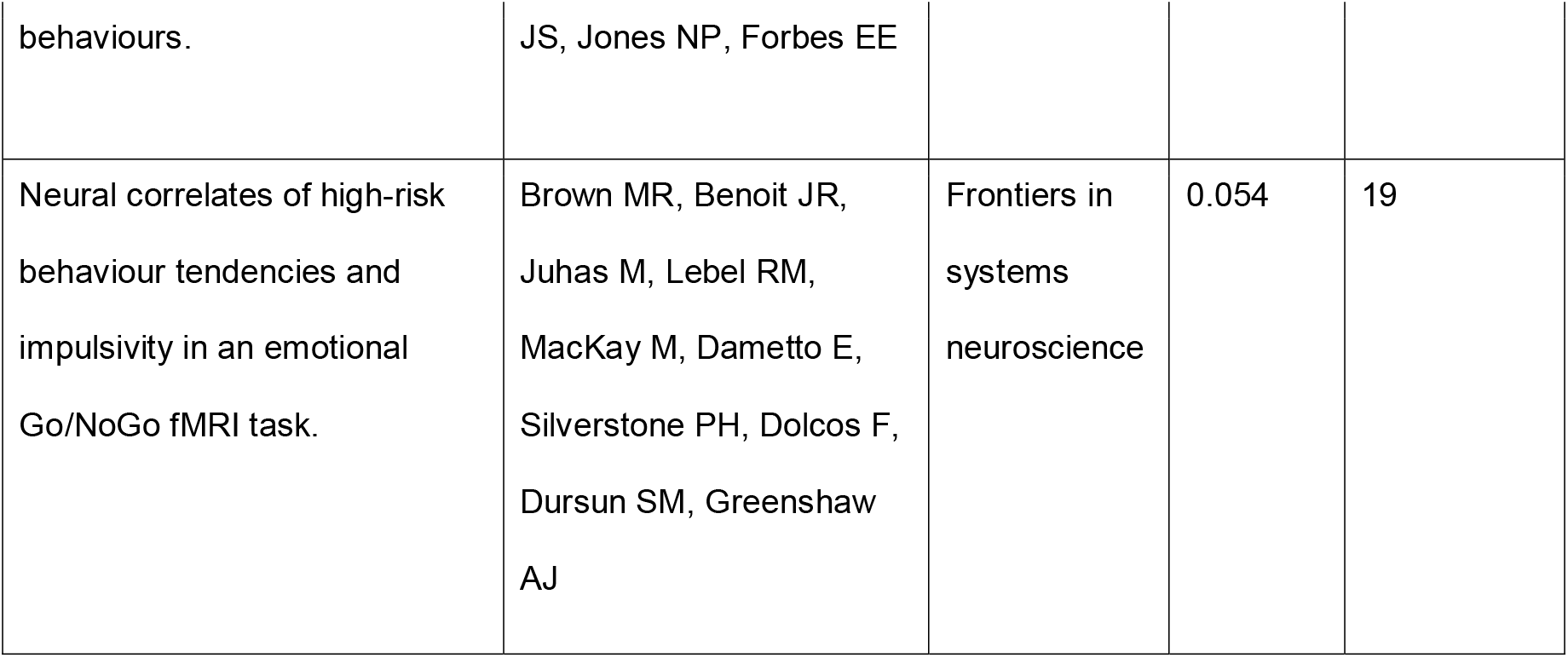
Summary of studies used for the meta-analysis of fMRI studies on risky behaviours (provided by Neurosynth). Neurosynth uses text mining techniques to search through published articles for certain keywords (here: ‘risky’) and then quantifies how important the keyword is in any particular published article, relative to all other searched articles. Specifically, Neurosynth uses a metric (i.e. ‘loading’) to quantify how often the key word (here: ‘risky’) was used in the respective article relative to all other articles in this meta-analysis. Its value ranges from 0 to 1 and increases proportionally with the number of times a word appears in the respective published article. Neurosynth typically uses a cutoff of .05 for articles to be included in the meta-analysis. For further information see ref 31.

## Footnotes

1 Self-reports of the number of sexual partners have been implicated in risky behaviours related to alcohol abuse (i.e., binge drinking) and unprotected sex, specifically in young adults^2^, irrespective of gender or sexual orientation.

## References

1. Knight, F. H. Risk, Uncertainty and Profit. (Houghton-Mifflin, 1921).

2. Eckel, C. C. & Füllbrunn, S. C. Thar SHE Blows? Gender, Competition, and Bubbles in Experimental Asset Markets. American Economic Review vol. 105 906–920 (2015).

3. Centers for Disease Control and Prevention. Incidence, Prevalence, and Cost of Sexually Transmitted Infections in the United States. https://www.cdc.gov/std/stats/sti-estimates-fact-sheet-feb-2013.pdf (2013).

4. Sacks, J. J., Gonzales, K. R., Bouchery, E. E., Tomedi, L. E. & Brewer, R. D. 2010 National and State Costs of Excessive Alcohol Consumption. Am. J. Prev. Med. 49, e73–e79 (2015).

5. Blincoe, L., Miller, T. R., Zaloshnja, E. & Lawrence, B. A. The economic and societal impact of motor vehicle crashes, 2010 (Revised). https://trid.trb.org/view/1311862 (2015).

6. U.S. Department of Health and Human Services. The Health Consequences of Smoking: 50 Years of Progress. A Report of the Surgeon General. Atlanta, GA: U.S. Department of Health and Human Services, Centers for Disease Control and Prevention, National Center for Chronic Disease Prevention and Health Promotion, Office on Smoking and Health (2014).

7. Karlsson Linnér, R. et al. Genome-wide association analyses of risk tolerance and risky behaviors in over 1 million individuals identify hundreds of loci and shared genetic influences. Nat. Genet. 51, 245–257 (2019).

8. Elliott, L. T. et al. Genome-wide association studies of brain imaging phenotypes in UK Biobank. Nature 562, 210–216 (2018).

9. Thompson, P. M. et al. Genetic influences on brain structure. Nat. Neurosci. 4, 1253–1258 (2001).

10. Grubb, M. A., Tymula, A., Gilaie-Dotan, S., Glimcher, P. W. & Levy, I. Neuroanatomy accounts for age-related changes in risk preferences. Nat. Commun. 7, 13822 (2016).

11. Jung, W. H., Lee, S., Lerman, C. & Kable, J. W. Amygdala Functional and Structural Connectivity Predicts Individual Risk Tolerance. Neuron 98, 394–404.e4 (2018).

12. Nasiriavanaki, Z. et al. Prediction of individual differences in risky behavior in young adults via variations in local brain structure. Front. Neurosci. 9, 359 (2015).

13. Button, K. S. et al. Power failure: why small sample size undermines the reliability of neuroscience. Nat. Rev. Neurosci. 14, 365–376 (2013).

14. Marek, S. et al. Towards Reproducible Brain-Wide Association Studies. bioRxiv 2020.08.21.257758 (2020).

15. Dohmen, T. et al. INDIVIDUAL RISK ATTITUDES: MEASUREMENT, DETERMINANTS, AND BEHAVIORAL CONSEQUENCES. Journal of the European Economic Association vol. 9 522–550 (2011).

16. Cardon, L. R. & Palmer, L. J. Population stratification and spurious allelic association. Lancet 361, 598–604 (2003).

17. Nave, G., Jung, W. H., Karlsson Linnér, R., Kable, J. W. & Koellinger, P. D. Are Bigger Brains Smarter? Evidence From a Large-Scale Preregistered Study. Psychol. Sci. 956797618808470 (2018).

18. Romer, D. Adolescent risk taking, impulsivity, and brain development: implications for prevention. Dev. Psychobiol. 52, 263–276 (2010).

19. Miller, K. L. et al. Multimodal population brain imaging in the UK Biobank prospective epidemiological study. Nature Neuroscience vol. 19 1523–1536 (2016).

20. Sudlow, C. et al. UK biobank: an open access resource for identifying the causes of a wide range of complex diseases of middle and old age. PLoS Med. 12, e1001779 (2015).

21. Harper, C. The neurotoxicity of alcohol. Hum. Exp. Toxicol. 26, 251–257 (2007).

22. Daviet, R., Aydogan, G., Jagannathan, K. & Spilka, N. Multimodal brain imaging study of 19,825 participants reveals adverse effects of moderate drinking. bioRxiv (2020).

23. Kranzler, H. R. et al. Topiramate Treatment for Heavy Drinkers: Moderation by a GRIK1Polymorphism. American Journal of Psychiatry vol. 171 445–452 (2014).

24. Ashburner, J. & Friston, K. J. Voxel-Based Morphometry—The Methods. Neuroimage 11, 805–821 (2000).

25. Botvinik-Nezer, R. et al. Variability in the analysis of a single neuroimaging dataset by many teams. Nature 1–7 (2020).

26. Alfaro-Almagro, F. et al. Image processing and Quality Control for the first 10,000 brain imaging datasets from UK Biobank. Neuroimage 166, 400–424 (2018).

27. Bycroft, C. et al. The UK Biobank resource with deep phenotyping and genomic data. Nature 562, 203–209 (2018).

28. Hill, W. D. et al. Molecular Genetic Contributions to Social Deprivation and Household Income in UK Biobank. Curr. Biol. 26, 3083–3089 (2016).

29. Masouleh, S. K., Eickhoff, S. B., Hoffstaedter, F., Genon, S. & Alzheimer’s Disease Neuroimaging Initiative. Empirical examination of the replicability of associations between brain structure and psychological variables. eLife vol. 8 (2019).

30. Yarkoni, T., Poldrack, R. A., Nichols, T. E., Van Essen, D. C. & Wager, T. D. Large-scale automated synthesis of human functional neuroimaging data. Nat. Methods 8, 665–670 (2011).

31. Mohr, P. N. C., Biele, G. & Heekeren, H. R. Neural processing of risk. J. Neurosci. 30, 6613–6619 (2010).

32. Wu, C. C., Sacchet, M. D. & Knutson, B. Toward an affective neuroscience account of financial risk taking. Front. Neurosci. 6, 159 (2012).

33. Schonberg, T., Fox, C. R. & Poldrack, R. A. Mind the gap: bridging economic and naturalistic risk-taking with cognitive neuroscience. Trends Cogn. Sci. 15, 11–19 (2011).

34. Kable, J. W. & Levy, I. Neural markers of individual differences in decision-making. Curr Opin Behav Sci 5, 100–107 (2015).

35. Chang, L. J., Yarkoni, T., Khaw, M. W. & Sanfey, A. G. Decoding the role of the insula in human cognition: functional parcellation and large-scale reverse inference. Cereb. Cortex 23, 739–749 (2013).

36. Evans, B. E., Greaves-Lord, K., Euser, A. S., Franken, I. H. A. & Huizink, A. C. The relation between hypothalamic-pituitary-adrenal (HPA) axis activity and age of onset of alcohol use. Addiction vol. 107 312–322 (2012).

37. Kreek, M. J., Nielsen, D. A., Butelman, E. R. & LaForge, K. S. Genetic influences on impulsivity, risk taking, stress responsivity and vulnerability to drug abuse and addiction. Nat. Neurosci. 8, 1450–1457 (2005).

38. O’Doherty, J. et al. Dissociable roles of ventral and dorsal striatum in instrumental conditioning. Science 304, 452–454 (2004).

39. Dosenbach, N. U. F., Fair, D. A., Cohen, A. L., Schlaggar, B. L. & Petersen, S. E. A dual-networks architecture of top-down control. Trends Cogn. Sci. 12, 99–105 (2008).

40. Ivanov, I., Schulz, K. P., London, E. D. & Newcorn, J. H. Inhibitory control deficits in childhood and risk for substance use disorders: a review. Am. J. Drug Alcohol Abuse 34, 239–258 (2008).

41. Heilman, R. M., Crişan, L. G., Houser, D., Miclea, M. & Miu, A. C. Emotion regulation and decision making under risk and uncertainty. Emotion 10, 257–265 (2010).

42. Tobler, P. N., Christopoulos, G. I., O’Doherty, J. P., Dolan, R. J. & Schultz, W. Riskdependent reward value signal in human prefrontal cortex. Proc. Natl. Acad. Sci. U. S. A. 106, 7185–7190 (2009).

43. Huizink, A. C., Ferdinand, R. F., Ormel, J. & Verhulst, F. C. Hypothalamic-pituitary-adrenal axis activity and early onset of cannabis use. Addiction 101, 1581–1588 (2006).

44. Margittai, Z. et al. Combined Effects of Glucocorticoid and Noradrenergic Activity on Loss Aversion. Neuropsychopharmacology 43, 334–341 (2018).

45. Grotzinger, A. D. et al. Hair and salivary testosterone, hair cortisol, and externalizing behaviors in adolescents. Psychol. Sci. 29, 688–699 (2018).

46. Buckner, R. L. The cerebellum and cognitive function: 25 years of insight from anatomy and neuroimaging. Neuron 80, 807–815 (2013).

47. Mechelli, A., Price, C., Friston, K. & Ashburner, J. Voxel-Based Morphometry of the Human Brain: Methods and Applications. Current Medical Imaging Reviews vol. 1 105–113 (2005).

48. Knoch, D. et al. Disruption of right prefrontal cortex by low-frequency repetitive transcranial magnetic stimulation induces risk-taking behavior. J. Neurosci. 26, 6469–6472 (2006).

49. Allen Institute for Brain Science. BrainSpan atlas of the developing human brain. http://www.brainspan.org/ (2015).

50. Loh, P.-R. et al. Efficient Bayesian mixed-model analysis increases association power in large cohorts. Nat. Genet. 47, 284–290 (2015).

51. Camerer, C. F. et al. Evaluating the replicability of social science experiments in Nature and Science between 2010 and 2015. Nat Hum Behav 2, 637–644 (2018).

52. Dickie, D. A. et al. Permutation and parametric tests for effect sizes in voxel-based morphometry of gray matter volume in brain structural MRI. Magnetic Resonance Imaging vol. 33 1299–1305 (2015).

## Supplementary References

1. Karlsson Linnér, R. et al. Genome-wide association analyses of risk tolerance and risky behaviors in over 1 million individuals identify hundreds of loci and shared genetic influences. Nat. Genet. 51, 245–257 (2019).

2. Desiderato, L. L. & Crawford, H. J. Risky sexual behavior in college students: Relationships between number of sexual partners, disclosure of previous risky behavior, and alcohol use. Journal of Youth and Adolescence vol. 24 55–68 (1995).

3. Mata, R., Frey, R., Richter, D., Schupp, J. & Hertwig, R. Risk Preference: A View from Psychology. J. Econ. Perspect. 32, 155–172 (2018).

4. Mata, R., Josef, A. K. & Hertwig, R. Propensity for Risk Taking Across the Life Span and Around the Globe. Psychol. Sci. 27, 231–243 (2016).

5. Lönnqvist, J.-E., Verkasalo, M., Walkowitz, G. & Wichardt, P. C. Measuring individual risk attitudes in the lab: Task or ask? An empirical comparison. Journal of Economic Behavior & Organization vol. 119 254–266 (2015).

6. Okbay, A. et al. Genome-wide association study identifies 74 loci associated with educational attainment. Nature 533, 539–542 (2016).

7. Croson, R. & Gneezy, U. Gender Differences in Preferences. J. Econ. Lit. 47, 448–474 (2009).

8. Dohmen, T. et al. INDIVIDUAL RISK ATTITUDES: MEASUREMENT, DETERMINANTS, AND BEHAVIORAL CONSEQUENCES. Journal of the European Economic Association vol. 9 522–550 (2011).

9. Cardon, L. R. & Palmer, L. J. Population stratification and spurious allelic association. Lancet 361, 598–604 (2003).

10. Alfaro-Almagro, F. et al. Image processing and Quality Control for the first 10,000 brain imaging datasets from UK Biobank. Neuroimage 166, 400–424 (2018).

11. Sallet, J. et al. The organization of dorsal frontal cortex in humans and macaques. J. Neurosci. 33, 12255–12274 (2013).

12. Neubert, F.-X., Mars, R. B., Sallet, J. & Rushworth, M. F. S. Connectivity reveals relationship of brain areas for reward-guided learning and decision making in human and monkey frontal cortex. Proc. Natl. Acad. Sci. U. S. A. 112, E2695–704 (2015).

13. Boorman, E. D., Behrens, T. E. J., Woolrich, M. W. & Rushworth, M. F. S. How Green Is the Grass on the Other Side? Frontopolar Cortex and the Evidence in Favor of Alternative Courses of Action. Neuron vol. 62 733–743 (2009).

14. Lim, S.-L., -L. Lim, S., O’Doherty, J. P. & Rangel, A. The Decision Value Computations in the vmPFC and Striatum Use a Relative Value Code That is Guided by Visual Attention. Journal of Neuroscience vol. 31 13214–13223 (2011).

15. Barron, H. C., Dolan, R. J. & Behrens, T. E. J. Online evaluation of novel choices by simultaneous representation of multiple memories. Nature Neuroscience vol. 16 1492–1498 (2013).

16. Kable, J. W. & Glimcher, P. W. The neural correlates of subjective value during intertemporal choice. Nat. Neurosci. 10, 1625 (2007).

17. Ongur, D. The Organization of Networks within the Orbital and Medial Prefrontal Cortex of Rats, Monkeys and Humans. Cerebral Cortex vol. 10 206–219 (2000).

18. Pauli, W. M., Nili, A. N. & Tyszka, J. M. A high-resolution probabilistic in vivo atlas of human subcortical brain nuclei. Sci Data 5, 180063 (2018).

19. Poppenk, J., Evensmoen, H. R., Moscovitch, M. & Nadel, L. Long-axis specialization of the human hippocampus. Trends Cogn. Sci. 17, 230–240 (2013).

20. Chang, L. J., Yarkoni, T., Khaw, M. W. & Sanfey, A. G. Decoding the role of the insula in human cognition: functional parcellation and large-scale reverse inference. Cereb. Cortex 23, 739–749 (2013).

21. Manichaikul, A. et al. Robust relationship inference in genome-wide association studies. Bioinformatics 26, 2867–2873 (2010).

22. Loh, P.-R. et al. Efficient Bayesian mixed-model analysis increases association power in large cohorts. Nat. Genet. 47, 284–290 (2015).

23. Loh, P.-R., Kichaev, G., Gazal, S., Schoech, A. P. & Price, A. L. Mixed-model association for biobank-scale datasets. Nat. Genet. 50, 906–908 (2018).

24. International HapMap 3 Consortium et al. Integrating common and rare genetic variation in diverse human populations. Nature 467, 52–58 (2010).

25. Bulik-Sullivan, B. K. et al. An atlas of genetic correlations across human diseases and traits. Nat. Genet. 47, 1236–1241 (2015).

26. Watanabe, K. et al. A global overview of pleiotropy and genetic architecture in complex traits. Nat. Genet. (2019) doi:10.1038/s41588-019-0481-0.

27. Meddens, S. F. W. et al. Genomic analysis of diet composition finds novel loci and associations with health and lifestyle. doi:10.1101/383406.

28. Owens, M. M. et al. Neuroanatomical foundations of delayed reward discounting decision making. NeuroImage vol. 161 261–270 (2017).

29. Jung, W. H., Lee, S., Lerman, C. & Kable, J. W. Amygdala Functional and Structural Connectivity Predicts Individual Risk Tolerance. Neuron vol. 98 394–404.e4 (2018).

30. Hagler, D. J., Jr et al. Image processing and analysis methods for the Adolescent Brain Cognitive Development Study. Neuroimage 202, 116091 (2019).

31. Yarkoni, T., Poldrack, R. A., Nichols, T. E., Van Essen, D. C. & Wager, T. D. Large-scale automated synthesis of human functional neuroimaging data. Nat. Methods 8, 665–670 (2011).

